# Functional role of myosin-binding protein H in thick filaments of developing vertebrate fast-twitch skeletal muscle

**DOI:** 10.1101/2024.05.10.593199

**Authors:** Andrew F. Mead, Neil B. Wood, Shane R. Nelson, Bradley M. Palmer, Lin Yang, Samantha Beck Previs, Angela Ploysangngam, Guy G. Kennedy, Jennifer F. McAdow, Sarah M. Tremble, Marilyn J. Cipolla, Alicia M. Ebert, Aaron N. Johnson, Christina A. Gurnett, Michael J. Previs, David M. Warshaw

## Abstract

Myosin-binding protein H (MyBP-H) is a component of the vertebrate skeletal muscle sarcomere with sequence and domain homology to myosin-binding protein C (MyBP-C). Whereas skeletal muscle isoforms of MyBP-C (fMyBP-C, sMyBP-C) modulate muscle contractility via interactions with actin thin filaments and myosin motors within the muscle sarcomere “C-zone,” MyBP-H has no known function. This is in part due to MyBP-H having limited expression in adult fast-twitch muscle and no known involvement in muscle disease. Quantitative proteomics reported here reveal MyBP-H is highly expressed in prenatal rat fast-twitch muscles and larval zebrafish, suggesting a conserved role in muscle development, and promoting studies to define its function. We take advantage of the genetic control of the zebrafish model and a combination of structural, functional, and biophysical techniques to interrogate the role of MyBP-H. Transgenic, FLAG-tagged MyBP-H or fMyBP-C both localize to the C-zones in larval myofibers, whereas genetic depletion of endogenous MyBP-H or fMyBP-C leads to increased accumulation of the other, suggesting competition for C-zone binding sites. Does MyBP-H modulate contractility from the C-zone? Globular domains critical to MyBP-C’s modulatory functions are absent from MyBP-H, suggesting MyBP-H may be functionally silent. However, our results suggest an active role. Small angle x-ray diffraction of intact larval tails revealed MyBP-H contributes to the compression of the myofilament lattice accompanying stretch or contraction, while *in vitro* motility experiments indicate MyBP-H shares MyBP-C’s capacity as a molecular “brake”. These results provide new insights and raise questions about the role of the C-zone during muscle development.

## Introduction

The “C-zone” is a defined region of the vertebrate muscle sarcomere (Fig. 1A), characterized primarily by the presence of myosin-binding protein C (MyBP-C), which binds to the myosin thick filament backbone (1, 2). MyBP-C is a key regulator of muscle contractility, whose importance is emphasized by its involvement in human muscular diseases including hypertrophic cardiomyopathy and distal arthrogryposis (3–7). Multiple, functionally distinct isoforms of MyBP-C are encoded by three paralogous genes: *MYBPC1* (sMyBP-C “*slow skeletal*”), *MYBPC2* (fMyBP-C “*fast skeletal*”), and *MYBPC3* (cMyBP-C “*cardiac*”) (Fig. 1B) (8–11). These genes diverged and specialized early in vertebrate evolution, establishing C-zone-based mechano-regulation of muscle contractility as a ubiquitous feature of vertebrate striated muscle (8). While MyBP-C has been studied extensively, it is not the only myosin thick filament-associated protein found in the C-zone. Two structurally related but shorter proteins, myosin-binding protein H-like (MyBP-HL) and myosin-binding protein H (MyBP-H), are expressed in adult mammalian atrial and fast-twitch skeletal muscle, respectively, and are encoded by separate genes (*MYBPHL*, *MYBPH*) (Fig. 1B) (1, 12–14), though the evolutionary relationships of these genes to the larger MyBP-C gene family have not been explored. Recent studies have implicated MyBP-HL in human cardiomyopathies (12, 15). Much less attention has been paid to its skeletal muscle counterpart MyBP-H as expression in adult muscle is low compared to MyBP-C, and no disease-causing mutations have been identified (16). The present study stems in large part from proteomics data reported here, which shows that prenatal rat fast-twitch limb muscle samples contain high levels of MyBP-H during the latter stages of muscle development prior to birth. Remarkably, we find a similar expression pattern in the rapidly developing myotomal muscles of five-day-old zebrafish larvae, indicating a possible novel and broadly conserved role for MyBP-H during muscle growth, and supporting use of zebrafish as a model system to study MyBP-H function.

**Figure 1.**
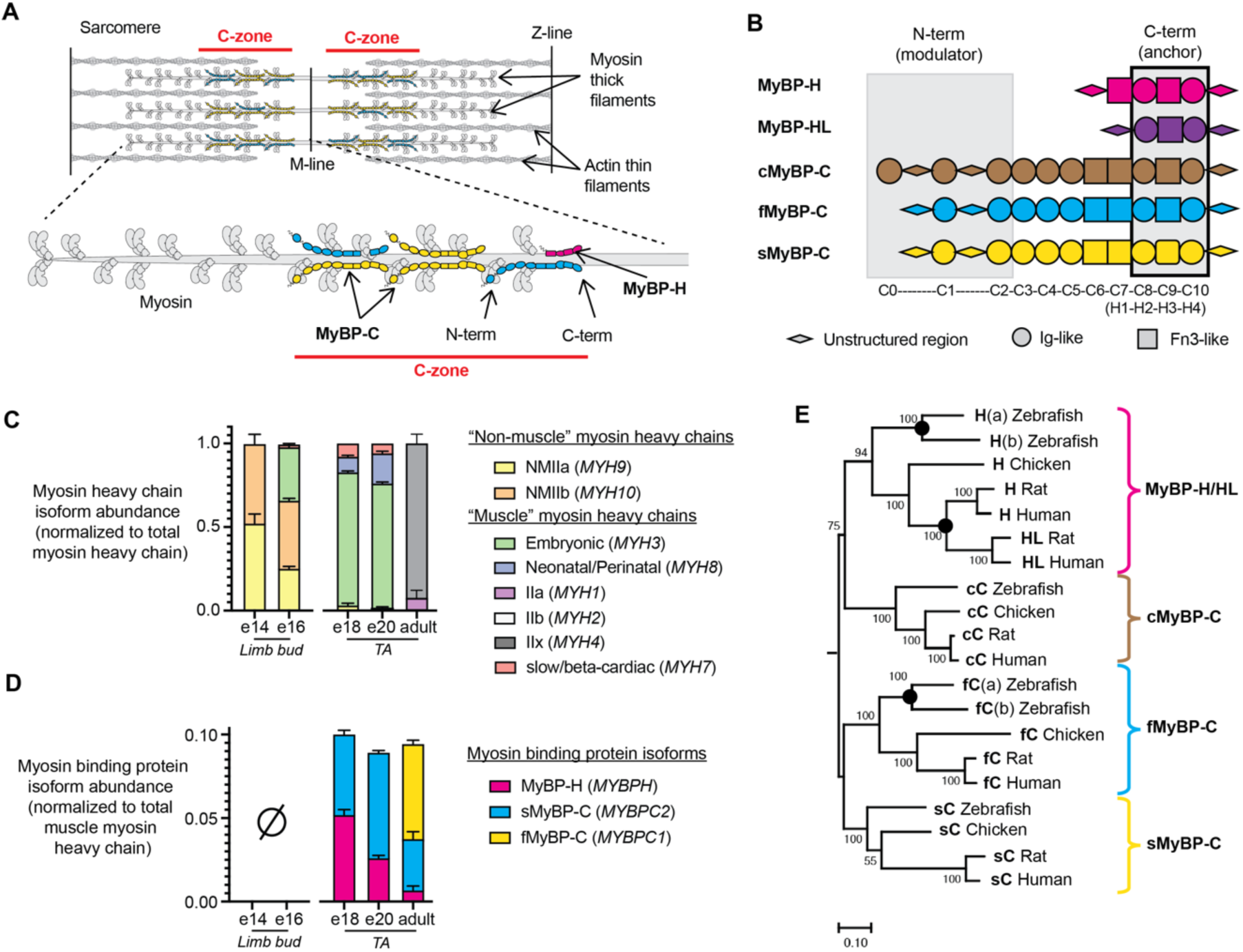
The sarcomere “C-zone” is home to the MyBP-C/H family of regulatory proteins. (A) A schematic of the vertebrate skeletal muscle sarcomere, consisting of interdigitating myosin thick- and actin thin-filaments showing the “C-zone” (red) and expanded view of half-thick filament illustrating a mixture of MyBP-C/H molecules within the C-zone with their N- and C-termini identified. (B) Domain structure and homology among MyBP-C and MyBP-H isoforms showing conservation of C-terminal, myosin thick filament anchoring domains. (C) Normalized abundance of individual myosin heavy chain isoforms (with gene names in parentheses) in embryonic hindlimb buds and prenatal and adult rat tibialis anterior (TA) muscle samples relative to total myosin heavy chain abundance at each timepoint. (D) Abundance of individual myosin binding protein (MyBP) isoforms in embryonic hindlimb bud (none detected) and prenatal and adult rat TA muscle samples, relative to total muscle myosin heavy chain. Data are presented as means ± 1 S.D.. (E) A midpoint-rooted Maximum Likelihood consensus tree of the MyBP family based on an alignment of 3 C-terminal globular domains (294 amino acids) conserved among all family members from four species. The percentage of trees, from 100 bootstrapped replicates, in which the associated sequences clustered together is shown next to the branches. Initial tree(s) for the heuristic search were obtained automatically by applying Neighbor-Join and BioNJ algorithms to a matrix of pairwise distances estimated using the JTT model, and then selecting the topology with superior log likelihood value. The tree is drawn to scale with branch lengths equal to the number of substitutions per site. Sequences cluster into 4 distinct clades, corresponding to the four paralogous MyBP gene family members. Orthologous sequences from each species are represented within each clade, indicating divergence and specialization early in vertebrate evolution. The grouping of mammalian MyBP-H and MyBP-HL, as well as zebrafish fMyBP-C (a/b) and MyBP-H (a/b) is indicative of more recent, lineage-specific gene duplication events (nodes with black dots).

MyBP-H was first identified as an impurity in contractile protein isolates of rabbit skeletal muscle (16), but later confirmed to be a thick filament associated protein that localized to the C-zone (1, 13, 14) (Fig. 1A). MyBP-H is a ∼59 kDa protein consisting of 4 globular domains; 2 immunoglobulin-like (Ig-like) and 2 fibronectin-III-like (Fn3-like) domains connected in series by flexible linkers (Fig. 1B). These 4 globular domains share significant sequence and structural homology to the 4 C-terminal Ig- and Fn3-like domains of MyBP-C (17), which serve to anchor both MyBP-H and MyBP-C to the myosin thick-filament backbone. In immuno-electron microscopic images from adult, rabbit psoas, MyBP-H and MyBP-C appear as stripes in the C-zone that are spaced 43 nm apart, matching the period of the myosin helical repeat within the thick filament. However, MyBP-H is restricted to a single stripe closest to the M-line, whereas MyBP-C is localized to the subsequent 8 distal stripes, depending on isoform and fiber type (1).

Based on our current understanding of the relationship between MyBP-C structure and function, a C-zone rich in MyBP-H may have important consequences for sarcomere contractility. In stark contrast to skeletal MyBP-C, MyBP-H lacks the 6 N-terminal Ig- and Fn3-like domains that extend away from the thick filament (18) and are believed to interact with the myosin head and/or the actin-thin filaments to modulate contractility in an isoform-dependent manner (19). Specifically, these interactions have been shown *in vitro* to sensitize the actin-thin filament to calcium ions and to exert a load that resists thin filament sliding (19), a “braking” effect also observed in single muscle fibers (20). In place of MyBP-C’s N-terminal modulatory domains, MyBP-H has a long (78-134 amino acid), proline-alanine rich, unstructured domain that is somewhat similar to the very N-terminus of the skeletal muscle MyBP-Cs (9). With MyBP-H devoid of the N-terminal Ig-/Fn3-like domains that are critical to the modulatory capacity of MyBP-C, one may hypothesize that MyBP-H serves a structural role in the thick filament but acts as a functional null. Alternatively, MyBP-H may be capable of affecting contractility directly, either via its pro/ala rich N terminus, or through direct interactions between its globular domains and myosin heads within the thick filament.

In the present study we explored the evolutionary relationship of MyBP-H to the MyBP-C gene family and employ a versatile zebrafish larval muscle model to investigate the role of MyBP-H in relation to MyBP-C within the skeletal muscle C-zone. Within days of fertilization, zebrafish larvae exhibit swimming behavior that is powered by myotomal tail muscles comprised of ∼95% fast-twitch skeletal muscle fibers (21, 22). We and others have taken advantage of the transparency of the larval tail, and the parallel organization of the muscle fibers within, to characterize the mechanical performance of myotomal muscles at the sarcomere level (22–25). Critically, our phylogenetic analysis shows that zebrafish possess orthologs for all mammalian skeletal muscle MyBP-C and MyBP-H genes. In addition to being amenable to genetic manipulation, the ability to extract physiological data during muscle development gives zebrafish a distinct advantage over rodent models. Therefore, we used proteomic and biophysical approaches to define how genetically altering the expression level and localization of MyBP-H and MyBP-C in the fast-twitch skeletal muscle sarcomere C-zone of zebrafish larvae and adults provide insights to a physiological role for MyBP-H as a functional element that impacts the mechanical interaction between thick and thin filaments in the sarcomere.

## Results

### MyBP-H is highly enriched in developing fast-twitch mammalian skeletal muscle

In mature mammalian striated muscles, MyBP-C isoforms consistently sum to a molar ratio of approximately 0.1 relative to the summation of sarcomeric myosin heavy chain isoforms (19, 26), a relationship assumed to depend on the existence of limited number of binding sites within the thick filament C-zones. To confirm this molar ratio in an adult mammalian fast-twitch skeletal muscle and to determine whether it is conserved during prenatal muscle development, we identified the myosin binding proteins (MyBPs) and myosin heavy chain isoforms in adult and prenatal rat hindlimb tissue using liquid chromatography mass spectrometry (LCMS) and quantified their abundances using label-free analysis methods (see Methods). The relative distribution of myosin heavy chain isoforms within each muscle sample was determined from the average abundance of the top peptides unique to each myosin heavy chain isoform divided by the summed abundance of top peptides unique to each myosin heavy chain isoform (Fig. 1C, Table S1). The molar abundance of MyBP isoforms relative to myosin heavy chain was determined from the average abundance of the top peptides unique to each MyBP isoform divided by the summed abundance of top peptides shared between all muscle myosin heavy chain isoforms (Fig 1D, Table S2).

In adult tibialis anterior (TA) muscles, unique peptides were detected from fast-type myosin heavy chain isoforms IIa (*MYH2*), IIb (*MYH4*), and IIx (*MYH1*) (Table S1), with those from myosin heavy chain IIx being the most abundant (Fig. 1C). Unique peptides were also detected from MyBP-H, sMyBP-C, and fMyBP-C (Table S2), with those from the MyBP-C isoforms being the most abundant (Fig. 1D) as expected (19). The molar ratio of peptides from the MyBP-C isoforms to those from myosin heavy chain was between 0.088 and 0.095 (Fig. 1D), in agreement with the ∼0.1 molar ratio reported in other adult muscle types (19, 26). The low abundance of MyBP-H peptides is also consistent with its more restricted location within the C-zone, as predicted by immuno-transmission electron microscopy (1).

Is the molar ratio of MyBP to myosin heavy chain constant throughout prenatal muscle development, and is MyBP-H expressed to a greater extent than in the adult? To address this, we identified and quantified the expression of MyBP and myosin heavy chain isoforms in prenatal hindlimbs at embryonic days 14, 16, 18, and 20 (e14, e16, e18, e20, respectively). At e14 and e16, individual hindlimb muscles were not easily identifiable and therefore, whole hindlimb buds were removed, digested with trypsin and analyzed by LCMS.

At e14, peptides from the non-muscle myosin heavy chain isoforms NMIIA (*MYH9*) and NMIIB (*MYH10*) predominated (Fig. 1C, Table S1) and no MyBP peptides were detected (Fig. 1D, Table S2). The presence of only non-muscle myosin heavy chain isoforms and absence of MyBPs suggest the hindlimb muscles are predominately comprised of myoblasts at this developmental stage, as previously described (27). By e16, peptides from the embryonic (*MYH3*) and to a lesser extent slow/cardiac (*MYH7*), muscle myosin heavy chain isoforms appear (Fig. 1C), but MyBPs are not present (Fig 1D). The presence of the muscle myosin heavy chains is likely indicative of the formation of pre-myofibrils (28), while the absence of MyBPs may indicate that their incorporation into thick filaments takes place only after this key transition.

By e18, TA muscles could be identified, dissected, and processed for LCMS. The presence of the non-muscle myosin heavy chain isoforms was minimal, and the muscles were predominantly comprised of the embryonic (*MYH3*), neonatal (*MYH8*), and slow/cardiac (*MYH7*) muscle myosin isoforms (Fig. 1C, Table S1). Both the MyBP-H and sMyBP-C isoforms were expressed with a 0.104 molar ratio of MyBPs to myosin heavy chains, equivalent to that in the adult TA muscle (Fig. 1D, Table S2). These data were consistent with the muscles containing myofibrils. At e20, the distribution of myosin heavy chain isoforms was similar (Fig. 1C, Table S1), and the molar ratio of MyBPs to myosin heavy chains was 0.092, being comparable to that at e18 and in the adult TA muscle (Fig 1D, Table S2). Most importantly, MyBP-H made up a significant share of MyBP at both the e18 and e20 developmental time points, with peptide abundances relative to myosin heavy chain of 0.054 ± 0.003 and 0.026 ± 0.002, respectively. These levels were significantly higher than in adult TA muscle (0.007 ± 0.002, P<0.01) (Fig. 1D, Table S2), indicating that MyBP-H may occupy a larger share of the C-zone and thus serve an underappreciated role during these stages of development.

### MyBP-H is an early-diverging member of the MyBP-C family

Three MyBP-C gene paralogs *MYBPC1* (sMyBP-C protein), *MYBPC2* (fMyBP-C protein), and *MYBPC3* (cMyBP-C protein) diverged early in vertebrate evolution (8), and are conserved in all major vertebrate taxa. While amino acid sequence homology of MyBP-H and MyBP-HL to the C-terminal domains of MyBP-C has been noted (29), their evolutionary relationship to MyBP-C gene family is not well defined. Therefore, we acquired reference MyBP-C/H/HL sequences from representatives of three major vertebrate taxa: mammals (human, rat), avians (chicken), and fish (zebrafish) (File S1) and performed a phyologenetic analysis. Human, rat, and chicken genomes each possess a single ortholog of *MYBPH*, in addition to *MYBPC1, MYBPC2, MYBPC3.* Human and rat genomes possess *MYPBHL* genes, but none was identified in non-mammals. The zebrafish genome possesses four putative MyBP-C- and two putative MyBP-H-encoding genes, each annotated as an ortholog of either human *MYBPC1*, *MYBPC2*, *MYBPC3,* or *MYBPH*. *MYBPC2* and *MYBPH* are represented in the zebrafish genome by pairs of genes (*mybpc2a, mybpc2b; mybpha, mybphb*).

We first aligned the 19 full-length protein sequences (See *Methods*) (File S1). MyBP-H sequences aligned with the C-terminal region of MyBP-C, corresponding to globular domains C7-C10 (Fig. 1B, File S1), as shown previously (17). The same was true for the shorter MyBP-HL sequences, which aligned with MyBP-C domains C8-C10 (Fig. 1B, File S1). In order to remove the confounding effects of different protein sequence lengths and domain structures among the various MyBP-C and MyBP-H/HL sequences, we performed a second alignment of amino acid sequences corresponding to the 3 C-terminal globular domains shared by all isoforms, starting with a conserved proline at human sMyBP-C position 844 and ending with the conserved cysteine at human sMyBP-C position 1139 (Fig. 1B, black-outlined rectangle; File S1, green arrows). From this alignment we inferred phylogeny using the maximum likelihood method (see Methods) (Fig. 1E). In the resulting phenogram, MyBP-C sequences from all species grouped into three monophyletic clades corresponding to sMyBP-C, fMyBP-C, and cMyBP-C, as reported by (8), while the MyBP-H/HL sequences formed a fourth clade. Within each clade, tree topology matched the evolutionary relationships between species, as would be expected for orthologous sequences. The MyBP-H and fMyBP-C clades contained evidence of more recent, lineage-specific, gene-duplication events (Fig. 1E; nodes with black dots). In zebrafish, sequences from both pairs of genes (*mybpha, mybphb; mybpc2a, mybpc2b*) clustered together, indicating that, like many other zebrafish genes, they arose from a whole-genome duplication known to have taken place in ray finned fish (30). The tree also indicates MyBP-HL resulted from a relatively recent duplication of the MyBP-H gene in a common ancestor of humans and rats.

Together, these data indicate that genes encoding MyBP-H, sMyBP-C, fMyBP-C, and cMyBP-C are evolutionary paralogs, and arose from a common ancestral gene early in vertebrate evolution. Our analysis suggests that MyBP-H may have arisen from the duplication of an ancestral MyBP-C-like gene which subsequently lost its N-terminal globular domains. However, the close proximity of early branch points complicates determining the precise order in which the four clades evolved.

### MyBP-H binds across the full C-zone in developing zebrafish muscle

Since the MyBP-C/MyBP-H gene family was established early and persisted with few changes between the rat and zebrafish lineages, we took advantage of the versatility of the zebrafish as a model system to investigate the role of MyBP-H in the skeletal muscle C-zones. Zebrafish myotomal tail muscles develop rapidly and power a discreet set of locomotory maneuvers within days of fertilization, at which point sarcomere-level contractile function can be assessed using *ex vivo* intact muscle mechanics approaches (21, 22). To quantify MyBP composition in zebrafish larvae, we digested wildtype tails with trypsin at five days post-fertilization (5dpf) and performed label-free LCMS (see Methods). Notably, consistent results were achieved from single tails despite their small size (∼0.03 mg). Unique peptides from 8 myosin heavy chain genes were present in the samples, with peptide abundances from fast-type myosin isoforms accounting for ∼90%, and slow-type myosin isoforms ∼10% of the peptides (Table S3). It should be noted that there is a high degree of sequence similarity between myosin isoforms which limits the number of unique peptides and may impact the apparent absolute abundance of each isoform. However, the isoform abundances were similar to that expected from the relative cross-sectional areas of fast and slow twitch muscle in 5dpf tails, as reported previously and below (Table S3).

Unique peptides were found from zebrafish MyBP isoforms above the threshold for quantification. These were Mybphb, the gene product of *mybphb,* one of the two *MYBPH* orthologs, and Mybpc2b, the gene product of *mybpc2b,* one of the two *MYBPC2* orthologs, with the abundance of those peptides from Mybphb having a ∼20-fold greater molar abundance than those from Mybpc2b (Fig. 2A, Table S4). The summation of the abundance of the top peptides from Mybphb and Mybpc2b was used to determine the relative abundance of total MyBP protein to myosin heavy chain. The combined molar ratio of MyBP (Mybphb + Mybpc2b) to myosin heavy chain (0.082 ± 0.008) was slightly lower than prenatal rat TA at e18 (0.104 ± 0.001, P<0.01) but not e20 (0.092 ± 0.001, P>0.08), and within the range of that reported for mammalian muscle types (19, 26) (Fig. 1D, 2A). To determine whether high level of MyBP-H expression was unique to developing zebrafish muscle, as in the rat, or persisted through adulthood, we dissected fast myotomal swimming muscle tissue samples (∼3 mg) from adult (6-12 month-old) wildtype zebrafish and performed LCMS analyses. The peptides which were unique to specific myosin heavy chain isoforms were almost exclusively from fast myosin heavy chain genes (Table S3) in agreement with previous reports (31). Interestingly, adult zebrafish fast-twitch myotomal muscle also contained a high abundance of Mybphb peptides, though by this stage Mybpc2b accounted for a larger share of total MyBP (Mybpc2b: adult: 28.6 ± 4.7% vs 5dpf: 5.6 ± 0.6%, p<0.01) (Fig. 2A, Table S4). Additionally, peptides from Mybpha, encoded by *mybpha*, the second zebrafish *MYBPH* ortholog, were also detected in adult samples, though that protein accounted for less than 2% of total MyBP (Table S4). In all, MyBP abundance relative to myosin heavy chain was higher in adult than in larval samples (adult: 0.105 ± 0.006 vs. 5dpf: 0.082 ± 0.008, p<0.01), but still in line with those previously reported in mammalian muscle types (19, 26).

**Figure 2.**
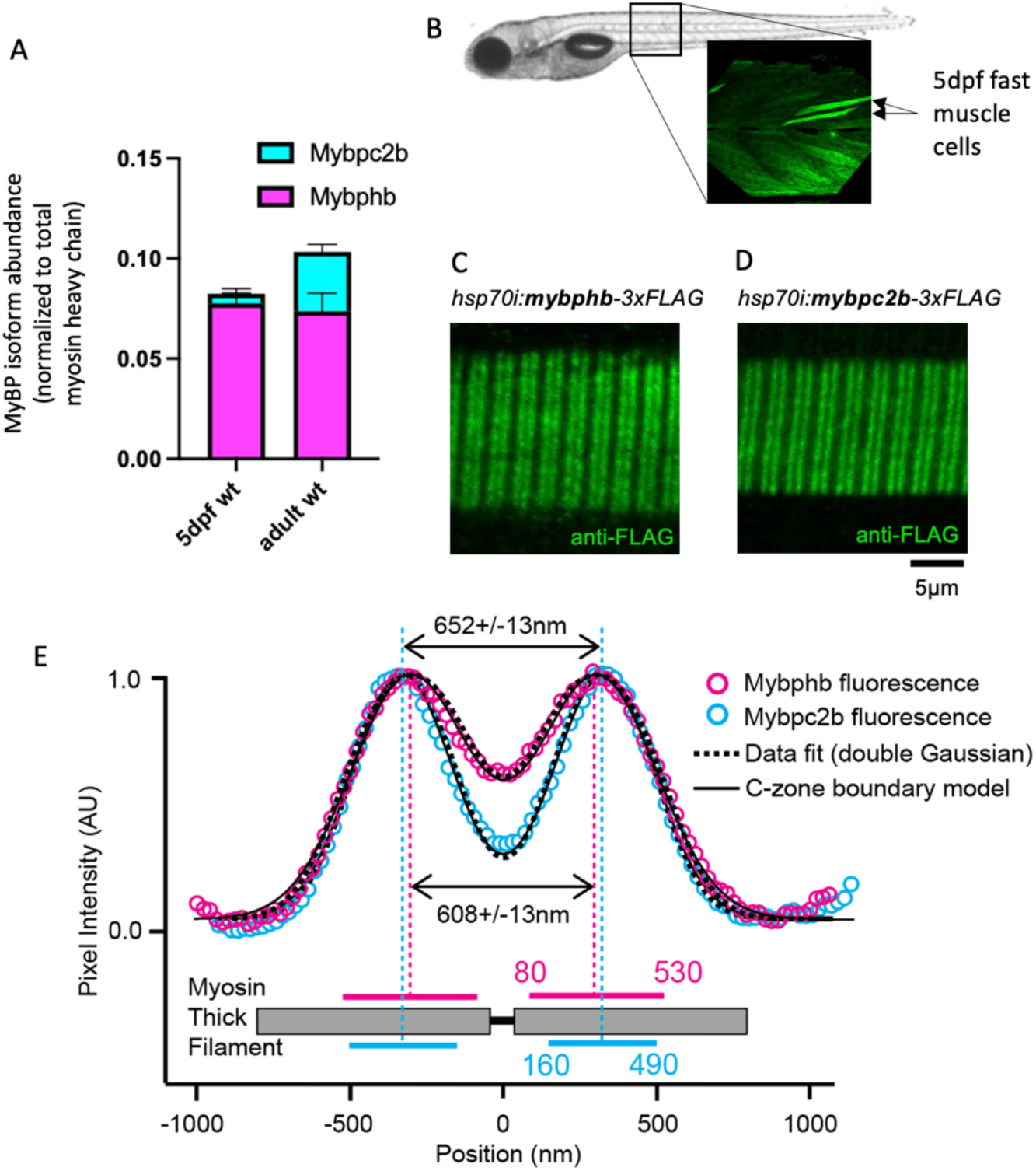
Quantification and distribution of MyBP-C and MyBP-H isoforms in fast-twitch zebrafish myotomal muscle. (A) Quantitative LCMS of 5dpf larval tails and adult, fast-twitch muscle samples reveal two MyBP protein isoforms present at levels above the threshold for quantification: Mybphb (MyBP-H) and Mybpc2b (fMyBP-C) and presented as the relative molar ratio to total myosin heavy chain. B) Mosaic fluorescence from immunostained FLAG peptide in 5dpf larval tail injected with *hsp70i:mypbhb-3xFLAG* construct 24 hours after heat-shock. Fast-twitch myotomal muscle cells (arrows) have a typical trapezoidal shape and off-axis orientation. (C, D) In fast-twitch larval muscle cells expressing transgenic *hsp70i:mypbhb-3xFLAG* (C) or *hsp70i:mypbc2b-3xFLAG* (D), anti-FLAG antibodies label two bands in each sarcomere, creating a fluorescence doublet pattern along the fiber. (E) For each construct, the aligned and integrated intensity of these two bands (open circles) *(Materials and Methods)* are well fit with two Gaussian peaks (dashed curves) with separations of 608 ± 13 nm (*hsp70i:mypbhb-3xFLAG*), and 652 ± 13 nm (*hsp70i:mypbc2b-3xFLAG* ). Theoretical intensity profiles generated by an analytical model (solid curves) are fit to the experimental data by assuming that the antibody fluorescence is equally distributed across regions of the thick filament, bounded at points 80 nm and 530 nm (Mybphb-3XFLAG) or 160 nm and 490 nm (Mybpc2b-3XFLAG) from the sarcomere center. These regions correspond well to the dimensions and location of the mammalian skeletal muscle ‘C-zone’. Data are presented as means ± 1 S. D..

Based on the amount of MyBP-H protein present in the zebrafish muscles, in the form of Mybphb, we hypothesized that it might be bound to more than the single C-zone ‘stripe’ previously observed in adult mammalian fast-twitch muscle (1). To test this hypothesis, we injected zebrafish embryos with transgene constructs encoding either Mybphb or Mybpc2b with a FLAG-tag fused to the C-terminal end of the protein under control of the *hsp70i* heat-shock promoter (see Methods) (32, 33). The larvae were heat-shocked at 4dpf to induce transgene expression and fixed 24 hours later (see Methods). Fixed larvae were then labeled using a monoclonal antibody against the FLAG-tag and fluorescently labeled secondary antibody (34). The protein localization appeared mosaic, reflecting transgene integration into the genomes of a subset of cells (Fig. 2B). Fast-twitch myotomal muscle fibers from larvae carrying either the *hsp70i:mybphb-3xFLAG* or the *hsp70i:mybpc2b-3xFLAG* transgene showed intracellular fluorescence localized to repeating doublets resembling MyBP-C C-zone staining patterns observed in other muscle types (Fig. 2C, D) (2, 19, 35).

To define the spatial distribution of the FLAG-tagged Mybphb and Mybpc2b proteins within these sarcomeres, we modified a analytic model previously used to define the localization of MyBP-C within rat muscle sarcomeres (see (19) and Methods). In brief, to improve the fluorescence signal-to-noise ratio, we aligned and averaged the fluorescence doublets from multiple sarcomeres and then fitted the resultant intensity profile by the sum of two Gaussians (Fig. 2E, dashed curves). The spacing between the peaks of the two Gaussians was 608 ± 13 nm for Mybphb and 652 ± 13 nm for Mybpc2b (Fig. 2E, dashed vertical lines), while peak widths were 184 ± 10 and 148 ± 12 nm, respectively. This fit was then used to constrain a model, which generated a dual-Gaussian fluorescence profile based on the point-spread function of the antibodies’ fluorophores (Fig. 2E, solid curves). Due to there being no previous direct structural data in zebrafish regarding MyBP localization within the sarcomere as in mammals (i.e. “stripes”), we based the model on an assumption of uniform distribution of fluorophores within bounded regions (i.e. C-zones) either side of the sarcomere center. The model then iteratively set the location of the inner and outer boundaries, comparing the predicted fluorescence profile to the fitted experimental data. The best fit to Mybphb-FLAG fluorescence was generated by a region of the myosin thick filament with boundaries 80 and 530 nm from the thick filament center (Fig. 2E, solid horizontal magenta lines). Mybpc2b-FLAG data was best fit by a narrower band of fluorescence with 160 and 490 nm boundaries (Fig. 2E, solid horizontal cyan lines). In a mammalian thick filament, these boundaries would correspond approximately to stripes 1-10 (Mybphb-FLAG) and 2-9 (Mybpc2b-FLAG), similar to the C-zone distributions reported for sMyBP-C (stripes 2-11) and fMyBP-C (stripes 4-11) in rat fast-twitch EDL muscle (19). These results suggest Mybphb can bind to thick filaments across the full range of binding sites accessible to Mybpc2b and beyond.

### Genetic depletion of MyBP-H in zebrafish is not fully compensated by increase in other MyBP isoforms

To determine whether Mybphb is necessary for normal muscle development and function, we used a CRISPR/Cas9 based approach to generate deletion alleles in zebrafish *mybphb* (See Methods) (36). Briefly, we injected single-cell wildtype embryos with Cas9 protein pre-complexed to guide RNAs targeted to two PAM sites situated in exons 1 and 4, raised larval F0s to adulthood, and recovered F1 embryos transmitting a ∼12 kb deletion of the intervening genomic DNA. The deletion allele was confirmed by PCR using a primer pair spanning the deletion. In addition to the deletion, Sanger sequencing of the resulting amplicon revealed a +2 frameshift, resulting in multiple downstream stop codons in the remnant exon 4 (Fig. S1B).

To determine how the genetic perturbation impacted protein accumulation, we performed LCMS analyses (see Methods) on five-day-old (5dpf) larval tails, heterozygous or homozygous for the deletion (Fig. 3A). These analyses demonstrated that the Mybphb to myosin heavy chain ratio was reduced (P<0.01) to ∼65% of wildtype levels in heterozygotes, indicating haploinsufficiency. No peptides unique to Mybphb were found in homozygous null samples (Fig. 3A), reflecting a complete loss of gene expression (Table S4).

**Figure 3.**
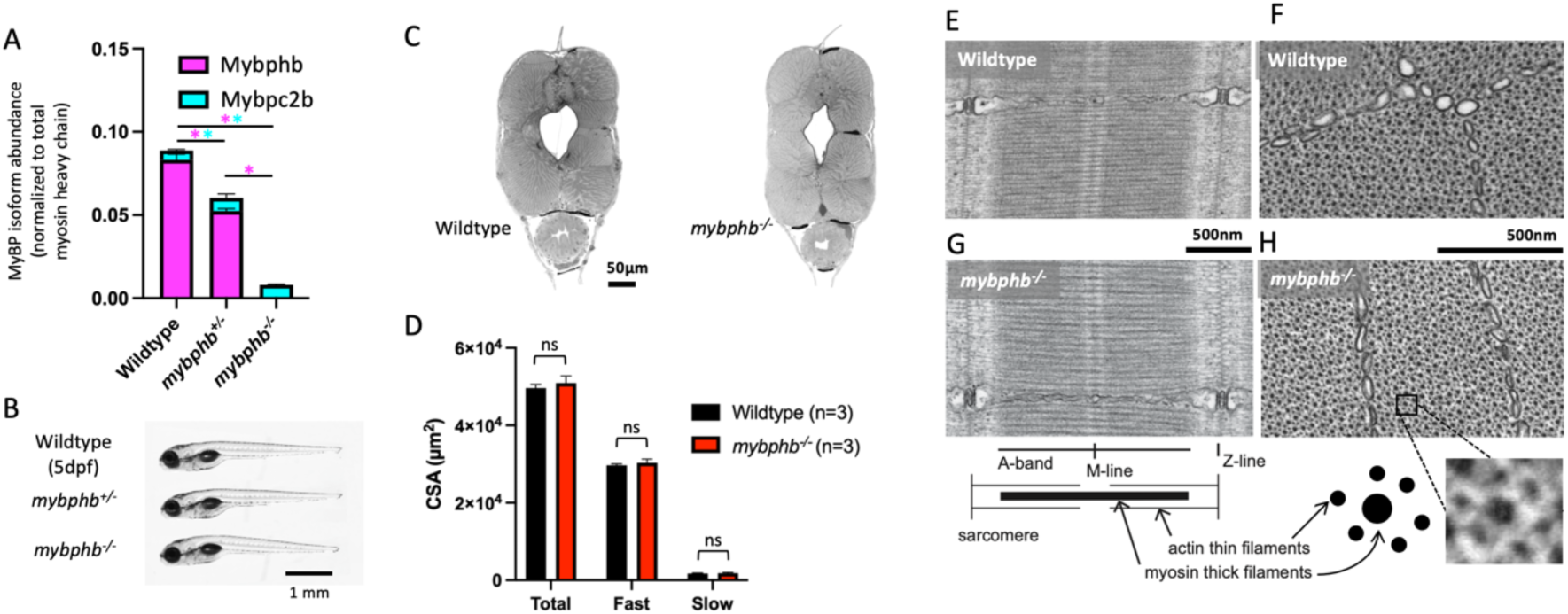
Effect of a *mybphb* deletion allele on MyBP isoform accumulation and morphology in 5dpf larval tails. (A) Quantitative LCMS of tails from heterozygous (*mybphb^+/-^* ) and homozygous (*mybphb^-/-^*) larvae show reduction and ablation of Mybphb, respectively, compared to wildtype. Statistics were calculated using a two-tailed Student’s t-test. Magenta or cyan ‘*’ denote p < 0.05 between bars of the corresponding colors. In each case, Mybpc2b accumulation increases slightly, but does not fully compensate for the loss of Mybphb. (B-D) *Mybphb^-/-^* larvae display no gross morphological phenotype or change in relative or absolute fast-twitch or slow-twitch muscle cross-section area. (E-H) TEM images of longitudinal (E,G) and transverse (F, H) ∼80 nm sections of 5dpf fast muscle cells reveal no deviations from normal muscle ultrastructure in the absence of Mybphb. Statistics were calculated using a two-tailed Student’s t-test. ‘ns’ denotes p > 0.05. Data are presented as Means ± S.D..

Due to the presence of Mybpc2b (Fig. 2A, 3A) and the overlap in its localization with Mybphb (Fig. 2C, D) in 5dpf wildtype larva, we hypothesized that the expression of Mybpc2b may be upregulated to account for the loss of Mybphb. The LCMS data demonstrate that the abundance of Mybpc2b is increased by a small but significant (P<0.01) amount relative to myosin heavy chain in both heterozygous *mybphp ^+/-^* and homozygous *mybphb^-/-^* larvae at 5dpf (Fig. 3A, Table S4). However, these compensatory increases in Mybpc2b, were not sufficient to provide the wildtype abundances of MyBP. The total abundance of MyBP was significantly reduced to ∼70% of wildtype levels in heterozygous and ∼8% of the wildtype levels in homozygous larvae. No additional peptides were identified from other MyBP isoforms. Loss of Mybphb protein did not affect the relative abundance of individual myosin heavy chain isoforms, as indicated by the quantification of unique peptides (Methods, Table S4).

### Larval muscle ultrastructure is unaffected in the absence of the zebrafish MyBP-H isoform (Mybphb)

Zebrafish heterozygous (*mybphb^+/-^*) or homozygous (*mybphb^-/-^*) for the deletion allele with their reduction or total loss of Mybphb protein, respectively, progress normally from embryogenesis to adulthood, showing no obvious deviation from wildtype siblings in body size or appearance (Fig. 3B). MyBP-C gene mutations have wide ranging morphological effects in other muscle settings (3–6, 37). Therefore, to determine whether the absence of Mybphb impacts larval muscle size or subcellular ultrastructure, we fixed and mounted tails from 5dpf wildtype and *mybphb^-/-^* larvae and imaged longitudinal and cross-sections by brightfield and transmission electron microscopy. The fast-twitch fibers in larval myotomal muscles undergo both rapid hyperplasia and hypertrophy in the first days of life (38). To quantify muscle size in 5dpf larvae, we stained wildtype and *mybphb^-/-^* tail cross-sections with toluidine blue for contrast and imaged them in brightfield (Fig. 3C) (see Methods). Myotomal muscle architecture is well described, and fast- and slow-twitch muscle fibers are spatially distinct, allowing for the quantification of total fast- and total slow-twitch muscle cross sectional area (CSA) relative to the CSA of the whole tail by manual analysis in Fiji/ImageJ (see (22) and Methods). At 5dpf, *mybphb^-/-^* tail CSA immediately cranial to the anal vent was (5.09x10^4^ ± 1.80x10^3^ µm^2^) not different from wildtype controls (4.93x10^4^ ± 1.28x10^3^ µm^2^) (Fig. 3D). Combined fast- and slow-twitch muscle CSA made up 3.22x10^4^ ± 0.77x10^3^ µm^2^ of total tail CSA in *mybphb^-/-^*, the bulk of which (94.6 ± 0.5%) consisted of fast-twitch muscle. Slow-twitch muscle accounted for the remaining 5.4 ± 0.5%, which at this stage occupies a ∼5 µm thick, single-cell layer just under the skin (22). None of these values differed significantly from wildtype controls indicating that loss of Mybphb protein has little or no effect on larval muscle growth under normal rearing conditions.

To define ultrastructural phenotypes in fast-twitch zebrafish muscle fibers associated with Mybphb loss, ultrathin (80 nm) longitudinal- (Fig. 3E, G), and cross-sections (Fig. 3F, H) were taken from fixed *mybphb^-/-^* (n=3) and wildtype (n=3) tails and imaged by TEM (see Methods). A qualitative assessment of key cellular and myofibrillar features revealed no deviations from wildtype muscle ultrastructure in mutant samples. Specifically, sarcomere length and general organization appeared unchanged. In cross section, the myofibrillar lattice appeared normal, as did overall cell shape as well as the shape, location, and distribution of myofibrils and other cellular structures.

### The absence of Mybphb protein blunts the effect of passive stretch and contraction on myofilament lattice compression

Recent structural studies show MyBP-C contributes to the spacing of the thick and thin filaments within the sarcomere, specifically to the compression of the myofilament lattice that occurs when muscle is stretched to longer sarcomere lengths (39, 40). To determine whether the absence of Mybphb has an impact on lattice spacing and compression with muscle length in live intact 5dpf tails, we used small angle x-ray scattering. This approach takes advantage of the multiple nanometer-scale periodicities formed by the ordered arrangement of molecular structures in myofibrils, each of which diffracts x-rays according to spacing and orientation (Fig. 4A) (41, 42). The 2D x-ray scattering patterns (see Methods) were characterized by two reflections along the equatorial axis (Fig. 4B, C). These reflections, called 1,0 and 1,1, are inherent to the repeating hexagonal arrangement of myosin thick filaments and actin thin filaments in vertebrate muscle. The 1,0 reflection is caused by parallel planes consisting entirely of myosin thick filaments (Fig. 4A), whereas the 1,1 reflection is caused by planes containing both myosin and actin. At resting length, lattice spacing determined from the position of 1,0 (d1,0) and 1,1 (d1,1) reflections in *mypbph^-/-^* tails were 43.0 ± 0.4 nm and 24.9 ± 0.2 nm (n=8) respectively, not statistically different from wildtype 42.9 ± 0.5 nm and 24.9 ± 0.3nm (n=7) (Fig. 4D, E). However, when tails were passively stretched to 110% of resting length and re-imaged, we observed a small but significant effect of the absence of Mybphb on the resulting compression of the lattice. The d1,1 spacing in the wildtype was reduced by 0.7 ± 0.2 nm after stretch, as expected from previous studies (43, 44), whereas in the *mybphb^-/-^* lavae, the reduction was 0.2 ± 0.4 nm (P<0.05) (Fig. 4D, E). A similar trend was observed in the D1,0 spacing, though the effect was not significant.

**Figure 4.**
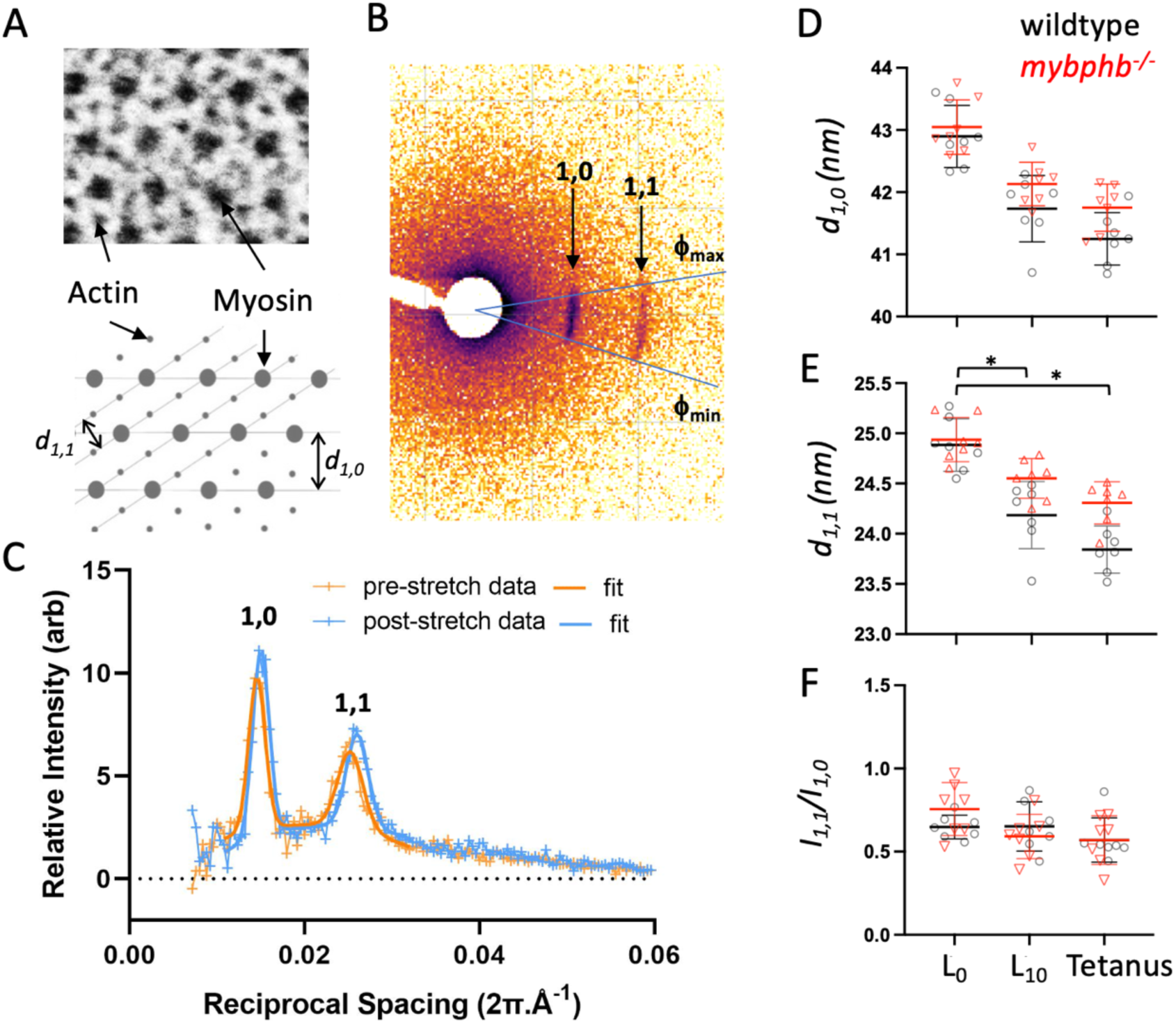
Small-angle x-ray scattering from live 5dpf wildtype and *mybphb^-/-^* tails. (A-C) The distances between planes in the paracrystalline lattice formed by myofibrillar actin and myosin filaments are measured by the spacing of equatorial x-ray scattering reflections. (A) TEM of zebrafish fast-twitch muscle cross section and schematic showing the distribution of actin and myosin filaments, and major spatial planes created by the array of actin, thin and myosin, thick filaments. (B) Representative x-ray scatter showing 1,0 and 1,1 equatorial reflections caused by planes in (A). Circumferential spread of the reflections around the origin is caused by inhomogeneity of muscle fiber angles within the tail. (C) Representative plot of background-subtracted radial intensity over the angle phi showing position and relative intensity of 1,0 and 1,1 reflections before and after stretch. (D, E) Spacing of d_1,0_ (D) and d_1,1_ (E) planes in wildtype (n=7) and *mybphb^-/-^*(n=8) tails at rest length (L_0_), after being stretched to rest length + 10% (L_10_), and during a 300 Hz, electrically stimulated tetanic contraction at L_10_ (Tetanus). Loss of Mybphb resulted in significant reductions in d_1,1_ lattice compression (genotype x treatment effect) with stretch (p=0.039) and tetanus (p=0.009), but not d_1,0_ (stretch, p=0.219; tetanus p=0.138). (F) An increase in the ratio of 1,1 to 1,0 reflection intensities (I_1,1_/I_1,0_) can indicate a shift of mass from thick to thin filaments (i.e. myosin heads moving towards or attaching to the thin filament). No such shift was seen in either group with stretch or tetanic stimulation. Statistics were performed using 2 way Analysis of Variance. ‘*’ denotes a genotype x treatment effect with P < 0.05. Data are presented as means ± 1 S.D..

To test whether the absence of Mybphb affects lattice spacing in active muscle, tails were subsequently imaged during electrically stimulated tetanic contractions. Our previous work demonstrated that myotomal sarcomeres within tails stretched to 110% of rest length, will shorten during tetanic contraction back to 100% rest length due to the series elasticity within the preparation, enabling us to compare passive (Fig. 4 D-F; “L0”) and active (Fig. 4 D-F; “Tetanus”) at the same sarcomere length. As with passive stretch, lattice compression associated with contraction was also reduced in the *mybphb^-/-^* tails, with the difference being statistically significant for d1,1 (1.1 ± 0.2nm, wildtype; 0.6 ± 0.3nm, *mybphb^-/-^*; p=0.009), but not d1,0 (1.6 ± 0.4nm, wildtype; 1.2 ± 0.7nm, *mybphb^-/-^*; p=0.138)(Fig. 4D, E). The ratio of the 1,1 to 1,0 reflection intensities (I1,1/I1,0) can also be informative, as the motion of myosin heads towards actin thin filaments shifts intensity from the myosin thick filament only 1,0 plane to the 1,1 plane containing both myosin thick- and actin thin-filaments. No significant changes in I1,1/I1,0 were observed with stretch or stimulation in these samples. This was not entirely unexpected given that passive stretch did not activate the muscle resulting in myosin heads attaching to the actin thin filament. Additionally, upon contraction, the rapid kinetics of myosin head attachment to and detachment from the thin filament for these ultrafast larval tail muscles (22) suggests that relatively few myosin heads are attached to the actin thin filament and generating force at any one time. Therefore, the expected change in the I1,1/I1,0 intensity ratio should be minimal. Collectively, these lattice compression data suggest MyBP-H, like MyBP-C may contribute to myofilament lattice mechanics in ways that could affect contractility.

### Larval tail muscle function is unaffected by the absence of Mybphb

To quantify the effect of the absence of Mybphb on muscle function, we analyzed mutant larvae for changes in locomotory behavior and performed intact mechanical analyses of tail muscle function. Within days of fertilization, larvae respond to mechanical stimulus with a stereotypical startle response, which includes a burst of swimming from a series of rapid powerful contractions of myotomal muscles (45). To identify locomotory deficits associated with loss of Mybphb, we observed the response of 10dpf wildtype, and *mybphb^-/-^* sibling larvae to a tap stimulus. No genotype effect was observed in either average velocity (Fig. 5A) or total distance traveled (Fig. 5B) following the stimulus.

**Figure 5.**
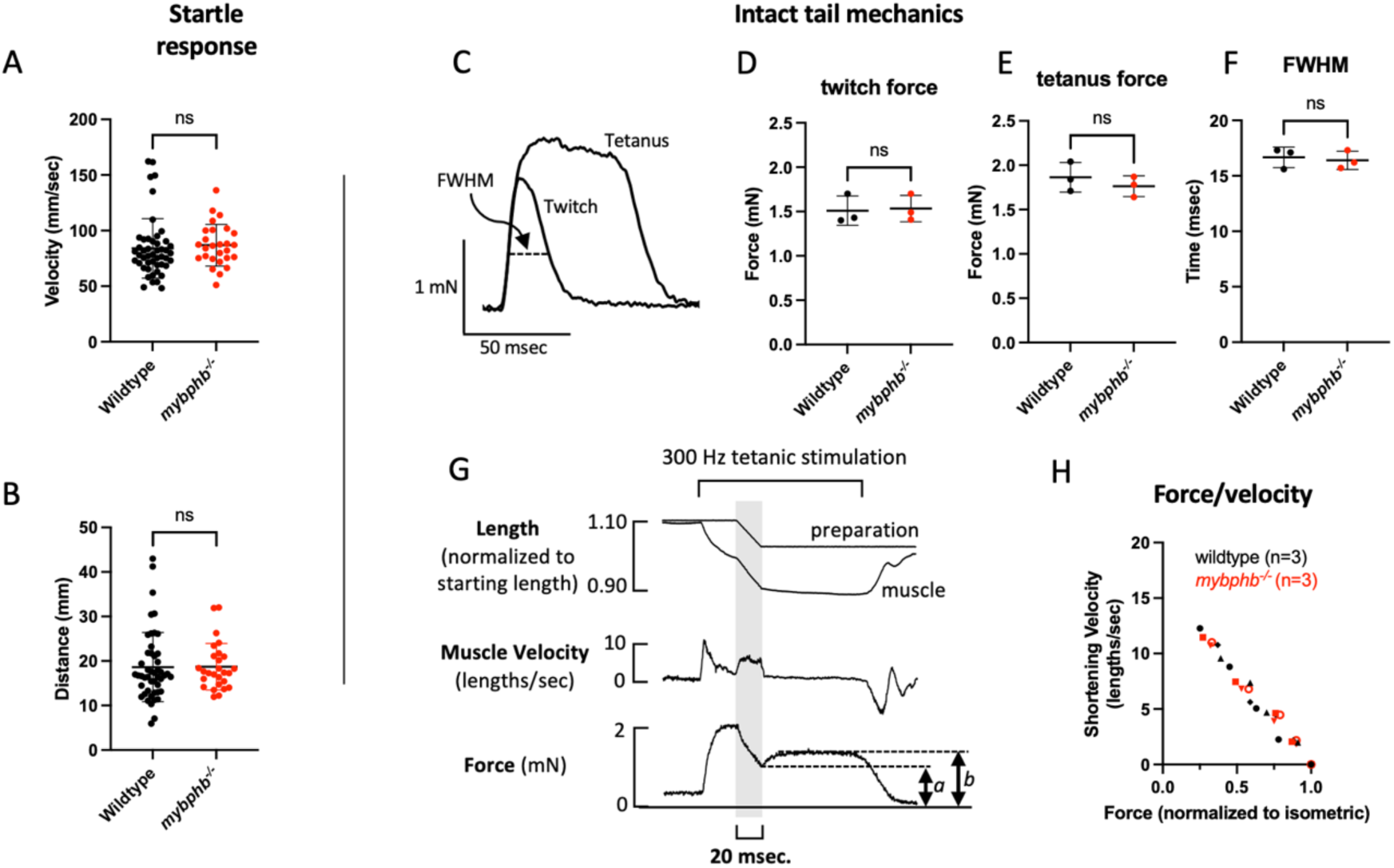
Myotomal muscles from *mybphb^-/-^* larvae function normally. (A, B) 10dpf larvae react to a mild mechanical stimulus with a brief burst of high speed swimming. No difference was observed between wildtype (n=48) and *mybphb^-/-^* (n=27) sibling larvae in average velocity during the escape maneuver (A) or total distance traveled (B). (C-H) *In vitro* mechanical function of 5dpf larval tails (Wildtype n=3, *mybphb^-/-^* n=3). *Mybphb^-/-^* larvae developed normal forces in response to a single 0.4 msec electrical stimulus (twitch, D), and to a 100 msec, 300 Hz train of stimuli (tetanus, E). (F) Twitch full width at half maximum (FWHM), which depends on rates of activation and relaxation, was also unaffected. (G, H) To measure the force/velocity relationship, tails were allowed to shorten at a series of fixed velocities during tetanic stimulation. Active force at the end of each ramp (*a*) was normalized to isometric force after recovery (*b*) and plotted against muscle velocity in (H). No difference in force/velocity was seen over the range of velocities possible within the limits of the instrumentation. Significance was determined using a two-tailed Student’s t-test. ‘ns’ denotes P > 0.05. Data are presented as Means ± 1 S.D..

We next evaluated the mechanical function of larval myotomal muscles directly. The arrangement of myotomal muscle fibers in parallel with the long axis of the tail enables fundamental muscle mechanical parameters to be measured in intact larvae. We clamped 1 mm long tail segments from 5dpf wildtype and *mybphb^-/-^* siblings between a force transducer and a length servo and measured the responses to electrically-evoked twitch and tetanic contractions as described previously (22) and in Methods (Fig. 5C). No genotype effect was observed on maximal twitch force (wildtype: 1.51 ± 0.17 mN, n=3 vs. *mybphb^-/-^*: 1.53 ± 0.15 mN, n=3; P>0.05) (Fig. 5C, D) or twitch duration as measured by full width at half max (FWHM) (wildtype: 16.7 ± 0.9 msec, n=3 vs. *mybphb^-/-^*: 16.4 ± 0.8 msec, n=3; P>0.05) two parameters that depend on rates of force development and relaxation, as well as calcium transients (Fig. 5C, F). The same was true for maximal tetanic force (wildtype: 1.86 ± 0.17 mN, n=3 vs. *mybphb^-/-^*: 1.76 ± 0.12 mN, n=3; P>0.05) indicating isometric steady-state force generating capacity is not dependent on Mybphb (Fig. 5C, E). However, twitch and tetanus do not capture the muscle’s ability to produce mechanical power, which is dependent on both force and shortening velocity. Therefore, we measured the force/velocity relationship by allowing tails to shorten at fixed velocities during tetanic contractions (Fig. 5G; see Methods). This analysis showed no difference in force/velocity relationship between *mybphb^-/-^* larvae and their wildtype siblings (Fig. 5H) at velocities up to ∼12 sarcomere lengths/second. Importantly, limitations inherent to the preparation and the apparatus prevented accurate comparisons at higher velocities, or predictions of maximum unloaded shortening velocity (Vmax).

### Mybphb slows actin filament motility in native thick filament C-zones

Genetic depletion of Mybphb in zebrafish larvae enabled us to make functional and structural comparisons between wildtype muscles containing both Mybphb and Mybpc2b protein versus mutant muscles (*mybphb^-/-^*) without Mybphb but with a remaining fraction of Mybpc2b protein (Fig. 3A). In order to study Mybphb function directly and in isolation, we performed *in vitro* motility experiments using muscle from a second mutant zebrafish line (see Methods) carrying a nonsense mutation in the *mybpc2b* gene *(SA10810,* Fig. S1C) so that fish homozygous for the mutation were effectively a *mybpc2b* null (i.e. *mybpc2b^-/-^*). As with *mybphb^-/-^* , the *mybpc2b^-/-^* embryos develop normally to adulthood with no obvious phenotype (Fig. S1C). This was in contrast to an earlier study, which showed a severe muscle phenotype after knock down of *mybpc2b* using morpholinos, although that technique has been associated with off-target effects (46, 47). Quantitative LCMS of fast-twitch myotomal muscle from *mybpc2b^-/-^* adults demonstrated the absence of Mybpc2b peptides with no apparent change in the distribution of myosin heavy chain isoforms (Fig. 6A, Table S3). Interestingly, the abundance of Mybphb was enhanced in the absence of Mybpc2b to maintain the ratio of MyBPs relative to myosin heavy chain present in the adult wildtype zebrafish (Fig. 2A), indicating that sarcomere C-zones in these muscles are fully occupied by Mybphb (Fig. 6A, Table S4).

**Figure 6.**
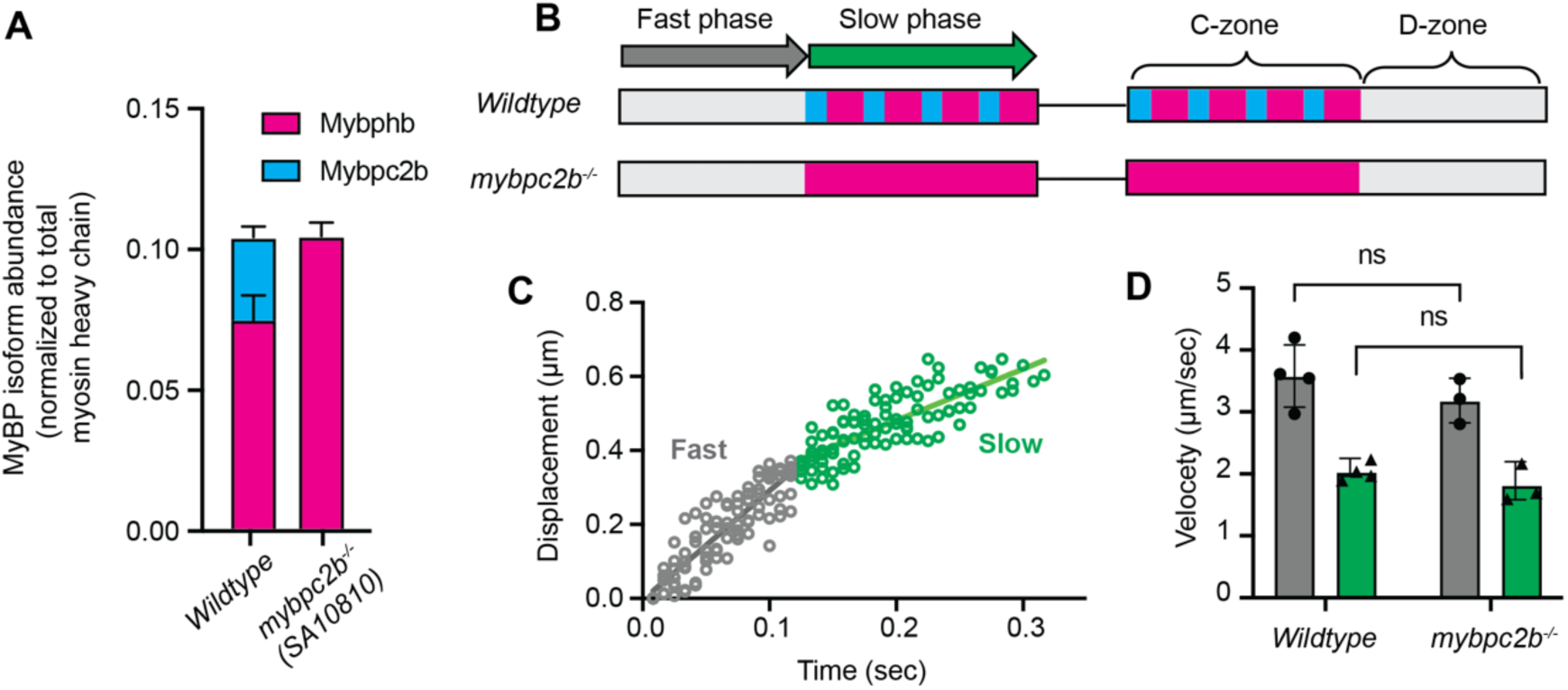
Mybphb acts as a molecular “brake” in the C-zone. (A) Abundance of MyBP isoforms in fast-twitch muscle samples from adult wildtype and *mybpc2b^-/-^* zebrafish relative to total myosin heavy chain. A homozygous null mutation in *mybpc2b* (SA10810) ablates Mybpc2b and increases the abundance of Mybphb, maintaining a consistent molar ratio of MyBPs to myosin heavy chains observed in the wiltype zebrafish. (B-D) The sliding of fluorescently labeled actin filaments along native myosin thick filaments isolated from wildtype and *mybpc2b^-/-^*adult zebrafish fast muscle. (B) An illustration of wildtype and mutant thick filaments with C-zones colored to represent MyBP isoform content as in A. The native thick filament assay observes the movement of a fluorescently labeled actin filament being propelled over the thick filament surface by the myosin heads emanating from the thick filament. Actin filament displacement trajectories have an initial fast phase associated with the D-zone, which is devoid of MyBP, followed by a potential slower phase as it moves over the MyBP within the C-zone. (C) Actin filament displacement vs. time data for 13 different actin filaments moving over different wildtype fast-twitch muscle thick filaments demonstrating dual-phase trajectories as illustrated in B with the fast velocity phase (grey data) and slow velocity phase (green data) identified. (D) The fast (grey) and slow (green) velocity phases for actin filament motion over wildtype thick filaments containing a mix of Mybphb and Mybpc2b (74 thick filaments from n=4 animals) as in A and *mybpc2b^-/-^*thick filaments containing exclusively Mybphb (116 thick filaments from n=3 animals). Statistics were calculated using a two-tailed Student’s t-test. ‘ns’ denotes P > 0.05. Data are presented as means ± 1 S.D..

We previously showed that fluorescently labeled actin filaments slow when entering the C-zone of native, rat skeletal myosin thick filaments where MyBP-C alone resides (Fig. 6B), demonstrating its capacity as a molecular “brake”. Importantly, actin filament slowing was eliminated in the C-zone when the MyBP-C N terminus was proteolytically cleaved (19). Since Mybphb, lacks analogous N-terminal domains, we hypothesized that thick filaments containing Mybphb exclusively would be similarly unable to slow actin, which would make the “braking” capacity in the C-zone dependent on the relative abundance of Mybpc2b with its N terminal domains. Therefore, we isolated native thick filaments from the *mybpc2b^-/-^* and wildtype adult fast-twitch myotomal muscle and performed a single filament sliding assay (see Methods). The native thick filaments were deposited onto the surface of a flow cell and the sliding of ∼250 nm long fluorescently labeled actin filaments on these thick filaments was quantified in the presence of 100 μM ATP at room temperature (23° C). Approximately, 70% of the displacement trajectories (Fig. 6C) on wildtype thick filaments, which has a ∼75/25 ratio of Mybphb to Mybpc2b (Fig. 6A), showed two distinct velocities over distances that corresponded to predicted D-(devoid of MyBPs) and C-zone dimensions. Specifically, these trajectories were characterized by an initial fast phase of velocity lasting ∼350 nm that transitioned to a ∼50% slower phase of velocity lasting ∼400 nm (Fig. 6C, D), initially assumed to be due to the presence of Mybpc2b and its N-terminal domains. Despite our expectations, actin filaments sliding on thick filaments from the *mybpc2b^-/-^* fish that only contained Mybphb protein, also demonstrated two distinct phases of velocity (Fig. 6A) that were similar to that observed on wildtype thick filaments (Fig. 6D). These data suggest that the “braking” capacity of the C-zone is not solely dependent on Mybpc2b and that Mybphb itself is sufficient to act as a molecular “brake”.

## Discussion

The sarcomere C-zone is considered a nexus for striated muscle structure and function, due to the presence of MyBP-C, a critical contractile modulator (10, 19). Adult mammalian skeletal muscles express sMyBP-C and fMyBP-C isoforms from 2 different genes, *MYBPC1* and *MYBPC2*, respectively, with *MYBPC1* subject to alternative splicing (48). The expression of these two isoforms allows for the contractile modulation to be fine-tuned (19) to meet the diverse physiological demands of skeletal muscles. Both the *MYBPC1* and *MYBPC2* genes have garnered significant attention, as mutations in these genes are linked to various skeletal muscle myopathies (4–6, 37). In contrast, MyBP-H, being another member of the MyBP family that co-occupies the skeletal muscle C-zone, has received little attention, in part due to its minimal expression in adult mammalian muscle. Here we report that MyBP-H is highly expressed in prenatal mammalian fast-twitch skeletal muscles, and its expression may be common to developing fast-twitch muscles across vertebrate species. Moreover, we demonstrate that *mybphb*, an ortholog of mammalian *MYBPH* is highly expressed in zebrafish, and therefore take advantage of the ability to manipulate its expression in this model system to demonstrate that MyBP-H may be playing an unexpected role in the modulation of actomyosin function.

### MyBP-H structure and subcellular localization during muscle development

Nearly 40 years ago, the presence of MyBP-H was identified as a “novel thick filament protein” that shared C-zone occupancy with sMyBP-C in embryonic chicken fast-twitch pectoralis (14). In the present study, we identified the expression and quantified the abundances of MyBP-H, sMyBP-C, fMyBP-C, as well as myosin heavy chain isoforms in rat hindlimbs over a developmental period spanning initial myofibrillogenesis (e14, e16 hindlimb buds) to muscle fiber growth and maturation (e18, e20, adult fast-twitch TA muscles) using mass spectrometry-based proteomics (Fig. 1C, D). Interestingly, MyBP-H expression appeared to be restricted to a brief window of time during the later stages of muscle development (e16-e18), a period characterized by rapid hypertrophy of established muscle fibers and the proliferation of a secondary wave of muscle fiber formation (27). At the earliest stage (e18) MyBP-H expression is high and exceeds sMyBP-C, the only other MyBP isoform present (Fig. 1D). By e20, the balance shifts in favor of sMyBP-C, which outnumbers MyBP-H by ∼2:1. Ultimately, in the adult rat TA muscle, MyBP-H abundance is minimal, with fMyBP-C and sMyBP-C being the predominant MyBP isoforms, as observed in other adult, mammalian fast-twitch muscle (11, 19). In addition, we report a similar finding in zebrafish, where MyBP-H (in the form of Mybphb, the product of the zebrafish *MYBPH* ortholog *mybphb*) is the predominate MyBP isoform in ultrafast swimming muscles of rapidly growing larvae, though expression remains high in adults (Fig. 2A). Therefore, with the earlier reports in chicken (49), these observations in rat and zebrafish mark three major vertebrate taxa in which MyBP-H is highly abundant during fast-twitch muscle development. This conserved expression pattern is in accord with our phylogenetic analysis, which shows that ancestral *MYBPH* arose around the same time as *MYBPC1, MYBPC2, and MYBPC3* (Fig. 1E), suggesting that MyBP-H, like its MyBP-C cousins, specialized early in vertebrate evolution, and that its role remains broadly conserved.

All members of the MyBP family, including MyBP-H, possess homologous C-terminal globular domains, which serve to anchor these proteins to the thick filament backbone (29, 49). Regardless of the functional/structural roles of MyBP-H, does it bind to and compete for the same sites on thick filaments when co-expressed with MyBP-C? Our proteomic data from the rat TA muscles demonstrate the combined abundance of MyBP-H and MyBP-C relative to all myosin heavy chain isoforms, remains relatively constant across the three developmental timepoints where MyBPs are present (e18, e20, adult). These data are consistent with MyBP-H and both sMyBP-C and fMyBP-C competing for a fixed number of binding sites within the thick filament C-zones, with their proportions potentially governed by the proteostatic regulation of each MyBP (19).

Using genetic approaches in the zebrafish, we were able to address the question of MyBP-H in the C-zone directly. The expression of FLAG-tagged MyBP-H (Mybphb) and fMyBP-C (Mybpc2b) in zebrafish larvae and subsequent immunofluorescent labeling allowed for the demonstration that both proteins localize to the same region of the sarcomere corresponding to the mammalian muscle C-zone, with only small differences in distribution (Fig. 2). This is similar to that observed for MyBP-H and sMyBP-C in embryonic chicken pectoralis (50), but quite different than the specific localization of MyBP-H in a single ultrastructural stripe adjacent to the M-line of adult rabbit psoas muscle (1). In addition to sharing the C-zone, our data further indicate that MyBP-H and fMyBP-C likely compete for the same binding sites. Specifically, genetic depletion of either MyBP-H (*mybphb^-/-^*) or fMyBP-C (*mybpc2b^-/-^*) results in an increase in abundance of the remaining isoform (Fig. 3A, 6A). Specifically, fMyBP-C (Mybpc2b) marginally increases from ∼5% of the total wildtype MyBP abundance to ∼8% in *mybphb^-/-^*larval tails (Fig. 3A), whereas in adult *mybpc2b^-/-^* fast-twitch muscle, loss of fMyBP-C is fully compensated for by MyBP-H (Mybphb) (Fig. 6A). If C-zone occupancy for each of the MyBPs is related to the two MyBPs competing for shared binding sites, then once a competing MyBP is eliminated, at least some increased abundance of the remaining MyBP might be expected with or without compensatory upregulation at the transcript level. Such a ‘design principle’, in which opposing isoforms compete for a share of available biding sites, could theoretically allow the cell to more finely tune the relative abundance of different isoforms, and thus the overall modulatory effect of the C-zone. This simple mechanism could also explain the replacement/compensation of sMyBP-C for fMyBP-C and vice versa in skeletal muscles of genetic knockout mouse models that normally express both isoforms (39, 51).

Does MyBP-H expression in the context of muscle development provide evidence for its role? In addition to our proteomic data demonstrating that MyBP-H is highly expressed in developing muscles, a growing number of muscle gene expression studies have shown MyBP-H to be upregulated in settings where muscle remodeling, regeneration, and hypertrophy are common themes such as in resistance training (52), Duchenne muscular dystrophy (53) and amyotrophic lateral sclerosis (54, 55). Thus, the expression of MyBP-H may be part of a muscle development and repair program. While the role of the MyBPs during development is not well understood, there is evidence that dysregulation of the C-zone MyBP occupancy can disrupt muscle development and structure. Mutations to sMyBP-C cause developmental muscle disease, such as distal arthrogryposis, lethal congenital contracture syndrome-4, and myogenic tremors in humans (6, 37, 56). These diseases feature hypercontractile phenotypes, which result in limb contractures and malformation *in utero*. In a recent study of sMyBP-C knock-out mice, homozygote pups do not survive more than 24 hours post-birth displaying congenital limb contractures and respiratory distress due to an atrophied diaphragm (51). Even transient reduction of sMyBP-C by CRISPR/Cas9 approaches in adult mouse FDB muscle results in compromised sarcomere structure and force production (57). Human disease phenotypes have also been recapitulated in zebrafish by mutations in *mybpc1*, which encodes zebrafish sMyBP-C isoforms (58). Although no known mutations to *MYBPH* have been reported that result in developmental myopathies, the gene’s broad conservation and high expression levels during development strongly indicate a contribution to overall muscle development and physiology, even if its role is a subtle one. The period of TA muscle development represented by e18 and e20 is complex and heterogeneous, with rapid hypertrophy of existing fibers progressing alongside the generation of new fibers via myoblast fusion. (27). The distribution of MyBP-H and sMyBP-C among these populations of muscle cells will need to be explored.

The replacement of MyBP-H by MyBP-C isoforms in fast-twitch muscles of rat and, to a lesser extent, zebrafish going from developmental stages to adulthood suggests that the structural and/or functional roles of MyBP-H may be especially relevant in growing muscle (Figs. 1D, 6A). However, in the zebrafish, the complete absence of MyBP-H does not appear to grossly impact organismal and myotomal muscle development, since larvae appear morphologically normal (Fig. 3) and develop to adulthood and breed. Even at the sarcomere level, no overt ultrastructural defects are apparent (Fig. 3). This surprising observation does not rule out a functional role for MyBP-H within the C-zone. Interestingly, in the rat TA innervation and initial *in utero* activation of developing muscle fibers begins over the timeframe that the abundance of MyBP-H diminishes (Fig. 1D) and is supplanted by sMyBP-C as muscle fibers become more active at embryonic days 22 and 23 just prior to birth (27). Therefore, the impact of MyBP-H and sMyBP-C on contractility is likely to be relevant during these late stages of prenatal muscle development and depend on their relative abundance in the C-zone. A similar concept has recently been proposed whereby the normal mixture of MyBP-HL, a paralog of MyBP-H that is expressed only in mammalian atrial muscle, and cardiac MyBP-C (cMyBP-C), which together occupy the C-zone, are critical to regulating the kinetics of myofibril force relaxation (12). Therefore, comparisons of MyBP-H and MyBP-C molecular structure and function may offer clues to the functional role that MyBP-H serves when present.

### Functional impact of MyBP-H

MyBP-H shares structural homology with the four C-terminal, globular domains of sMyBP-C and fMyBP-C but is missing the six globular domains and their linkers that constitute the central and N-terminal regions of MyBP-C (Fig. 1B). The N terminus has long been considered the business end of the MyBP-C molecule. By extending away from the thick filament backbone, its binding partner interactions with adjacent myosin heads and actin thin filaments are believed to modulate contractility (18, 19, 37, 59–61). Specifically, the N terminus of both sMyBP-C and fMyBP-C beginning with its proline/alanine-rich N-terminal extension that is followed by the C1 and C2 Ig-like domains, linked by the flexible M-domain (Fig. 1B), are sufficient to serve as a “brake” to slow myosin generated thin filament movement and to sensitize the thin filament to calcium in *in vitro* motility assays (19). Without an equivalent N terminus, one might expect MyBP-H to be mechanically silent both *in vivo* and *in vitro*. However, this was not the case. In motility experiments using native, isolated thick filaments from adult MyBP-C2b null zebrafish fast-twitch muscles, in which only MyBP-H is present in the C-zone, we observed slowing of single actin filaments within the C-zone (Fig. 6). Therefore, MyBP-H can serve as a molecular “brake”, as seen previously in assays from both skeletal (19) and cardiac muscle (26, 62) in which MyBP-Cs occupy the C-zone and require intact N-terminal domains for the observed thin filament slowing.

In the absence of the N-terminal regions as in MyBP-C, what MyBP-H structural domains could be responsible for the observed “braking” action (Fig. 6)? MyBP-H from all species has a significant N-terminal proline/alanine-rich extension that is presumably unstructured and can range in length from 118 amino acids in zebrafish to 74 amino acids in humans. Such a long N-terminal extension could interact with myosin heads and/or the thin filament within the sarcomere to bring about the slowing observed in the native thick filament studies (Fig. 6). In fact, the proline/alanine N-terminal extension of the myosin essential light chain has been shown to interact with actin (63, 64) and by a similar interaction, MyBP-H could serve as an internal load to slow actin filament velocity (Fig. 6). Alternatively, two recent CryoEM structural studies of native human cardiac thick filaments have challenged the view that the MyBP-C N terminus is the sole determinant of MyBP-C’s modulatory capacity (65, 66). These high-resolution structural studies uncovered potential novel interactions between myosin motor domains on the thick filament surface with the MyBP-C C-terminal domains. These binding interactions may contribute to stabilizing the myosin interacting heads motif that effectively inhibits myosin’s ability to interact with the thin filament. Such an interaction, which involves domains shared between MyBP-C and MyBP-H, could influence the availability of active motors in the C-zone, thus contributing the observed slowing on the native thick filaments (Fig. 6).

Does the ability of MyBP-H to slow actin filament sliding along native thick filaments, translate into higher order mechanical effects in the zebrafish larvae? Although sarcomere structure in the MyBP-H null (*mybphb^-/-^*) larvae appears normal, the electron microscopic images are static pictures at a fixed length of myotomal muscle (Figs. 3E-H). Given the highly ordered structure of muscle fibers in the larval tail, small angle x-ray scattering was used (Fig. 4) to define the *in vivo* myofibrillar lattice spacing and its changes following stretch in live, relaxed 5dpf tails and following an activated tail contraction. As expected, both passive lengthening and active contraction of the tail muscles resulted in compression of the myofibrillar lattice spacing (Fig. 4D, E) due to constant muscle fiber volume properties (67) and myosin heads binding to the thin filament (68), respectively. However, in *mybphb^-/-^*these lattice spacing compressions were reduced, suggesting that MyBP-H does contribute to lattice spacing mechanics, presumably through its binding partner interactions. Interestingly, these results align with similar observations in a fMyBP-C knockout mouse (39). More recently, a lattice spacing expansion was also reported in mouse muscle fibers after removal of the N-terminal domains of fMyBP-C alone, an effect possibly mediated by loss of a mechanical connection between thick and thin filaments (40).This again raises the question of a possible actin-binding role for the proline/alanine extension of MyBP-H. How such reductions in lattice spacing compression in the absence of MyBP-H impacts muscle mechanical function, such as power production, would be speculative at best. Therefore, a direct assessment of tail muscle mechanics was performed.

Comparing wildtype and *mybphb^-/-^* larval tail mechanics, the absence of MyBP-H (Mybphb), does not appear to impact the maximum twitch and tetanic force or the kinetics of twitch activation and relaxation (Fig. 5C-F). In addition, the most basic physiological property of muscle, the Force/velocity relation, was also unaffected by the loss of MyBP-H (Fig. 5H), at least at low to moderate velocities, which could be probed by our instrumentation. If MyBP-H shares a similar “braking” action as MyBP-C, and potentially through a viscous drag effect on the thin filament that increases with increasing velocity (69), then the lack of an observed effect on maximum isometric force generation may not be surprising, i.e., viscous drag is zero under isometric conditions. Previous *in vitro* motility force/velocity studies using cardiac-specific MyBP-C, N-terminal fragments showed maximum isometric force, too, was not affected by the presence of the C0-C3 fragment, but that the force/velocity relation was shifted to lower velocities at each load due to a viscous drag effect (69). If MyBP-H does impose a viscous drag, then its greatest effect should be observed at maximum unloaded velocities. In fact, the native thick filament assay was under unloaded conditions where the slowing is observed. Due to limitations inherent to the intact tail muscle mechanics preparation, we were unable to probe MyBP-H’s viscous drag at these high velocities in the force/velocity relation (Fig. 5). Regardless, our data suggest that MyBP-H is not mechanically silent.

What is the significance of no structural or gross functional difference in *mybphb^-/-^* zebrafish? A key point is that the fish used in this study were reared and housed in a controlled laboratory setting, which lacks important stresses associated with life in the wild. Exposure to sub-ideal environmental conditions may uncover a loss of fitness or a deficit in an adaptive response associated with the loss of MyBP-H. For example, being ectothermic animals, fish are known to adjust muscle gene expression levels in response to changes in environmental temperature in order to maintain energetic and biomechanical homeostasis (70). It is possible that the composition of the C-zone is under such regulation. While *mybphb^-/-^*larvae show only a small increase in fMyBP-C (Mybpc2b) accumulation (Fig. 2) under laboratory conditions, an environmentally-induced upregulation of *mybpc2b* may overpopulate the C-zone with detrimental effects in the absence of competition from MyBP-H. The role of MyBP-H in mammalian development may overlap with that in the fish, but important contextual differences make predicting the effects of a null mutation in mammalian skeletal muscle difficult. Therefore, it will be critical to further investigate the expression timing and localization of MyBP-H in a mammalian model system.

### Conclusions

The regulation of contractile forces during myogenesis and development is critical, but the role of the C-zone in this process is not well understood. MyBP-H is expressed at high levels in rat fast-twitch muscle during a period late in fetal development after myogenesis, and in the myotomal muscle of highly motile larval zebrafish, indicating a broadly conserved developmental role. From its position in C-zone, MyBP-H is likely to have structural and mechanical effects both directly, through its own interactions with actin and/or myosin, and passively, through competition with other MyBP isoforms for binding sites within the C-zone. Despite lacking N-terminal globular domains analogous to those responsible for conferring MyBP-C’s modulatory properties, MyBP-H is capable of acting as a molecular “brake”. However, it is yet to be determined whether MyBP-H shares other properties attributed to MyBP-C, such as modulating actin thin filament calcium sensitivity. The remarkable absence of a strong structural or functional phenotype in MyBP-H null zebrafish myotomal muscle, despite its normally high abundance, points to the sometimes-subtle nature of C-zone based contractile regulation, which has made the interplay of multiple MyBP-C isoforms in mature skeletal muscle a challenge to study. The ability of zebrafish to develop normally with a nearly MyBP-free C-zone, opens the door to future experiments to dissect MyBP function through the transgenic introduction of individual MyBP isoforms, and versions with specific disease mutations.

## Materials and Methods

### Animal Handling and Ethics

All protocols were approved by the Institutional Animal Care and Use Committees of the University of Vermont (UVM), Washington University in St. Louis (WUSL), or Brookhaven National Laboratories (BNL), and were in compliance with the Guide for the Use and Care of Laboratory Animals published by the National Institutes of Health.

For proteomic analysis of rat limb muscle, timed-pregnant Sprague Dawley rats, 12-14 weeks of age and weighing 375 ± 25 g (Charles River Laboratories International, Quebec, CA), were obtained. Pregnant rats were single-housed, kept under pathogen-free conditions and had free access to standard food and water. The animals were allowed to acclimatize for at least 72 h prior to use at the UVM animal care core facility prior to being euthanized using 3% isoflurane (NDC 57319-507-06, Phoenix Pharmaceuticals Inc., USA) at gestational days 14, 16, 18 or 20. When no longer responsive to a hard pinch to the feet, rats were decapitated with a small animal guillotine. An abdominal incision was made and the uterine horn with conceptus removed. Fetuses were immediately removed and decapitated under cold PBS, and their anterior tibialis (TA) muscles were dissected and flash-frozen for processing. Additional incisions were made in the adult (dam) hindlimbs and skin exposed to reveal the underlying muscle structures. Samples of adult TA muscle were dissected and similarly processed.

Zebrafish were housed either at the UVM or WUSL at 28° C in laboratory aquarium systems under standard laboratory conditions. Embryos and larvae were maintained in E3 medium (5 mM NaCl, 0.17 mM KCl, 0.33 mM CaCl2, 0.33 mM MgSO4, 0.0001% Methylene Blue (Sigma-Aldrich, St. Louis, MO)) and incubated at 28°C before use. Unused larvae were anesthetized with buffered MS222 (Sigma-Aldrich, St. Louis, MO) (200µg/ml), and euthanized on ice. For mechanical, proteomic, x-ray and imaging analysis, 5dpf tails were removed by cutting immediately caudal to the yolk sack with a scalpel and processed accordingly. Heads and remaining body structures were frozen for genomic DNA extraction and genotyping. For proteomic analysis and myosin thick filament isolation from adult zebrafish, adult fish 6-18 months of age were euthanized by submersion in ice-cold water after being anesthetized with buffered MS222 for 10 minutes. Skin of the trunk and tail was removed and white (fast-twitch) myotomal muscles were dissected and processed. For larval swimming analysis, embryos were reared as above at WUSL until 10 days post-fertilization, assayed for swimming performance, and then euthanized by submersion in ice-cold water for ten minutes after anesthesia.

### Identification of proteins and quantification of their abundances by liquid chromatography mass spectrometry (LCMS)

Rat hindlimb buds, rat tibialis anterior (TA) muscle, 5dpf zebrafish larvae, and adult fast skeletal zebrafish muscle samples were digested with trypsin in preparation for LCMS. Pooled (6–10) prenatal rat hindlimb buds or individual 5dpf zebrafish tails were placed in 1.5 mL microcentrifuge tubes and 150 µL 0.1% RapiGest SF Surfactant (Waters Corporation) was added. Rat TA muscles and fast-twitch adult zebrafish muscles were placed in dissection chambers, the samples were triturated with forceps in 75 µL 0.1% RapiGest then transferred to a 1.5 mL microcentrifuge tube. Each chamber was rinsed with another 75 µL aliquot of 0.1% RapiGest, which was then transferred to the microcentrifuge tube. The microcentrifuge tubes were heated at 50 °C for 45 minutes to 1 hour. Proteins were reduced by addition of 0.75 µL of 1M dithiothreitol to each tube and heating at 100 °C for 10 min. Cysteines were alkylated by addition of 22.5 µL of 100 mM iodoacetamide in 50 mM ammonium bicarbonate and incubation in the dark at 22°C for 30 min. Proteins were digested to peptides by adding 25 µL of 50 mM ammonium bicarbonate containing 5 µg of trypsin (Promega) and incubating at 37 °C for 18 hours. The samples were dried down in a speed vacuum device. Trypsin was deactivated and RapiGest cleaved by addition of 100 µL of 7% formic acid in 50 mM ammonium bicarbonate and heating at 37°C for 1 hour. The resultant peptides were dried down. RapiGest was cleaved again by addition of 100 µL of 0.1% trifluoroacetic acid and heating at 37 °C for 1 hour. The resultant peptides were dried down and reconstituted a final time in 150 µL of 0.1% trifluoroacetic acid. The tubes were centrifuged at 18,800 RCF for 5 minutes (Thermo, Sorvall Legend Micro 21R) to pellet the surfactant. The top 125 µL of solution was transferred into a mass spectrometry (MS) analysis vial.

The resultant peptides were separated by injection of 20 µL of each sample onto an XSelect UPLC HSS T3 column (3.5 μm, 1.0 × 150 mm) (Waters Corporation) attached to a UltiMate 3000 ultra-high pressure liquid chromatography (UHPLC) system (Dionex) as previous described (71). Briefly, the UHPLC effluent was infused into a Q Exactive Hybrid Quadrupole-Orbitrap mass spectrometer through an electrospray ionization source (Thermo Fisher Scientific). Data were collected in data-dependent MS/MS mode with the top five most abundant ions being selected for fragmentation. Peptides were identified from the resultant MS/MS spectra by searching against either a *Rattus norvegicus* (downloaded from UniProt 2/2015) or *Danio rerio* (downloaded from UniProt 2/2015) proteome database using SEQUEST in the Proteome Discoverer 2.2 (PD 2.2) software package (Thermo Fisher Scientific). The potential loss of methionine from the N-terminus of each protein (-131.20 Da), the loss of methionine with addition of acetylation (-89.2 Da), carbamidomethylation (C; 57.02 Da), oxidation (M, P; 16.0 Da: M; 32.00 Da), methylation (K; 14.0d Da), acetylation (K; 42.02 Da), and phosphorylation (S, T, Y; 80.0 Da) were accounted for by variable mass changes. The Minora Feature Detector was used in PD 2.2 to identify LC peaks with the exact mass, charge states, elution time, and isotope pattern as the SEQUEST derived peptide spectral matches (PSMs) across the samples in the entire study.

Label-free quantitative analyses were performed as previously described (62). Briefly, the areas under each LC peak were calculated and reported in PD 2.2 as peptide abundances, and the values were exported to Excel (Microsoft). The abundances of single MyBP or myosin isoforms relative to all MBP or myosin isoforms in each sample were determined from the average abundance of the top 3 ionizing peptides unique to each isoform divided by the summation of the average abundances of the top 3 ionizing peptides from all isoforms within each sample. The relative abundance of MyBP to myosin in the rat samples was determined from the summation of the average abundances of the top 3 ionizing peptides from each MyBP isoform divided by the summation of the average abundances of the top 3 ionizing peptides from each myosin isoform. The relative abundance of MyBP to myosin in the zebrafish samples was determined from the summation of the average abundances of the top 3 ionizing peptides from each MyBP isoform divided by the summation of the average abundances of the top 3 ionizing peptides from myh6 and smyh2 and the average abundance of the top 3 peptides common to all other myosin isoforms. Peptides and quantifications are summarized in Dataset S1.

### Phylogenetic analysis

MyBP protein sequences from human, rat, chicken, and zebrafish were obtained from the National Center for Biotechnology Information at the National Library of Medicine. Accession numbers are noted in File S1. All sequences used are RefSeq curated reference sequences, except for Chicken MYBPC1 (XP_046765984), which was generated from reference genomic sequence (NC_052532.1) by the Gnomon automated prediction program. In genes with multiple predicted or confirmed splice variants, the longest RefSeq sequence isoform was used. The conserved 294-AA region containing the C-terminal-most 3 globular domains used to generate the phylogenetic tree in Figure 1 did not contain any predicted or confirmed variably spliced sequence.

Full-length and homologous C-terminal domain AA sequences were aligned using the MUltiple Sequence Comparison by Log-Expectation (MUSCLE) algorithm (72) in MEGA11 software (Version 11.0.13) (73). Evolutionary analyses of MyBP C-terminal domain sequences were conducted in MEGA11 using the Maximum Likelihood method and JTT matrix-based model (74). The initial tree for the heuristic search was obtained automatically by applying Neighbor-Join and BioNJ algorithms to a matrix of pairwise distances estimated using the JTT model, and then selecting the topology with superior log likelihood value.

### Generation of genetically modified zebrafish

#### Transient expression of FLAG-tagged MyBP sequences

To transiently express tagged MyBP sequence in zebrafish, we used the Tol2 Kit system for transposon-mediated transgenesis as described in (33). Briefly, cDNA from zebrafish *mybpc2b* (NM_001013511) and *mybphb* (NM_001100137) cDNAs were chemically synthesized (BioBasic, Inc., Markham, Ontario, Canada) in frame with sequence encoding 3XFLAG tag (3XAspTyrLysAspAspAspAspLys) (75) and subcloned into the gateway donor vector pDONR221 (Thermo Fisher Scientific, Waltham, MA). These were then combined in an an LR reaction (Multisite Gateway Three Fragment Vector Construction Kit, Thermo Fisher Scientific, Waltham, MA) with tol2kit 5’ entry, 3’ entry, and destination vectors to generate pTol2 containing a 1.5kb promoter element from the zebrafish *hsp70I* gene, *mybpc2b-3XFLAG or mybphb-3XFLAG* sequence, and SV40 polyA (Fig. S1C) (32) (33). The *hsp70I* promoter/enhancer induces transgene expression upon a transient increase in environmental temperature, allowing us to control the timing of expression and avoid interference with early embryogenesis or myogenesis. Embryos were injected at the single-cell stage with pTol2 plasmid DNA and transposase mRNA (33). Injected embryos developed normally to 4dpf at which point they were screened for cardiac eGFP fluorescence from a separate cmlck:eGFP reporter gene included in the construct (Fig. S1A). *Heat shock*: petri dishes containing 4dpf larvae and 50ml E3 medium were transferred to a 37°C incubator for 60 minutes before being returned to 28°C prior to fixation for immunofluorescence studies at 5dpf.

#### Mybphb deletion allele

The null mutation in zebrafish *mybphb* was generated by dual, simultaneous spCas9-mediated double strand breaks at PAM sites in *mybphb* exons 1 and 4, followed by NHEJ and excision of 12kb of intervening sequence, using methods described in(76) (Fig. S1, B). PAM sites and guide RNA sequences were chosen using the ChopChop website (https://chopchop.cbu.uib.no) in knock-out mode searching the latest *Danio rerio* reference assembly (GRCz11) (77, 78). All components of the reaction were purchased from Integrated DNA technologies. Inc. (Coralville, IA), including spCas9 protein, trRNA and chemically synthesized crRNAs (Table S5). 1-cell embryos were injected with a cocktail consisting of precomplexed dgRNPs targeting both sites and allowed to mature to adulthood before being outcrossed to wildtype (AB) strain. PCR was performed on genomic DNA from F1 embryos using primer pairs spanning the predicted ∼12kb deletion (Table S5). Presence of a deletion allele was indicated by the appearance of a ∼200bp amplicon resulting from the repair (Fig. S1B). A single F1 individual carrying a deletion allele was used to establish the colony. Subsequent genotyping was performed by PCR as above, with wildtype alleles identified using an alternative reverse primer located within the deleted sequence (Fig. S1B). The precise sequence of the deletion allele was determined by bi-directional Sanger sequencing of the genotyping PCR amplicon, which confirmed the excision of ∼12kb of intervening DNA, and additionally revealed a +2 frameshift. Elimination of gene expression was confirmed by quantitative LCMS as described, which showed the absence of Mybphb-unique peptides in all homozygous fast myotomal muscle samples.

#### Mybpc2b SA10810 mutants

The SA10810 allele was generated by the Sanger Institute Zebrafish Mutation Project (79) and contains a C>T substitution in *mybpc2b* exon 7 leading to a premature stop codon (Fig S1C). Embryos carrying the SA10810 mutation in *mybpc2b* were acquired from the Zebrafish International Research Consortium (ZIRC, www.zirc.org) and raised to adulthood under standard laboratory rearing and housing conditions. Adults were fin-clipped and genotyped by PCR amplification of the region containing mutation site (Table S5). Amplicons were Sanger sequenced in both directions using the same primers and SA10810-positive carriers were outcrossed to wildtype (AB strain). Heterozygous mating pairs were then crossed to generate homozygous embryos and raised to adulthood. Elimination of gene expression was confirmed by the absence of Mybpc2b-unique peptides in fast myotomal muscle samples assayed by quantitative LCMS as described.

### Imaging of larval zebrafish

#### Fixation and embedding of larval zebrafish tails for brightfield and transmission electron microscopy

For histological examination, 5dpf wildtype and *mybphb^-/-^* sibling larvae from *mybphb ^+/-^* parents were prepared and imaged as described (22). Larvae were euthanized and heads were removed for genotyping. Trunk/tails were fixed by immersion (0.1 M PIPES, 2.5% glutaraldehyde, 1% paraformaldehyde) at room temperature for 1 h, then stored at 4° C. Larvae were then embedded in plastic (80) before sectioning. To visualize myotomal muscle morphology, larvae were oriented to obtain longitudinal or cross sections of muscles, and semithin sections (∼1 mm) were cut with glass knives on a Reichert Ultracut microtome (Leica Biosystems, Wetzlar, Germany), mounted on glass slides, stained with toluidine blue to highlight structural details. Light micrographs were obtained using a Zeiss Axiocam 208 (Carl Zeiss Microscopy, White Plains, NY). To visualize individual sarcomeres, larvae were oriented as above and ultrathin sections (∼80 nm) were cut with a diamond knife. These were retrieved onto 200 mesh nickel grids and contrasted with uranyl acetate (2% in 50% ethanol) and lead citrate. Electron micrographs were obtained with a JEM 1400 transmission electron microscope (JEOL USA, Peabody, MA) operating at 80 kV and a bottom mounted AMT digital camera and software (Advanced Microscopy Techniques, Woburn, MA).

Cross-sectional area (CSA) measurements of whole tails, fast myotomal muscles, and slow myotomal muscles were made by tracing features in 40X light micrographs of toluidine blue stained thick sections. Anatomical landmarks for fast and slow myotomal muscle are well defined (81), which allowed us to outline each muscle type in FIJI/ImageJ software (82) and calculate CSA with the software’s built-in area function. CSA values were measured in sections made immediately cranial to the anal vent, which corresponds to the center of tail sections when mounted for mechanical studies (wildtype n=3, *mybphb^-/-^*n=3).

#### Immunolabeling of F0 zebrafish carrying hsp70i:mybpc2b-3xFLAG or hsp70i:mybpc2b-3xFLAG transgenes

5dpf larvae, injected and heat shocked as described above were euthanized by submersion in tricaine solution (0.05% in E3) for 10 minutes, collected in 1.5 ml Eppendorf tubes and immunolabeled following protocol based on (34) and personal communications with Jared Talbot (University of Maine). Larvae were fixed in PBS containing 4% paraformaldehyde overnight at 4° C, then washed in phosphate buffered saline (PBS) and permeabilized with 10μg/ml proteinase K (Invitrogen, Inc.) in PBST (0.1% Tween-20 in PBS) for 45 minutes. Larvae were then washed in PBST (5x5min), blocked for 2-3 hours with K-block (0.5% TritonX-100, 4% NGS, 2%NSS, 1%DMSO in PBS) and incubated overnight with mouse monoclonal ANTI-FLAG^®^ M2 antibody (Cat. F1804, Sigma-Aldrich, Inc., St Louis, MO) at a concentration of 1:800 in K-block. After washing in PBST (5x5 min), tails were incubated with anti-mouse Igg secondary antibody conjugated to Alexa Fluor^TM^ 488 (Cat. A11001, Life Technologies, Inc., Eugene, OR) at 1:1000 dilution in K-block 2 hours. After final wash in PBST (5x5min), larvae were mounted on slides in Fluorosheild^®^ anti-fade medium (Sigma-Aldrich, St. Louis, MO) for imaging.

#### Immunofluorescence imaging

Immunolabeled larvae were examined with a Nikon A1R-ER Laser Scanning Confocal Microscope system using an Apo TIRF 60x Oil DIC N2 objective and running NIS-Elements software (Nikon Instruments, Tokyo, Japan). Images were acquired at 1024 x 1024 pixels with a resolution of 104 nm/pixel. Laser power, scanning speed and galvanometer detector settings were not changed between samples.

#### Immunofluorescence image analysis and modeling

Analysis of MyBP transgene immunofluorescence staining patterns in fast larval muscle cells followed procedures detailed in (19) with modifications. Briefly, we decomposed confocal immunofluorescence images of anti-FLAG positive fast muscle cells into a set of 1 pixel-wide intensity line scans and then aligned these to each other by maximizing the pairwise cross-correlation. The aligned intensity line scans were normalized for intensity, and then fit with a double Gaussian to determine peak width and peak-to-peak spacing for the fluorescence doublets. Averaged intensity profiles were generated by averaging intensities within aligned line scans in 20nm-wide bins.

The point spread function width was determined as the standard deviation (σ=122 ± 19nm) of a 2-D Gaussian fit to point sources in the background/periphery of confocal images. Intensity profiles were then simulated, as the summed intensity of multiple point sources uniformly distributed within arbitrary boundaries of virtual C-zones equidistant from the sarcomere center. Goodness-of-fit was determined as the root mean squared difference (RMSD) between simulated intensity profiles and the experimental averaged intensity profiles. The model then iteratively adjusted the location of the C-zone boundaries to determine the best fit the experimental data.

### Larval swimming performance

To assay swimming performance, the DanioVision and EthoVision software (Noldus, Wageningen, the Netherlands) were used to record and quantify larval movement at 10dpf as described previously (83). Sibling larvae from heterozygous (*mybphb ^+/-^*) mating pairs were loaded into the DanioVision in 24-well cell culture plates with egg water at random, and acclimated in the DanioVision box for 5-10 min. The culture plate was automatically tapped after acclimation, and the startle response was recorded for 3sec; EthoVision software tracked and recorded fish movement. Average velocity (Fig. 5A) was calculated for each fish as total distance traveled (Fig. 5B) divided by time spent swimming. Larvae were subsequently genotyped by PCR as described above. Statistical analyses were performed only between control (n=48) and *mybphb^-/-^* (n=27) groups assayed on the same day.

### Larval muscle mechanics

Twitch, tetanus, and Force:Velocity measurements in larval tails were performed according to previously published protocols with slight modifications noted here (22). 5dpf sibling larvae from heterozygous (*mybphb ^+/-^* ) mating pairs were selected at random, euthanized by tricaine overdose (0.05% in E3), and transferred to Ringer’s solution (117.2 mM NaCl, 4.7 mM KCl, 1.2 mM KH2PO4, 1.2 mM MgCl2, 2.5 mM CaCl2, 25.2 mM NaHCO3, 11.1 mM glucose; oxygenated and equilibrated to pH 7.4 with a mixture of 95% O2 and 5% C02). Tails were removed by cutting with dissection scissors immediately caudal to the swim bladder, and mounted in a 2-mL Ringer’s-filled bath between a force transducer (400C; Aurora Scientific, Aurora, Ontario, Canada) and linear servo motor/controller (MC1; SI, Heidelberg, Germany) using spring clamps set 1 mm apart. Tails were positioned such that the anal vent was equidistant from the attachment points. Mounted tails were illuminated in bright field by a xenon arc lamp through a fiber optic light guide and a diffuser, which provided the intensity and stability required for high-speed imaging. A charge-coupled device (CCD) camera (IL5, Fastec imaging, San Diego, CA) with custom optics and a resolution of 0.95 µm/pixel at 5000 frames/s, was used to record the preparation throughout each experimental run. For all experiments, servo motor position, stimulation timing, and camera triggering was preprogramed using custom-made control and data acquisition software written in IGOR (WaveMetrics, Portland, OR) running on a desktop PC and controlled via an A/D board (PCle6251; National Instruments, Austin, TX). Force, motor position, and camera timing pulse were sampled at 40 kHz and saved to a hard disk. Each experimental run was triggered by a signal from the camera corresponding to the first acquired image, which simplified the synchronization of the various signals.

After attachment, the length of each preparation was adjusted by moving the position of the force transducer using a calibrated micromanipulator to remove any strain imposed during attachment, and thus returning tail sections to their *in vivo* length (L0). Preparations were then further stretched to 1.05 X L0 for mechanical studies. The temperature of the preparation was maintained at 23° C by fresh, oxygenated Ringer’s, pumped at a constant velocity (0.7 mL/min or 6 exchanges/min) through the experimental chamber after being heated or cooled in a custom countercurrent heat exchanger. Tails were then left to rest for 20 min. prior to stimulation. Twitches were evoked with a single 0.4msec, 7V current pulse via a MyoPacer stimulator (IonOptix, Inc. Westwood, MA) using the attachment clamps as electrodes. Tetanic contractions were evoked with a 100msec long train of 0.4msec 7V current pulses at 300 Hz.

To measure the effect of shortening velocity on active force during steady-state activation, preparations were stretched by servo control at 0.5 preparation lengths/second to 1.15 x L0, before being tetanized (100 ms, 300 ) (Fig. 5G). At 40 ms after the initiation of the tetanus, preparations were shortened by servo control from 1.15–1.05 x L0 at a constant velocity. Under these conditions, we were limited to preparation shortening velocities less than 20 lengths/s by the resolution of the equipment. The same protocol was then repeated without stimulation to measure the tail’s passive force response to the length changes. The passive force trace was subtracted from the stimulated force trace to determine active force. The proportion of maximal active isometric force developed, whereas shortening at a particular velocity was defined as the minimal force achieved before the end of the shortening ramp divided by the active force measured 20 ms after the end of the ramp, at which point the recovery of isometric force had reached a plateau. Myotome length was measured throughout the experiment as described in (22), and myotome shortening velocity was calculated as the proportional change in myotome length over the duration of the ramp. Stimulated and passive shortening runs were repeated in the same preparation at multiple imposed shortening velocities: 2, 5, 10, 20 preparation lengths/second.

### Small angle X-ray diffraction

X-ray diffraction measurements were performed at the Life Science X-ray Scattering (LiX) beamline at Brookhaven National Laboratories (BNL) in Upton, NY, USA (84). The proposed use of zebrafish larvae was reviewed and approved by the Institutional Animal Care and Use Committees of UVM and BNL. Five-day old larvae were randomly selected, euthanized by submersion in tricaine (0.05%) for 10 minutes, and dissected under a dissection microscope. Heads were removed and flash-frozen for genotyping, and ∼1mm length of the tail, centered on the anal vent, was clamped between two stainless steel clamps housed in a 1 mm wide space bordered by two removable, mylar windows. The clamps were connected by tungsten wire to external leads that permitted electrical stimulation to the tail muscle (MyoPacer, IonOptix, Inc. Westwood, MA). X-ray reflection patterns were recorded for three conditions: 1) resting pre-stretch (L0) with expected 1.86 um sarcomere length (SL), 2) resting after 10% stretch (L10) with ∼2.0 um SL, and 3) tetanized at L10 with 300 Hz electrical field stimulation (7V, 0.4msec) using the clamps as electrodes.

X-ray diffraction patterns were collected according to (84) based on a small-angle X-ray scattering (SAXS) detector (Pilatus 1M) located at the end of a 3.56 m flight path. X-ray energy was 15.155 keV with photon flux of ∼1.1⨯10^12^ photons/s at the sample. Beam size at the sample was ∼50 x 50 um^2^ FWHM. Any one position on the sample was irradiated with five 100 ms exposures to allow rejection of exposures that did not contribute to overall signa-to-noise ratio (SNR). Using SNR defined as fitted peak intensity / (peak + background intensity), we found that all five exposures consistently contributed to defining the signal. To minimize radiation damage, additional positions were selected at least 100 µm away from any previous position.

Images included equatorial reflections indicative of d1,0 and d1,1 spacing (Fig. 4A). The crescent shape of these reflections arose from the natural heterogeneous orientation of sarcomeres within the tail muscle. The total reflection intensity as a function of radius, T(r), from the origin in reciprocal space was represented by integrating over the crescent shape for d1,0 between the angles corresponding to 50% maximum intensity such that T(r) = ʃ I(r, f) r df. Background intensity was similarly calculated as a function of radius, B(r), and was taken from the average of background angles on either side of the d1,0 reflection. The resulting reflection intensity Int(r)=T(r)-B(r) was then fit to two Gaussian distributions with the radius for d1,1 set to √3⨯radius of d1,0 and a second-order polynomial representation of background (Fig. 4C). Larval heads were subsequently genotyped, (wildtype n=7, *mybphb^-/-^* n=8).

### In vitro motility

#### Isolation of native myosin thick filaments

Filaments were isolated from myofibrils prepared from adult fast myotomal muscle samples. Briefly, fast (white) myotomal muscle samples were dissected from adult zebrafish and immediately washed in in Rigor buffer (100 mM Na acetate, 8mM MgCl2, 1 mM EGTA, 2 mM imidazole, 10 mM phosphocreatine, pH 6.8) with protease inhibitors (PMSF, Pepstatin and Aprotinin, 0.004 mg/mL each). Tissue was homogenized by two rounds (∼10s each) at 21k RPM in a Tissue Tearor (Biospec Products, Bartlesville, OK), and then chemically skinned in Rigor buffer containing 0.5% (v/v) Triton X-100 for 30 min with agitation. Myofibrils were washed 3X by centrifugation at 1000xg for 5 min, followed by resuspension of the pellet in 1mL of Relaxing buffer (Rigor Buffer with 5mM ATP) with protease inhibitors. Native thick filaments were then released from the myofibrils by enzymatic digestion with the addition of Calpain-1 from porcine erythrocytes (CALBIOCHEM, 0.1U/μl, final concentration) and Calcium Acetate (2mM final concentration) for 10 minutes at room temperature. Undigested muscle was pelleted by centrifugation at 3000rpm for 3 minutes, leaving a supernatant containing isolated thick filaments. Preparation quality (monodisperse, intact filaments) was verified by visualization in “Ratiometric Mode” on a REFEYN OneMP mass photometer (Oxford, UK).

#### Native thick filament motility

Motility assays were performed as detailed previously (26). Sigmacote (Sigma-Aldrich, St. Louis, MO) treated flow cells were incubated with freshly prepared native thick filaments and allowed to incubate at room temperature for 20 min. The flow cell surface was then blocked with BSA (1 mg/mL), followed by 1μM F-actin from chicken pectoralis to block non-function myosin “dead heads.” Finally, motility buffer (25 mM KCl, 1 mM EGTA, 10 mM DTT, 25 mM imidazole (pH 7.4), 4 mM MgCl2, 100 μM ATP) with an oxygen scavenging system (0.1 μg/mL glucose oxidase, 0.018 μg/mL catalase, 2.3 μg/mL glucose) with freshly sonicated rhodamine-phalloidin labeled actin filaments were introduced into the flow cell. The motion of the actin was captured under total internal reflection fluorescence imaging at 100 fps on a Nikon TE2000U microscope with PlanApo objective (100x, 1.49 n.a.), equipped with an XR Turbo G intensified CCD camera (Stanford Photonics, Palo Alto CA).

#### Analysis of motility data

Movies were analyzed as described previously (19, 26), with slight modifications. Trajectories were identified visually in FIJI/ImageJ (81), followed by tracking with sub-pixel accuracy using SpotTracker2D. Linear trajectories spanning <800nm were selected for further analysis, with displacement determined as Pythagorean distance from the filament’s initial position. Biphasic velocities were determined by fitting Displacement vs Time trajectories with 2 straight line segments using the segmented model regression function, implemented in the “Segmented” library for statistical programming language R. Trajectories were classified as a “single phase” if either segment lasted <50ms (5 frames), the velocity difference between the two segments <10%, or if the velocity of either segment is zero (p>0.05).

### Statistics and blinding

All sample preparations, experiments, and analysis were performed with the investigator blinded to genotype. Data are reported as means ± 1 standard deviation. Statistics were performed using GraphPad Prism 10 software (GraphPad Software, Boston, MA). For pairwise comparisons between wildtype and mutant samples, dependent variables were compared using two-tailed, unpaired Student’s t-tests (protein abundance, larval muscle swimming performance, intact tail mechanics, CSA measurements, *in vitro* motility). A two-way analysis of variance (ANOVA) design with fixed effects was used to compare the effect of stretch or tetanic stimulation on lattice spacing in X-ray studies of wildtype and mutant zebrafish larvae (treatment, genotype, treatment x genotype). Additional statistical methods are described in the sections above where appropriate.

## Acknowledgments

We thank Todd Clason (Imaging/Physiology Core Facility, University of Vermont) and Tim Connolly (Microscopy Imaging Center, University of Vermont) for expert assistance in tissue sectioning and transmission electron microscopy. We thank Jared Talbot (University of Maine) and Filip Braet (University of Sydney) for expert advice in zebrafish immunolabeling. We thank Ashley Waldron (University of Vermont, AddGene) for expert assistance in zebrafish genetic approaches. We thank Sebastian Duno-Miranda (University of Vermont) and Brandon Bensel (University of Vermont) for insightful conversations. Imaging work was performed at the Microscopy Imaging Center at the University of Vermont (RRID# SCR_018821). Confocal microscopy was performed on a Nikon A1R-HD point scanning confocal supported by NIH award number S10OD025030. The LiX beamline is part of the Center for BioMolecular Structure (CBMS), which is primarily supported by the NIH (P30GM133893), and by the DOE Office of Biological and Environmental Research (KP1605010). LiX also received additional support from NIH (S10OD012331). As part of NSLS-II, a national user facility at Brookhaven National Laboratory, work performed at the CBMS is supported in part by the U.S. Department of Energy, Office of Science, Office of Basic Energy Sciences Program under contract number DE-SC0012704. The REFEYN OneMP mass photometer was purchased from funds from NIH award number R35GM141743-03S1.

This work was supported by funds from the NIH to D.M.W. (R01HL150953), to M.J.P (R01HL157487), to C.A.G. (P50HD103525 and R01AR067715), to M.J.C. (NS120419-01A1), and a generous gift from Arnold and Mariel Goran to D.M.W..

## Glossary of protein and gene names

MyBP-H: Myosin-binding protein H, generic term
fMyBP-C: Fast myosin-binding protein C, generic term
sMyBP-C: Slow myosin-binding protein C, generic term
cMyBP-C: Cardiac myosin-binding protien C, generic term
MyBP: Myosin-binding protein C/H family member, generic term
*MYBPH*: Human MyBP-H gene
*MYBPC1*: Human sMyBP-C gene
*MYBPC2*: Human fMyBP-C gene
*MYBPC3*: Human cMyBP-C gene
Mybpha: Zebrafish MyBP-H isoform a; *mybpha* protein product
Mybphb: Zebrafish MyBP-H isoform b; *mybphb* protein product
Mybpc2a: Zebrafish fMyBP-C isoform a; *mybpc2a* protein product
Mybpc2b: Zebrafish fMyBP-C isoform b; *mybphc2b* protein product
*mybpha*: Zebrafish MyBP-H gene (little to no expression in skeletal muscle)
*mybphb*: Zebrafish MyBP-H gene (highly expressed in skeletal muscle)
*mybpc2a*: Zebrafish fMyBP-C gene (little to no expression in skeletal muscle)
*mybpc2b*: Zebrafish fMyBP-C gene (highly expressed in skeletal muscle)
*mybpc3*: Zebrafish cMyBP-C gene (only expressed in cardiac muscle)

