## Appendix: Figure S1, Tables S1-5 for "Functional role of myosin-binding protein H in thick filaments of developing vertebrate fast-twitch skeletal muscle"

[a]Department of Molecular Physiology and Biophysics, Larner College of Medicine, University of Vermont, Burlington, VT 05405; [b]Cardiovascular Research Institute, University of Vermont, Burlington, VT 05405; [c]National Synchrotron Light Source II, Brookhaven National Laboratory, Upton, NY 11973; [d]Department of Neurlogical Sciences, Larner College of Medicine, University of Vermont, Burlington, VT 05405; [e]Department of Electrical and Biomedical Engineering, College of Engineering and Mathematical Sciences, University of Vermont, Burlington, VT 05405; [f]Department of Neurology, Washington University School of Medicine in St. Louis, St. Louis, MO 63110; [g]Department of Biology, College of Arts and Sciences, University of Vermont, Burlington, VT 05405; [h]Department of Developmental Biology, Washington University School of Medicine in St. Louis, St. Louis, MO 63110

Corresponding authors: David M. Warshaw, Michael J. Previs

**This PDF file includes:**

Figure S1
Tables S1 to S5
Legend for file S1
Legend for Datasets S1

**Other supporting materials for this manuscript include the following:**

File S1
Dataset S1

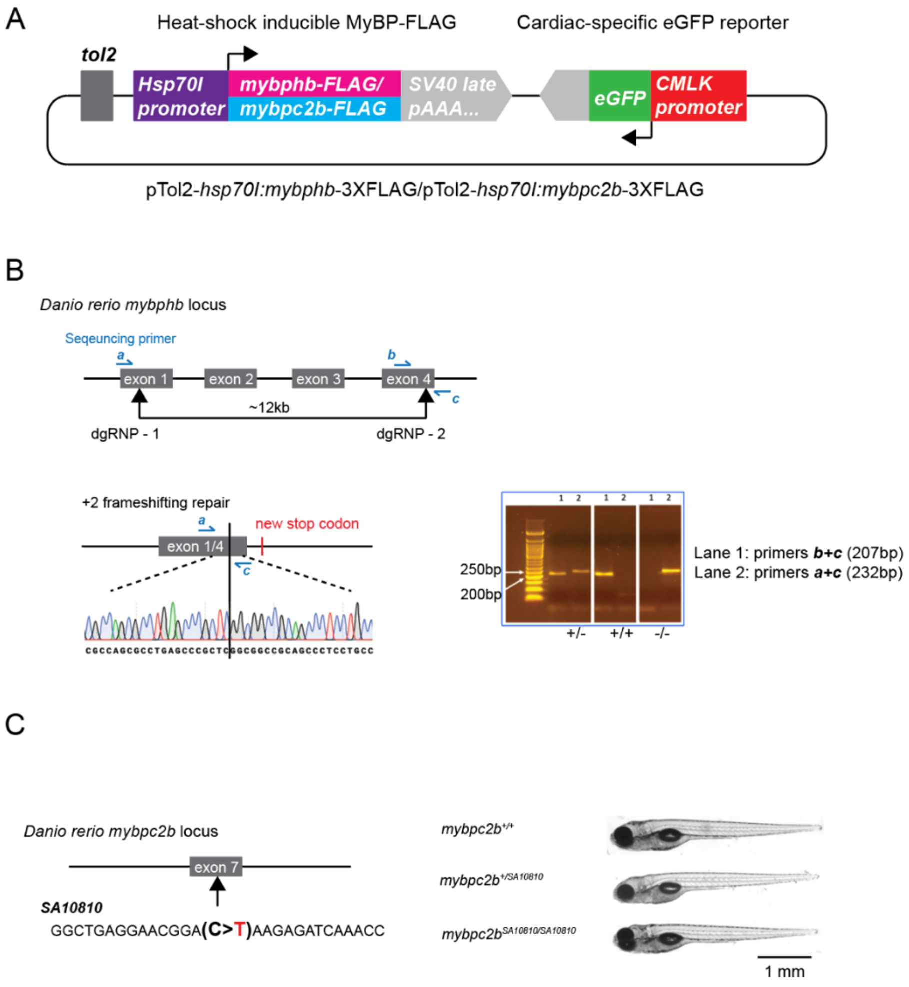


**Fig. S1.** Genetically modified zebrafish. (A) Transient mosaic expression of FLAG-tagged MyBP was accomplished using the modular 'tol2kit' approach for assembly of transgene constructs and transposase-mediated integration into gDNA. *pTol2-Hsp70l:mybphb-3XFLAG* and *pTol2-hsp70l:mybpc2b-3XFLAG* plasmids were generated by cloning of chemically synthesized transgene cDNA along with hsp70l promoter and SV40 late poly-A sequences into a backbone vector containing tol2 ITR sequence and a cardiac-specific eGFP reporter cassette as described in Kwan et al. 2004. (B) The *mybphb* null allele (Figs. 3-5) was generated by simultaneous CRISPR/Cas9 mediated double strand breaks by spCas9 protein precomplexed to guide RNA targeting sites in exons 1 and 4 (see Table S5), followed by cell-mediated repair and excision of the intervening ~12kb of genomic DNA. Genotyping was accomplished by PCR using primers a, b, c, as shown (see Table S5). Precise characterization of the mutant allele was accomplished by PCR followed by Sanger sequencing. (C) The SA10810 mutation in mybpc2b (Fig. 6) consists of a single base substitution in exon 7 creating a premature STOP codon. Genotyping was accomplished by PCR and bidirectional Sanger sequencing of the mutation site (see Table S5).

| | | | e14 Limb Bud | | e16 Limb Bud | | e18 TA | | e20 TA | | adult TA | |
|---|---|---|---|---|---|---|---|---|---|---|---|---|
| Gene | NCBI Gene ID | Protein name | abundance (relative to total MYH) | SD | abundance (relative to total MYH) | SD | abundance (relative to total MYH) | SD | abundance (relative to total MYH) | SD | abundance (relative to total MYH) | SD |
| *MYH1* | 287408 | Fast IIa | -- | -- | -- | -- | -- | -- | -- | -- | 0.070 | 0.050 |
| *MYH2* | 691644 | Fast IIb | -- | -- | -- | -- | -- | -- | -- | -- | 0.003 | 0.004 |
| *MYH4* | 360543 | Fast IIx | -- | -- | -- | -- | -- | -- | -- | -- | 0.927 | 0.054 |
| *MYH7* | 29557 | Slow/Beta-cardiac | -- | -- | 0.018 | 0.001 | 0.080 | 0.004 | 0.060 | 0.004 | -- | -- |
| *MYH3* | 24583 | Embryonic | -- | -- | 0.322 | 0.024 | 0.794 | 0.005 | 0.747 | 0.003 | -- | -- |
| *MYH8* | 252942 | Neonatal | -- | -- | -- | -- | 0.093 | 0.002 | 0.179 | 0.008 | -- | -- |
| *MYH9* | 25745 | NMIIA | 0.526 | 0.056 | 0.252 | 0.011 | 0.033 | 0.008 | 0.015 | 0.002 | -- | -- |
| *MYH10* | 79433 | NMIIB | 0.474 | 0.056 | 0.408 | 0.014 | -- | -- | -- | -- | -- | -- |

**Table S1.** Myosin heavy chain isoform abundance relative to total myosin heavy chain in rat hindlimb bud and TA.

| Gene | NCBI Gene ID | Protein name | e14 Limb Bud abundance (relative to sarcomeric MYH) | e14 Limb Bud SD | e16 Limb Bud abundance (relative to sarcomeric MYH) | e16 Limb Bud SD | e18 TA abundance (relative to sarcomeric MYH) | e18 TA SD | e20 TA abundance (relative to sarcomeric MYH) | e20 TA SD | adult TA abundance (relative to sarcomeric MYH) | adult TA SD |
|---|---|---|---|---|---|---|---|---|---|---|---|---|
| *MYBPC1* | 362867 | sMyBP-C | -- | -- | -- | -- | 0.050 | 0.003 | 0.065 | 0.002 | 0.035 | 0.005 |
| *MYBPC2* | 292879 | fMyBP-C | -- | -- | -- | -- | -- | -- | -- | -- | 0.057 | 0.002 |
| *MYBPH* | 83708 | MyBP-H | -- | -- | -- | -- | 0.054 | 0.003 | 0.026 | 0.002 | 0.007 | 0.002 |

**Table S2.** Myosin binding protein isoform abundance relative to total sarcomeric myosin heavy chain in rat hindlimb bud and TA.

| | | 5dpf tail Wildtype (n=5) | | 5dpf tail *mybphb+/-* (n=3) | | 5dpf tail *mybphb-/-* (n=4) | | Adult Fast Wildtype (n=3) | | Adult Fast *mybpc2b-/-* (n=3) | |
|---|---|---|---|---|---|---|---|---|---|---|---|
| Gene | NCBI Gene ID | abundance (relative to total MYH) | SD | abundance (relative to total MYH) | SD | abundance (relative to total MYH) | SD | abundance (relative to total MYH) | SD | abundance (relative to total MYH) | SD |
| *myhc4* | 334274 | 0.005 | 0.004 | 0.023 | 0.022 | 0.009 | 0.009 | 0.629 | 0.048 | 0.615 | 0.017 |
| *myhz1.3* | 100008070 | 0.025 | 0.004 | 0.046 | 0.010 | 0.039 | 0.007 | 0.335 | 0.048 | 0.335 | 0.001 |
| *myhz1.1* | 58142 | 0.542 | 0.037 | 0.499 | 0.064 | 0.505 | 0.038 | -- | -- | -- | -- |
| *myhz2* | 246275 | 0.304 | 0.026 | 0.300 | 0.016 | 0.320 | 0.026 | -- | -- | -- | -- |
| *myha* | 100149148 | 0.005 | 0.002 | 0.009 | 0.004 | 0.007 | 0.003 | 0.023 | 0.011 | 0.037 | 0.020 |
| *myh6* | 386711 | 0.011 | 0.003 | 0.011 | 0.001 | 0.013 | 0.003 | 0.006 | 0.002 | 0.007 | 0.002 |
| *smyhc1* | 407636 | 0.104 | 0.018 | 0.105 | 0.020 | 0.101 | 0.012 | -- | -- | -- | -- |
| *smyhc2* | 100124599 | 0.005 | 0.001 | 0.007 | 0.004 | 0.007 | 0.002 | 0.006 | 0.002 | 0.006 | 0.001 |

**Table S3.** Myosin heavy chain isoform abundance relative to total myosin heavy chain in zebrafish.

| | | | 5dpf tail Wildtype (n=5) | | 5dpf tail *mybphb+/-* (n=3) | | 5dpf tail *mybphb-/-* (n=4) | | Adult Fast Wildtype (n=3) | | Adult Fast *mybpc2b-/-* (n=3) | |
|---|---|---|---|---|---|---|---|---|---|---|---|---|
| Gene | NCBI Gene ID | Protein name | abundance (relative to total MYH) | SD | abundance (relative to total MYH) | SD | abundance (relative to total MYH) | SD | abundance (relative to total MYH) | SD | abundance (relative to total MYH) | SD |
| *mybpc2b* | 541384 | Mybpc2b | 0.005 | 0.000 | 0.007 | 0.002 | 0.006 | 0.001 | 0.030 | 0.004 | -- | -- |
| *mybpha* | 393530 | Mybpha | -- | -- | -- | -- | -- | -- | 0.001 | 0.000 | 0.001 | 0.000 |
| *mybphb* | 321053 | Mybphb | 0.077 | 0.008 | 0.050 | 0.003 | 0.000 | 0.000 | 0.074 | 0.009 | 0.104 | 0.005 |

**Table S4.** Myosin binding protein isoform abundance relative to total myosin heavy chain in zebrafish.

| | name | sequence (5'->3') | source |
|---|---|---|---|
| *mybphb deletion genotyping primers (see Fig. S1B)* | "a" (forward) | AACCAGCACCGATAAAGAAGTC | IDT, Coralville, IA |
| | "b" (forward, deletion) | GTGTCCTGCCGTTATGTGATTA | IDT, Coralville, IA |
| | "c" (reverse) | TCCTTCAAAGAATCCTGTTTCC | IDT, Coralville, IA |
| *mybpc2b SA10810 genotyping primers (see Fig. S1C)* | forward | TGTAAATGTGGGATGTTTCTGC | IDT, Coralville, IA |
| | reverse | AGCATTTCTCATCCATTTCACA | IDT, Coralville, IA |
| *mybphb CRISPR/Cas9 target sequence for crRNA generation (see Fig S1A)* | exon 1 site (-strand) | CGGTGCGGGTTCGGCTGGAGCGG | IDT, Coralville, IA |
| | exon 4 site (+strand) | GTCGTGGCCATGAACCCCGGCGG | IDT, Coralville, IA |

**Table S5.** Genotyping primers and CRISPR/Cas9 genomic target sequences.

**File S1 (separate file).** MUSCLE alignment of full-length myosin binding protein isoforms from human, rat, chicken, and zebrafish reference sequences. Accession numbers are included for each aligned sequence. Green boxes surrounding the conserved proline at human sMyBP-C position 844 and the conserved cysteine at human sMyBP-C position 1139 represent the boundaries of 3 globular domains shared among all isoforms. An alignment of the intervening sequence was used to generate the tree in Fig. 1.

**Dataset S1 (separate file).** Summary of LCMS data. A spreadsheet with peptides identified by LCMS from myosin heavy chain and myosin binding protein isoforms in 5dpf zebrafish tails, adult zebrafish fast myotomal muscle, rat hindlimb bud and rat TA muscle, and their abundances.
