## Supplementary material for "Functional role of myosin-binding protein H in thick filaments of developing vertebrate fast-twitch skeletal muscle": File S1

```
                                                                                        20                   40
                                                                                        |                    |
XP 046765984 myosin-binding protein C slow-type Gallus gallus             ----PEPTKK DETETENALP PPEQKTPQDG ---------- ---------- 26
NP 001038124 myosin-binding protein C fast-type Gallus gallus             ---MPEPSKA APKKEAKK-- ---------- ---------- ---------- 15
NP 001384345 myosin-binding protein C cardiac-type Gallus gallus          ---MPEPAKK AVSAFTKKPK TTEVAAGSTA VFEAETEKTG IKVKWQRAGT 47
NP 001026199 myosin-binding protein H Gallus gallus                       ---MTGKTAP AAA------- ---------- ---------- ---------- 10
NP 002456 myosin-binding protein C slow-type Homo sapiens                 ---MPEPTKK EENEVPAPAP PPE------- ---------- ---------- 20
NP 004524 myosin-binding protein C fast-type Homo sapiens                 ---MPE-AKP AAKKAPKGKD AP-------- ---------- ---------- 18
NP 000247 myosin-binding protein C cardiac-type Homo sapiens              ---MPEPGKK PVSAFSKKPR SVEVAAGSPA VFEAETERAG VKVRWQRGGS 47
NP 004988 myosin-binding protein H Homo sapiens                           ---MMEKNTS EGPACS---- ---------- ---------- ---------- 13
NP 001010985 myosin-binding protein H-like Homo sapiens                   ---MEAATAP EVAAGSKL-- ---------- ---------- ---------- 15
NP 001094228 myosin-binding protein C slow-type Rattus norvegicus         ---MPEPTKK EENEVPAPAP PPE------- ---------- ---------- 20
NP 001381991 myosin-binding protein C fast-type Rattus norvegicus         ---MPE-AKP AAKKAPKGKD AP-------- ---------- ---------- 18
NP 001099960 myosin-binding protein C cardiac-type Rattus norvegicus      ---MPEPGKR PVSAFTKKPR SVEVTAGSAA VFEAETERSG LKVQWQRDGS 47
NP 114001 myosin-binding protein H Rattus norvegicus                      ---MTGKATP EASVSTSE-- ---------- ---------- ---------- 15
NP 001014064 myosin-binding protein H-like Rattus norvegicus              ---METATTL EIASCSQR-- ---------- ---------- ---------- 15
NP 001007323 myosin-binding protein C slow-type Danio rerio               ---MPEPTKK DDG------- ---------- ---------- ---------- 10
NP 001039324 myosin binding protein C2a Danio rerio                       ---MPEPKKP AAKEEA---- ---------- ---------- ---------- 13
NP 001013529 myosin binding protein C2b Danio rerio                       ---MPEPTEA EVK------- ---------- ---------- ---------- 10
NP 001037814 myosin-binding protein C cardiac-type Danio rerio            ---MPEPTKK IVSAFSKKPK SQTAEIGARV IFEAETEKPG VKVKWQRESK 47
NP 956852 myosin binding protein Ha Danio rerio .                         MPAKPAPIKK SAKKAAEK-- ---------- ---------- ---------- 18
NP 001093607 myosin binding protein Hb Danio rerio                        MPAKPAPIKK SAKAAKKE-- ---------- ---------- ---------- 18
Consensus                                                                 ---MPEPTKK EAKAXSKK-- ---------- ---------- ----------
```

Conservation (100% – 0%)

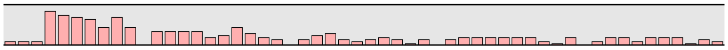

```
                                                                                   60                   80                   100
                                                                                   |                    |                    |
XP 046765984 myosin-binding protein C slow-type Gallus gallus             ---------- ---------- ---------- ---------- ---------- 26
NP 001038124 myosin-binding protein C fast-type Gallus gallus             ---------- ---------- ---------- ---------- ---------- 15
NP 001384345 myosin-binding protein C cardiac-type Gallus gallus          EITDSEKYAI KAEGNKHSLT ISNVGKDDEV TYAVIAGTSK VKFEL----- 92
NP 001026199 myosin-binding protein H Gallus gallus                       ---------- ---------- ---------- ---------- ---------- 10
NP 002456 myosin-binding protein C slow-type Homo sapiens                 ---------- ---------- ---------- ---------- ---------- 20
NP 004524 myosin-binding protein C fast-type Homo sapiens                 ---------- ---------- ---------- ---------- ---------- 18
NP 000247 myosin-binding protein C cardiac-type Homo sapiens              DISASNKYGL ATEGTRHTLT VREVGPADQG SYAVIAGSSK VKFDLKVIE- 96
NP 004988 myosin-binding protein H Homo sapiens                           ---------- ---------- ---------- ---------- ---------- 13
NP 001010985 myosin-binding protein H-like Homo sapiens                   ---------- ---------- ---------- ---------- ---------- 15
NP 001094228 myosin-binding protein C slow-type Rattus norvegicus         ---------- ---------- ---------- ---------- ---------- 20
NP 001381991 myosin-binding protein C fast-type Rattus norvegicus         ---------- ---------- ---------- ---------- ---------- 18
NP 001099960 myosin-binding protein C cardiac-type Rattus norvegicus      DIAANDKYGL AAEGKRHTLT VRDVGPDDQG SYAVIAGSSK VKFDLKVTE- 96
NP 114001 myosin-binding protein H Rattus norvegicus                      ---------- ---------- ---------- ---------- ---------- 15
NP 001014064 myosin-binding protein H-like Rattus norvegicus              ---------- ---------- ---------- ---------- ---------- 15
NP 001007323 myosin-binding protein C slow-type Danio rerio               ---------- ---------- ---------- ---------- ---------- 10
NP 001039324 myosin binding protein C2a Danio rerio                       ---------- ---------- ---------- ---------- ---------- 13
NP 001013529 myosin binding protein C2b Danio rerio                       ---------- ---------- ---------- ---------- ---------- 10
NP 001037814 myosin-binding protein C cardiac-type Danio rerio            DILPSHKYTI SAEDNKHSLT INNIIQEDAV GYAVIAGMSK VKFELKIKES 97
NP 956852 myosin binding protein Ha Danio rerio .                         ---------- ---------- ---------- ---------- ---------- 18
NP 001093607 myosin binding protein Hb Danio rerio                        ---------- ---------- ---------- ---------- ---------- 18
Consensus                                                                 ---------- ---------- ---------- ---------- ----------
```

Conservation (100% – 0%)

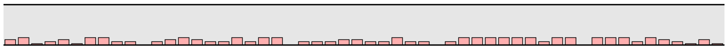

```
                                                                                             120                  140
                                                                                             |                    |
XP 046765984 myosin-binding protein C slow-type Gallus gallus             --------GT EKGKDMDVRE MASSSDEDDT SVSTPPADGM SKPSERKDSV 68
NP 001038124 myosin-binding protein C fast-type Gallus gallus             --------KE EKKEEKKEAP PPQEHKDEAP DDVHPP---- ---------- 43
NP 001384345 myosin-binding protein C cardiac-type Gallus gallus          --------KV KEPEKSEPVA PAEASPAPAA SELPAP---- ---------P 121
NP 001026199 myosin-binding protein H Gallus gallus                       --------KK APAAKKAPAP ASKKAPEPAP KEKPAP---- ---------- 38
NP 002456 myosin-binding protein C slow-type Homo sapiens                 --------EP SKEKEAGTTP AKD-EEEVSP PSALPPGLG- SRALERKDSD 60
NP 004524 myosin-binding protein C fast-type Homo sapiens                 --------KG APKEAPPKEA PAE-APKEAP PEDQSP---- ---------- 45
NP 000247 myosin-binding protein C cardiac-type Homo sapiens              --------AE KAEPMLAPAP APA-EATGAP GEAPAPAAEL GESAPSPKGS 137
NP 004988 myosin-binding protein H Homo sapiens                           ---------- ---------- ----PEETAS ESAKVPTA-- ---------- 27
NP 001010985 myosin-binding protein H-like Homo sapiens                   ---------- ---------- ----KVKEAS PADAEP---- ---------- 27
NP 001094228 myosin-binding protein C slow-type Rattus norvegicus         --------EP NKDKEAGTTP AKE-EDEASP PSALPPGLG- SRALERKDSE 60
NP 001381991 myosin-binding protein C fast-type Rattus norvegicus         --------KE AP--AAKQTP PAE-PPKEAP PEDQSP---- ---------- 43
NP 001099960 myosin-binding protein C cardiac-type Rattus norvegicus      --------PA PPEKAESAVA PTSMEAPETP KEVPALATQL EGNVSSPEGS 138
NP 114001 myosin-binding protein H Rattus norvegicus                      ---------- ---------- ----GTAPEP AKVPTP---- ---------- 27
NP 001014064 myosin-binding protein H-like Rattus norvegicus              ---------- ---------- --QVEEAADP ADAEG----- ---------- 28
NP 001007323 myosin-binding protein C slow-type Danio rerio               --------QP EEDGSTPKKL SIDLPNDSDP ESTRGR---- ------KDSV 42
NP 001039324 myosin binding protein C2a Danio rerio                       ---------- ---------- ----PPADKP AEEHVP---- ---------- 25
NP 001013529 myosin binding protein C2b Danio rerio                       ---------- ---------- ----PEEEAP GDDVEP---- ---------- 22
NP 001037814 myosin-binding protein C cardiac-type Danio rerio            EVKGVEKAKS PDPAAVEHKP EESKPEESKP EPKEPPPAPT ENSAPESTGP 147
NP 956852 myosin binding protein Ha Danio rerio .                         ---------- -EEKPAAPAP VPETAPVEAP PAEDPP---- ---------- 43
NP 001093607 myosin binding protein Hb Danio rerio                        ---------- ------EPAP APAPEPEPAP APEPAP---- ---------- 38
Consensus                                                                 ---------- ---------P --E-XPEEAP PEAPPP---- ----------
```

Conservation (100% – 0%)

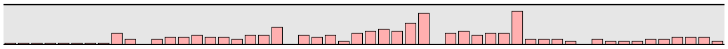

```
                                                                        160                           180                           200
                                                                          |                             |                             |
XP 046765984 myosin-binding protein C slow-type Gallus gallus          WSIGETAP-- -EEAEKRDDS QRSTLFIEKP QSGTVSVGGN ITFIAKVEAK 115
NP 001038124 myosin-binding protein C fast-type Gallus gallus          ---------- -------ETP DPEGLFLSKP QNVMVESGRD VTVSARVAGA 76
NP 001384345 myosin-binding protein C cardiac-type Gallus gallus       VESNQNPEVP PAETQPEEPV DPIGLFVTRP QDGEVTVGGN ITFTAKVAGE 171
NP 001026199 myosin-binding protein H Gallus gallus                    ---------- ---------- --------TP KEGHA----- ---------- 45
NP 002456 myosin-binding protein C slow-type Homo sapiens              WTLVETPP-G EEQAKQNANS QLSILFIEKP QGGTVKVGED ITFIAKVKAE 109
NP 004524 myosin-binding protein C fast-type Homo sapiens              ---------- -------TAE EPTGVFLKKP DSVSVETGKD AVVVAKVNGK 78
NP 000247 myosin-binding protein C cardiac-type Homo sapiens           SSAALNGP-- ----TPGAPD DPIGLFVMRP QDGEVTVGGS ITFSARVAGA 181
NP 004988 myosin-binding protein H Homo sapiens                        ---------- ---------- ---------- ---------- ---------- 27
NP 001010985 myosin-binding protein H-like Homo sapiens                ---------- ---------- ---------- ---------- ---------- 27
NP 001094228 myosin-binding protein C slow-type Rattus norvegicus      WSLGESPAGG EEQDKQNANS QLSTLFVEKP QTGSVKVGAN ITFIAKVKAE 110
NP 001381991 myosin-binding protein C fast-type Rattus norvegicus      ---------- -------TVE EPTGLFLKKP DSVSVETGKD TVILAKVNGK 76
NP 001099960 myosin-binding protein C cardiac-type Rattus norvegicus   VSVTQDGS-- -VAGSQGAPD DPIGLFLMRP QDGEVTVGGS IVFSARVAGA 185
NP 114001 myosin-binding protein H Rattus norvegicus                   ---------- ---------- ---------- ---------- ---------- 27
NP 001014064 myosin-binding protein H-like Rattus norvegicus           ---------- ---------- ---------- ---------- ---------- 28
NP 001007323 myosin-binding protein C slow-type Danio rerio            WSLGDGQP-P EEIDKQTENP PLSTLLIEKP QSGSITVGGD ITFIAKVEAK 91
NP 001039324 myosin binding protein C2a Danio rerio                     ---------- ---GEESQMS EISGLFVERP QTVAAIKGKD VTFVAKVDSS 62
NP 001013529 myosin binding protein C2b Danio rerio                     ---------- ---------T ELTGLFVEKP ESVVAIAGKD VTFVVKVDST 53
NP 001037814 myosin-binding protein C cardiac-type Danio rerio         PSDGVAPEAL PESHQPDTRQ DLTGLFTEKP HSGEVNVGEN IIFIARVCGE 197
NP 956852 myosin binding protein Ha Danio rerio .                      ---------- ---------- ---------- ---------- ---------- 43
NP 001093607 myosin binding protein Hb Danio rerio                      ---------- ---------- ---------- ---------- ---------- 38
Consensus                                                              ---------- ---------- ---GLF-EKP QSG-V-VG-D ITF-AKV-G-
```

Conservation (100% / 0%)

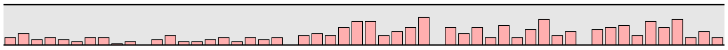

```
                                                                                               220                           240
                                                                                                 |                             |
XP 046765984 myosin-binding protein C slow-type Gallus gallus          DLLRKPNVKW FKGKWMDLAS KAGKHLQLKE SFERHTKIHT FEMHIIQAKE 165
NP 001038124 myosin-binding protein C fast-type Gallus gallus          ALPCAPAVKW FKGKWAELGD KSARC-RLRH SVD-DDKVHT FELTITKVAM 124
NP 001384345 myosin-binding protein C cardiac-type Gallus gallus       SLLKKPSVKW FKGKWMDLAS KVGKHLQLHD NYDRNNKVYT FEMEIIEANM 221
NP 001026199 myosin-binding protein H Gallus gallus                    ---------- ---------- ---------- ---------- ---------- 45
NP 002456 myosin-binding protein C slow-type Homo sapiens              DLLRKPTIKW FKGKWMDLAS KAGKHLQLKE TFERHSRVYT FEMQIIKAKD 159
NP 004524 myosin-binding protein C fast-type Homo sapiens              ELPDKPTIKW FKGKWLELGS KSGARFSFKE SHNSASNVYT VELHIGKVVL 128
NP 000247 myosin-binding protein C cardiac-type Homo sapiens           SLLKPPVVKW FKGKWVDLSS KVGQHLQLHD SYDRASKVYL FELHITDAQP 231
NP 004988 myosin-binding protein H Homo sapiens                        ---------- ---------- ---------- ---------- ---------- 27
NP 001010985 myosin-binding protein H-like Homo sapiens                ---------- ---------- ---------- ---------- ---------- 27
NP 001094228 myosin-binding protein C slow-type Rattus norvegicus      DLLRKPTVKW FKGKWMDLAS KAGKHLQLKE TFERQTRIYT FEMQIIKAKE 160
NP 001381991 myosin-binding protein C fast-type Rattus norvegicus      ELPGKPSIKW FKGKWQELGS KSGARFTFKE SHDSASNVYT VELHIGKVVL 126
NP 001099960 myosin-binding protein C cardiac-type Rattus norvegicus   SLLKPPVVKW FKGKWVDLSS KVGQHLQLHD SYDRASKVYL FELHITDAQA 235
NP 114001 myosin-binding protein H Rattus norvegicus                   ---------- ---------- ---------- ---------- ---------- 27
NP 001014064 myosin-binding protein H-like Rattus norvegicus           ---------- ---------- ---------- ---------- ---------- 28
NP 001007323 myosin-binding protein C slow-type Danio rerio            DLLRKPTVKW FKGKWMDLAS KTGKHLQLKE SFERLSKIHT FEMHIIKAKE 141
NP 001039324 myosin binding protein C2a Danio rerio                     QMLRKPAVKW FKGKWLDLAS KAGKHLQFKE TYDRNTKIYT YEMTIVKVVD 112
NP 001013529 myosin binding protein C2b Danio rerio                     NLTRKPTMKW LKGKWMDLGS KAGKHIQLKE TYDRNTKIYT YEMKLVKVVP 103
NP 001037814 myosin-binding protein C cardiac-type Danio rerio         SLLKKPTVKW FKGKWMDLAS KSGKHLQLKE HYDRNTKVYT FEMHIIAAKA 247
NP 956852 myosin binding protein Ha Danio rerio .                      ---------- ---------- ---------- ---------- ---------- 43
NP 001093607 myosin binding protein Hb Danio rerio                      ---------- ---------- ---------- ---------- ---------- 38
Consensus                                                              -LL-KP-VKW FKGKWMDLAS K-GKHLQLKE S-DR--KVYT FEMHI-KA--
```

Conservation (100% / 0%)

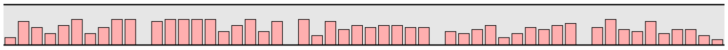

```
                                                                        260                           280                           300
                                                                          |                             |                             |
XP 046765984 myosin-binding protein C slow-type Gallus gallus          NYAGNYRCEV SYKDKFDSCS FDLEVTESSQ AAPSIDIRSA FKRSGDGQ-- 213
NP 001038124 myosin-binding protein C fast-type Gallus gallus          GDRGDYRCEV TAKEQKDSCS FSIDV-EAPR -SSEGNVLQA FKRTGEGKD- 171
NP 001384345 myosin-binding protein C cardiac-type Gallus gallus       TFAGGYRCEV STKDKFDSSN FNLIVNEAPV -SGEMDIRAA FRRTSLAGGG 270
NP 001026199 myosin-binding protein H Gallus gallus                    ---------- ---------- ---------- ---------- ---------- 45
NP 002456 myosin-binding protein C slow-type Homo sapiens              NFAGNYRCEV TYKDKFDSCS FDLEVHESTG TTPNIDIRSA FKRSGEGQ-- 207
NP 004524 myosin-binding protein C fast-type Homo sapiens              GDRGYYRLEV KAKDTCDSCG FNIDV-EAPR QDASGQSLES FKRTSEKKS- 176
NP 000247 myosin-binding protein C cardiac-type Homo sapiens           AFTGSYRCEV STKDKFDCSN FNLTVHEAMG -TGDLDLLSA FRRTSLAGGG 280
NP 004988 myosin-binding protein H Homo sapiens                        ---------- ---------- ---------- ---------- ---------- 27
NP 001010985 myosin-binding protein H-like Homo sapiens                ---------- ---------- ---------- ---------- ---------- 27
NP 001094228 myosin-binding protein C slow-type Rattus norvegicus      NYAGNYRCEV TYKDKFDSCS FDLEVHESTG TTPNIDIRSA FKRSGEGQ-- 208
NP 001381991 myosin-binding protein C fast-type Rattus norvegicus      GDRGDYRLEV KAKDVCDSCP FNVDV-EAPR QDSSGQSLES FKRSGDGKS- 174
NP 001099960 myosin-binding protein C cardiac-type Rattus norvegicus   TSAGGYRCEV STKDKFDSCN FNLTVHEAIG -SGDLDLRSA FRRTSLAGTG 284
NP 114001 myosin-binding protein H Rattus norvegicus                   ---------- ---------- ---------- ---------- ---------- 27
NP 001014064 myosin-binding protein H-like Rattus norvegicus           ---------- ---------- ---------- ---------- ---------- 28
NP 001007323 myosin-binding protein C slow-type Danio rerio            NYAGNYRCEV TYKDKFDSCS FDLEVKEVPE VSQSIDIRSA FKRSSEGQ-- 189
NP 001039324 myosin binding protein C2a Danio rerio                     GDAGGYRCEV TSKDKCDTSA FDVTV-EAAE ETQQTNILEA FKRSGDAG-- 159
NP 001013529 myosin binding protein C2b Danio rerio                     GDAGGYRCEV SAKDKCDSCT FEVTV-EAAE QEQQADILSA FKRADAG--- 149
NP 001037814 myosin-binding protein C cardiac-type Danio rerio         NFAGAYRCEV SSRDKFDSCN FDLIVHEART -TEGFDIRTA FRRTSTDAG- 295
NP 956852 myosin binding protein Ha Danio rerio .                      ---------- ---------- ---------- ---------- ---------- 43
NP 001093607 myosin binding protein Hb Danio rerio                      ---------- ---------- ---------- ---------- ---------- 38
Consensus                                                              --AG-YRCEV --KDKFDSC- F-L-V-EA-- -----DIRSA FKR---G---
```

Conservation (100% / 0%)

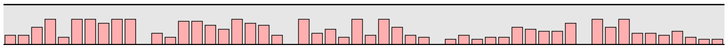

```
                                                                              320                  340
XP 046765984 myosin-binding protein C slow-type Gallus gallus          ---------- ---DDAGELD FSGLLKRR-- --------EV KQQEEEPEVD 240
NP 001038124 myosin-binding protein C fast-type Gallus gallus          ---------- ---DTAGELD FSGLLKKREV QVEEKK--KK KDEDDQFPPE 206
NP 001384345 myosin-binding protein C cardiac-type Gallus gallus       RRMTSAFLST EGLEESGELN FSALLKKRDS FLRTANR-GD GKSDSQPDVD 319
NP 001026199 myosin-binding protein H Gallus gallus                    ---------- ---------- ---------- --------PT PKEEHAPP-- 55
NP 002456 myosin-binding protein C slow-type Homo sapiens              ---------- ---EDAGELD FSGLLKKREV --------KQ --QEEEPQVD 234
NP 004524 myosin-binding protein C fast-type Homo sapiens              ---------- ---DTAGELD FSGLLKKREV VEEEKKKKKK DDDDLGIPPE 213
NP 000247 myosin-binding protein C cardiac-type Homo sapiens           RRISDSH--- ---EDTGILD FSSLLKKRDS FRTP----RD SKLEAPAEED 320
NP 004988 myosin-binding protein H Homo sapiens                        ---------- ---EPPGEVA VS-------- --------ES TREEQVPK-- 46
NP 001010985 myosin-binding protein H-like Homo sapiens                ---------- ---------- ---------- ---------- ---------- 27
NP 001094228 myosin-binding protein C slow-type Rattus norvegicus      ---------- ---EDAGELD FSGLLKRREV --------KQ --QEEEPEID 235
NP 001381991 myosin-binding protein C fast-type Rattus norvegicus      ---------- ---EDAGELD FSGLLKKREV VEEE-KKKKK DDDDLGIPPE 210
NP 001099960 myosin-binding protein C cardiac-type Rattus norvegicus   RRTSDSH--- ---EDAGTLD FSSLLKKSSF R-------RD SKLEAPAEED 321
NP 114001 myosin-binding protein H Rattus norvegicus                   ---------- ---EPSGQVA AS-------- --------ES TGQEQAPE-- 46
NP 001014064 myosin-binding protein H-like Rattus norvegicus           ---------- ---------- ---------- ---------- ---------- 28
NP 001007323 myosin-binding protein C slow-type Danio rerio            ---------- ---DDAGELD FSGLLKHRHG --------RE VKQEETPEVD 218
NP 001039324 myosin binding protein C2a Danio rerio                    ---------- ---EDAGDLD FSALLKKRE- --------RK QKEEPEVDVD 187
NP 001013529 myosin binding protein C2b Danio rerio                    ---------- ---EDEGDLD FSALLKAAK- --------KK KKPEPELEVD 177
NP 001037814 myosin-binding protein C cardiac-type Danio rerio         ---------- ---DDSGELD FSALLKKRDS FVKTGQ--RA VHVSTEPDVD 330
NP 956852 myosin binding protein Ha Danio rerio .                      ---------- ---------- ---------- ---------- ---------- 43
NP 001093607 myosin binding protein Hb Danio rerio                     ---------- ---------- ---------- --------AE PAPEPAPAE- 49
Consensus                                                              ---------- ---EDAGELD FSGLLKKR-- --------KK -KQEEEPEVD
```

Conservation

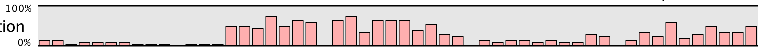

```
                                                                                   360                  380                  400
XP 046765984 myosin-binding protein C slow-type Gallus gallus          VWELLKN-AN PSEYEKIAFQ YGITDLRGML KRLKRMRR-E VKKSAAFAKG 288
NP 001038124 myosin-binding protein C fast-type Gallus gallus          IWELLKGVTK KSEYERIAFQ YGITDLRGML KRLKKVHV-E PKKSEAFIRK 255
NP 001384345 myosin-binding protein C cardiac-type Gallus gallus       VWEILRK-AP PSEYEKIAFQ YGITDLRGML KRLKRIKK-E EKKSTAFLKK 367
NP 001026199 myosin-binding protein H Gallus gallus                    ---------- ---------- ---------- ---------- ---------- 55
NP 002456 myosin-binding protein C slow-type Homo sapiens              VWELLKN-AK PSEYEKIAFQ YGITDLRGML KRLKRMRR-E EKKSAAFAKI 282
NP 004524 myosin-binding protein C fast-type Homo sapiens              IWELLKG-AK KSEYEKIAFQ YGITDLRGML KRLKKAKV-E VKKSAAFTKK 261
NP 000247 myosin-binding protein C cardiac-type Homo sapiens           VWEILRQ-AP PSEYERIAFQ YGVTDLRGML KRLKGMRR-D EKKSTAFQKK 368
NP 004988 myosin-binding protein H Homo sapiens                        ---------- ---------- ---------- ---------- ---------- 46
NP 001010985 myosin-binding protein H-like Homo sapiens                ---------- ---------- ---------- ---------- ---------- 27
NP 001094228 myosin-binding protein C slow-type Rattus norvegicus      VWELLKN-AN PNEYEKIAFQ YGITDLRGML KRLKRMRR-V EKKSAAFAKI 283
NP 001381991 myosin-binding protein C fast-type Rattus norvegicus      IWELLKG-AK KSEYEKIAFQ YGITDLRGML KRLKKAKV-E VKKSAAFTKK 258
NP 001099960 myosin-binding protein C cardiac-type Rattus norvegicus   VWEILRQ-AP PSEYERIAFQ HGVTDLRGML KRLKGMKH-D EKKSTAFQKK 369
NP 114001 myosin-binding protein H Rattus norvegicus                   ---------- ---------- ---------- ---------- ---------- 46
NP 001014064 myosin-binding protein H-like Rattus norvegicus           ---------- ---------- ---------- ---------- ---------- 28
NP 001007323 myosin-binding protein C slow-type Danio rerio            VWEILKN-AR PDEYEKIAFT YGITDLRGLL KRLKKTKK-E EKKSEAFAKR 266
NP 001039324 myosin binding protein C2a Danio rerio                    VWEILKN-AK PCDYEKIAFE YGITDLRGML TRLKKMKKGE PKKSDAFLKK 236
NP 001013529 myosin binding protein C2b Danio rerio                    VWELLKS-AH PSEYEKIAFK YGITDLRGML KRLKKMKVVE PKFSEAFLRR 226
NP 001037814 myosin-binding protein C cardiac-type Danio rerio         VWDILQK-AP PSEYEKIAFQ YGITDLRGML KRLKKMKK-E EKKSAAFLKK 378
NP 956852 myosin binding protein Ha Danio rerio .                      ---------- ---------- ---------- ---------- ---------- 43
NP 001093607 myosin binding protein Hb Danio rerio                     ---------- ---------- ---------- ---------- ---------- 49
Consensus                                                              VWELLK--A- PSEYEKIAFQ YGITDLRGML KRLKKMK--E EKKS-AF-KK
```

Conservation

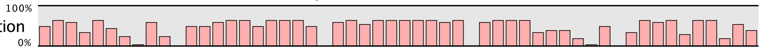

```
                                                                              420                  440
XP 046765984 myosin-binding protein C slow-type Gallus gallus          LDPAYQVDKG GKVRFMVELA DPTVELKWYK NGQEIRPS-- A-KYIFEHKG 335
NP 001038124 myosin-binding protein C fast-type Gallus gallus          LDPAYQVDKG NKIKLVVELS DPDLPLKWYK NGQLLKPS-- T-KYVFENVG 302
NP 001384345 myosin-binding protein C cardiac-type Gallus gallus       LDPAYQVDKG QKIKLMVEVA NPDADVKWLK NGQEIQVS-- GSKYIFEAIG 415
NP 001026199 myosin-binding protein H Gallus gallus                    ---------- ---------- ---------- ---------- ---------- 55
NP 002456 myosin-binding protein C slow-type Homo sapiens              LDPAYQVDKG GRVRFVVELA DPKLEVKWYK NGQEIRPS-- T-KYIFEHKG 329
NP 004524 myosin-binding protein C fast-type Homo sapiens              LDPAYQVDRG NKIKLMVEIS DPDLTLKWFK NGQEIKPS-- S-KYVFENVG 308
NP 000247 myosin-binding protein C cardiac-type Homo sapiens           LEPAYQVSKG HKIRLTVELA DHDAEVKWLK NGQEIQMS-- GSKYIFESIG 416
NP 004988 myosin-binding protein H Homo sapiens                        ---------- ---------- ---------- ---------- ---------- 46
NP 001010985 myosin-binding protein H-like Homo sapiens                ---------- ---------- ---------- ---------- ---------- 27
NP 001094228 myosin-binding protein C slow-type Rattus norvegicus      LDPAYQIDKG GRVRFVVELA DPKLEVKWFK NGQEIRPS-- T-KYIFEHKG 330
NP 001381991 myosin-binding protein C fast-type Rattus norvegicus      LDPAYQVDRG NKIKLVVEIS DPDLPLKWFK NGQEIKPS-- S-KYVFENVG 305
NP 001099960 myosin-binding protein C cardiac-type Rattus norvegicus   LEPAYQVNKG HKIRLTVELA DPDAEVKWLK NGQEIQMS-- G-RYIFESIG 416
NP 114001 myosin-binding protein H Rattus norvegicus                   ---------- ---------- ---------- ---------- ---------- 46
NP 001014064 myosin-binding protein H-like Rattus norvegicus           ---------- ---------- ---------- ---------- ---------- 28
NP 001007323 myosin-binding protein C slow-type Danio rerio            LEPAYQVDKG GKIRLIVDLA DPTVELKWYK NGQEIRPTPN NRKFIFEHKG 316
NP 001039324 myosin binding protein C2a Danio rerio                    LDPAYSVEKG KKIQLSVELT DPNAQVKWLK NGQEIKPS-- A-KYVFESVG 283
NP 001013529 myosin binding protein C2b Danio rerio                    LESAYSVNKG KKIVMSVEVA DPNAEVKWLR NGQEIKPS-- A-KYIMEADG 273
NP 001037814 myosin-binding protein C cardiac-type Danio rerio         LDPAYQVEKG HKIKLEIEVA NPDAEVKWLK NGQEIHPT-- GSKYIFESVG 426
NP 956852 myosin binding protein Ha Danio rerio .                      ---------- ------VEAA AP-------- ---------- ---------- 49
NP 001093607 myosin binding protein Hb Danio rerio                     ---------- ---------- ---------- ---------- ---------- 49
Consensus                                                              LDPAYQVDKG -KI-L-VELA DPD-EVKW-K NGQEI-PS-- --KYIFE--G
```

Conservation

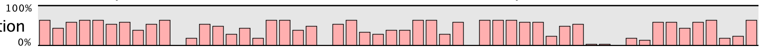

```
                                                                                       460                  480                  500
                                                                                         |                    |                    |
XP 046765984 myosin-binding protein C slow-type Gallus gallus             NQRILFINNC TMTDDARYYV TAGDEKCSTE LFVRDPPILV TKGLEDTSTY 385
NP 001038124 myosin-binding protein C fast-type Gallus gallus             LKRILTIHKC SLADDAAYEC RVNDEKCFTE VFVKEPPVTV VRGLEDQQVV 352
NP 001384345 myosin-binding protein C cardiac-type Gallus gallus          NKRILTINHC SLADDAAYEC VVAEEKSFTE LFVKEPPILI THPLEDQMVM 465
NP 001026199 myosin-binding protein H Gallus gallus                       ---------- ---------- ---------- ---------- ---------- 55
NP 002456 myosin-binding protein C slow-type Homo sapiens                 CQRILFINNC QMTDDSEYYV TAGDEKCSTE LFVREPPIMV TKQLEDTTAY 379
NP 004524 myosin-binding protein C fast-type Homo sapiens                 KKRILTINKC TLADDAAYEV AVKDEKCFTE LFVKEPPVLI VTPLEDQQVF 358
NP 000247 myosin-binding protein C cardiac-type Homo sapiens              AKRTLTISQC SLADDAAYQC VVGGEKCSTE LFVKEPPVLI TRPLEDQLVM 466
NP 004988 myosin-binding protein H Homo sapiens                           ---------- ---------- ---------- ---------- ---------- 46
NP 001010985 myosin-binding protein H-like Homo sapiens                   ---------- ---------- ---------- ---------- ---------- 27
NP 001094228 myosin-binding protein C slow-type Rattus norvegicus         NERIMFINNC ALTDDSEYYV TAGDEKCSTE LFVREPPIMV TKQLEDMNAY 380
NP 001381991 myosin-binding protein C fast-type Rattus norvegicus         KKRILTINKC TLADDAAYEV AVKDEKCFTE LFVKEPPVLI VTPLEDQQVF 355
NP 001099960 myosin-binding protein C cardiac-type Rattus norvegicus      AKRTLTISQC SLADDAAYQC VVGGEKCSTE LFVKEPPVLI TRSLEDQLVM 466
NP 114001 myosin-binding protein H Rattus norvegicus                      ---------- ---------- ---------- ---------- ---------- 46
NP 001014064 myosin-binding protein H-like Rattus norvegicus              ---------- ---------- ---------- ---------- ---------- 28
NP 001007323 myosin-binding protein C slow-type Danio rerio               TQRIMVINNC QMSDDAAYSV TAGEEKCTTE LFVKELPVNI VKELEPVKTT 366
NP 001039324 myosin binding protein C2a Danio rerio                       GIRKLTINKC TLADDAAYEC VVGEEKSFTE VFIQEPPVTI TKMLDDAHVV 333
NP 001013529 myosin binding protein C2b Danio rerio                       NIRTLTINKC SLADDAAYEC VVGTDKCFTE VFVKEPPITI TKLLDDYHVV 323
NP 001037814 myosin-binding protein C cardiac-type Danio rerio            NKRFLTINNC SLSDDAAYMC VVGDEKTVTE LFVKEPPVLI VRNLEDQMAM 476
NP 956852 myosin binding protein Ha Danio rerio .                         ---------- ---------- ---------- ---------- ---------- 49
NP 001093607 myosin binding protein Hb Danio rerio                        ---------- ---------- ---------- ---------- ---------- 49
Consensus                                                                 -KRILTIN-C -LADDAAY-C -VGDEKC-TE LFVKEPPVLI T--LEDQ-V-
```

Conservation

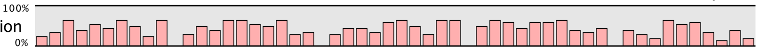


```
                                                                                                  520                  540
                                                                                                    |                    |
XP 046765984 myosin-binding protein C slow-type Gallus gallus             VGERVELSCE VSEDDANVKW FKNGVELTNE PKSR--YRIK VEGKKHTLII 433
NP 001038124 myosin-binding protein C fast-type Gallus gallus             VGDRVVLEAE VSEEGAQVMW LKDGVDVTRD DAFK--YRFK KDGKKHFLII 400
NP 001384345 myosin-binding protein C cardiac-type Gallus gallus          VGERVEFECE VSEEGATVKW EKDGVELTRE ETFK--YRFK KDGKKQYLII 513
NP 001026199 myosin-binding protein H Gallus gallus                       ---------- ---------- ---------- ---------- ---------- 55
NP 002456 myosin-binding protein C slow-type Homo sapiens                 CGERVELECE VSEDDANVKW FKNGEEIIPG PKSR--YRIR VEGKKHILII 427
NP 004524 myosin-binding protein C fast-type Homo sapiens                 VGDRVEMAVE VSEEGAQVMW MKDGVELTRE DSFKARYRFK KDGKRHILIF 408
NP 000247 myosin-binding protein C cardiac-type Homo sapiens              VGQRVEFECE VSEEGAQVKW LKDGVELTRE ETFK--YRFK KDGQRHHLII 514
NP 004988 myosin-binding protein H Homo sapiens                           ---------- ---------- ---------- ---------- ---------- 46
NP 001010985 myosin-binding protein H-like Homo sapiens                   ---------- ---------- ---------- ---------- ---------- 27
NP 001094228 myosin-binding protein C slow-type Rattus norvegicus         CGERVELEVE VSEDDANVKW FKNGEEIVPG PKSR--YRIK VEGKKHTLII 428
NP 001381991 myosin-binding protein C fast-type Rattus norvegicus         VGDRVEMSVE VSEEGAQVMW MKDGVEMTRE DSYKARYRFK KDGKRHILIY 405
NP 001099960 myosin-binding protein C cardiac-type Rattus norvegicus      VGQRVEFECE VSEEGAQVKW LKDGVELTRE ETFK--YRFK KDGRKHHLII 514
NP 114001 myosin-binding protein H Rattus norvegicus                      ---------- ---------- ---------- ---------- ---------- 46
NP 001014064 myosin-binding protein H-like Rattus norvegicus              ---------- ---------- ---------- ---------- ---------- 28
NP 001007323 myosin-binding protein C slow-type Danio rerio               VNERIELECE VSEEGAKVKW MKNGVEVPTG VRSR--YRVK SEGTKHWLII 414
NP 001039324 myosin binding protein C2a Danio rerio                       EGDKVEFEVE VSDEGANVKW MKDGVELSRD GK----YRFK KDGTKHWLII 379
NP 001013529 myosin binding protein C2b Danio rerio                       VGERVEFEIE VSEEGAHVIW LFEDQQLSRE DKDS-KYRFK KDGKRHCLII 372
NP 001037814 myosin-binding protein C cardiac-type Danio rerio            KGDRVEFECE VSEEGAQVKW EKDGVELTRD ESFK--YRFK KDGCKHVLII 524
NP 956852 myosin binding protein Ha Danio rerio .                         ---------- ---------- ---------- ---------- ---------- 49
NP 001093607 myosin binding protein Hb Danio rerio                        ---------- ---------- ---------- ---------- ---------- 49
Consensus                                                                 VG-RVE-ECE VSEEGA-VKW -KDGVELTRE ------YRFK KDGKKH-LII
```

Conservation

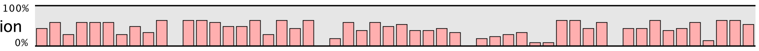


```
                                                                                       560                  580                  600
                                                                                         |                    |                    |
XP 046765984 myosin-binding protein C slow-type Gallus gallus             EEAAKNDNAT YSVMTTGGQS EAKLSVDLRP LKISLALEDQ TVRLGQEIHL 483
NP 001038124 myosin-binding protein C fast-type Gallus gallus             NEAELSDSAH YKIMTNGGES EAELSVEEKQ LEVLQDMADL TVKASEQAVF 450
NP 001384345 myosin-binding protein C cardiac-type Gallus gallus          NESTKEDSGH YTVKTNGGVS VAELIVQEKK LEVYQSIADL TVKARDQAVF 563
NP 001026199 myosin-binding protein H Gallus gallus                       ---------- ---------- ---------- ---------- ---------- 55
NP 002456 myosin-binding protein C slow-type Homo sapiens                 EGATKADAAE YSVMTTGGQS SAKLSVDLKP LKILTPLTDQ TVNLGKEICL 477
NP 004524 myosin-binding protein C fast-type Homo sapiens                 SDVVQEDRGR YQVITNGGQC EAELIVEEKQ LEVLQDIADL TVKASEQAVF 458
NP 000247 myosin-binding protein C cardiac-type Homo sapiens              NEAMLEDAGH YALCTSGGQA LAELIVQEKK LEVYQSIADL MVGAKDQAVF 564
NP 004988 myosin-binding protein H Homo sapiens                           ---------- ---------- ---------- ---------- ---------- 46
NP 001010985 myosin-binding protein H-like Homo sapiens                   ---------- ---------- ---------- ---------- ---------- 27
NP 001094228 myosin-binding protein C slow-type Rattus norvegicus         EGATKADSAE YSVMTTGGQS SAKLSVDLRP LKITTPLTDQ TVKLGKEVCL 478
NP 001381991 myosin-binding protein C fast-type Rattus norvegicus         SDVAQEDGGR YQVITNGGQC EAELIVEEKQ LEVLQDIADL TVKASEQAVF 455
NP 001099960 myosin-binding protein C cardiac-type Rattus norvegicus      NEATLEDAGH YAVRTSGGQA LAELIVQEKK LEVYQSIADL AVGAKDQAVF 564
NP 114001 myosin-binding protein H Rattus norvegicus                      ---------- ---------- ---------- ---------- ---------- 46
NP 001014064 myosin-binding protein H-like Rattus norvegicus              ---------- ---------- ---------- ---------- ---------- 28
NP 001007323 myosin-binding protein C slow-type Danio rerio               DDASKDDTGT YSLMATGGTS QAHVQVDLKP LKVLMDLQSV TVLLGQPLKL 464
NP 001039324 myosin binding protein C2a Danio rerio                       NEATTEDIGT YYVYTSGGES KAELEVEERE LEVLQSIADL TVKAAEQAVF 429
NP 001013529 myosin binding protein C2b Danio rerio                       QEATLEDNGM FYVYTNGGQS KGELIVEEKE LEVLQSIADL TVKSAEQAMF 422
NP 001037814 myosin-binding protein C cardiac-type Danio rerio            NDVTKEDCGH YRVKTNGGQS LAELMVQEKQ LEVYQSIADL TVKAKDQAVF 574
NP 956852 myosin binding protein Ha Danio rerio .                         ---------- ---------- ---------- ---------- ---------- 49
NP 001093607 myosin binding protein Hb Danio rerio                        ---------- ---------- ---------- ---------- ---------- 49
Consensus                                                                 -EAT-ED-G- Y-V-T-GGQS -AEL-V-EK- LEVLQ-IADL TVKA--QAVF
```

Conservation

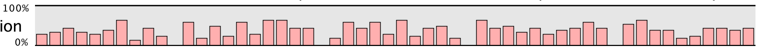

| | | 620 | | 640 | | |
|---|---|---|---|---|---|---|
| XP 046765984 myosin-binding protein C slow-type Gallus gallus | KCEIS-ENVE | GKWYKNGQLV | EASDRVKLYH | KGRIHRFIIA | SAAVDDEGEY | 532 |
| NP 001038124 myosin-binding protein C fast-type Gallus gallus | KCEVSDEKVT | GRWFRNGVEV | KPSKRIHISH | NGRFHKLVID | DVRPEDEGDY | 500 |
| NP 001384345 myosin-binding protein C cardiac-type Gallus gallus | KCEVSDENVK | GIWLKNGKEV | VPDERIKISH | IGRIHKLTIE | DVTPGDEADY | 613 |
| NP 001026199 myosin-binding protein H Gallus gallus | ---------- | ---------- | ---------- | ---------- | ---------- | 55 |
| NP 002456 myosin-binding protein C slow-type Homo sapiens | KCEI-SENIP | GKWTKNGLPV | QESDRLKVVH | KGRIHKLVIA | NALTEDEGDY | 526 |
| NP 004524 myosin-binding protein C fast-type Homo sapiens | KCEVSDEKVT | GKWYKNGVEV | RPSKRITISH | VGRFHKLVID | DVRPEDEGDY | 508 |
| NP 000247 myosin-binding protein C cardiac-type Homo sapiens | KCEVSDENVR | GVWLKNGKEL | VPDSRIKVSH | IGRVHKLTID | DVTPADEADY | 614 |
| NP 004988 myosin-binding protein H Homo sapiens | ---------- | ---------- | ---------- | ---------- | ---------- | 46 |
| NP 001010985 myosin-binding protein H-like Homo sapiens | ---------- | ---------- | ---------- | ---------- | ---------- | 27 |
| NP 001094228 myosin-binding protein C slow-type Rattus norvegicus | KCEI-SENVP | GKWTKNGLPV | QEGERLKVVH | KGRIHKLVIA | NALIEDEGEY | 527 |
| NP 001381991 myosin-binding protein C fast-type Rattus norvegicus | KCEVSDEKVT | GKWYKNGVEV | RPSKRITISH | VGRFHKLVID | DVRPEDEGDY | 505 |
| NP 001099960 myosin-binding protein C cardiac-type Rattus norvegicus | KCEVSDENVR | GVWLKNGKEL | VPDNRIKVSH | IGRVHKLTID | DVTPADEADY | 614 |
| NP 114001 myosin-binding protein H Rattus norvegicus | ---------- | ---------- | ---------- | ---------- | ---------- | 46 |
| NP 001014064 myosin-binding protein H-like Rattus norvegicus | ---------- | ---------- | ---------- | ---------- | ---------- | 28 |
| NP 001007323 myosin-binding protein C slow-type Danio rerio | NCEIYPGNVP | GRWYKNGQLV | QPNDRISITH | KTKNHSLDVE | SSTIHDAGDY | 514 |
| NP 001039324 myosin binding protein C2a Danio rerio | KCEVSDEKVT | GKWFKDGVEV | VASDRIKMSH | IGRTHKLVIS | DVKPEDAGDY | 479 |
| NP 001013529 myosin binding protein C2b Danio rerio | KCEVSDEKVT | GKWFKDGVEV | LPSDRIKISH | IGRIHKLTID | DVKPGDAGDY | 472 |
| NP 001037814 myosin-binding protein C cardiac-type Danio rerio | KCEVSDENVK | GIWYKNGVEV | KPDARTLITH | IGRIHKLTID | DVKPEDEGDY | 624 |
| NP 956852 myosin binding protein Ha Danio rerio . | ---------- | ---------- | ---------- | ---------- | ---------- | 49 |
| NP 001093607 myosin binding protein Hb Danio rerio | ---------- | ---------- | ---------- | ---------- | ---------- | 49 |
| Consensus | KCEVSDENV- | GKW-KNG-EV | -PS-RIKISH | -GR-HKL-ID | DV-PEDEGDY | |

Conservation 100% 0%

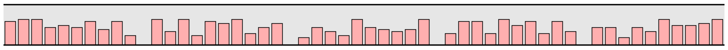

| | 660 | | 680 | | 700 | |
|---|---|---|---|---|---|---|
| XP 046765984 myosin-binding protein C slow-type Gallus gallus | MFVPDAYNIN | IPCKVHVV-- | --------DP | PKLHLDGLGE | ---NNTVTVV | 569 |
| NP 001038124 myosin-binding protein C fast-type Gallus gallus | TFIPDGYALS | LSAKLNFLEI | KVEYVPKQEP | PKIHLDCSGK | -AAENTIVVV | 549 |
| NP 001384345 myosin-binding protein C cardiac-type Gallus gallus | SFIPQGFAYN | LSAKLQFLEV | KIDFVPREEP | PKIHLDCLGQ | -SPD-TIVVV | 661 |
| NP 001026199 myosin-binding protein H Gallus gallus | ---------- | ---------- | ---------- | PKEEHAPPPK | ---------- | 65 |
| NP 002456 myosin-binding protein C slow-type Homo sapiens | VFAPDAYNVT | LPAKVHVI-- | --------DP | PKIILDGLDA | ---DNTVTVI | 563 |
| NP 004524 myosin-binding protein C fast-type Homo sapiens | TFVPDGYALS | LSAKLNFLEI | KVEYVPKQEP | PKIHLDCSGK | -TSENAIVVV | 557 |
| NP 000247 myosin-binding protein C cardiac-type Homo sapiens | SFVPEGFACN | LSAKLHFMEV | KIDFVPRQEP | PKIHLDCPGR | -IPD-TIVVV | 662 |
| NP 004988 myosin-binding protein H Homo sapiens | ---------- | ---------- | ---------- | PQ-------- | ---------- | 48 |
| NP 001010985 myosin-binding protein H-like Homo sapiens | ---------- | ---------- | ---------- | PQ---ASPGQ | ---------- | 34 |
| NP 001094228 myosin-binding protein C slow-type Rattus norvegicus | VFTPDAYNVP | LSAKVHVI-- | --------DP | PKIILDGLDA | ---DNTVTVI | 564 |
| NP 001381991 myosin-binding protein C fast-type Rattus norvegicus | TFVPDGYALS | LSAKLNFLEI | KVEYVPKQEP | PKIHLDCSGK | -TSDNSIVVV | 554 |
| NP 001099960 myosin-binding protein C cardiac-type Rattus norvegicus | SFVPEGFACN | LSAKLHFMEV | KIDFVPRQEP | PKIHLDCPGS | -TPD-TIVVV | 662 |
| NP 114001 myosin-binding protein H Rattus norvegicus | ---------- | ---------- | ---------- | PQ-------- | ---------- | 48 |
| NP 001014064 myosin-binding protein H-like Rattus norvegicus | ---------- | ---------- | ---------- | PQ-------- | ---------- | 30 |
| NP 001007323 myosin-binding protein C slow-type Danio rerio | TFVPEGYTQT | LSAKVHII-- | --------DP | PRVHLEALN- | -VQDNTVTIV | 552 |
| NP 001039324 myosin binding protein C2a Danio rerio | TFVPDGYALS | LSAKLNFLEI | KIDYVPRQEP | PKIHLDATGS | MVNKNTIIVV | 529 |
| NP 001013529 myosin binding protein C2b Danio rerio | TFVPDGYALS | ISAKLNFLEI | KIDYVPRQDP | PKIHLDVSGN | MVSQNTIIVV | 522 |
| NP 001037814 myosin-binding protein C cardiac-type Danio rerio | TFVPEGFAFN | LSAKLNFLEV | KIDFVPRQDP | PKIHLDCMGR | -TAESTILVV | 673 |
| NP 956852 myosin binding protein Ha Danio rerio . | ---------- | ---------- | ---------- | AAVEETTPAP | ---------- | 59 |
| NP 001093607 myosin binding protein Hb Danio rerio | ---------- | ---------- | ---------- | PASAEAAPAE | GAPA------ | 63 |
| Consensus | TFVPDGYA-- | LSAKL-FL-- | --------EP | PKIHLDCPGK | ---D-TI-VV |  |

Conservation 100% 0%

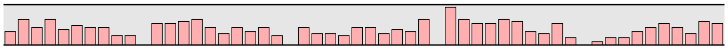

| | | 720 | | 740 | | |
|---|---|---|---|---|---|---|
| XP 046765984 myosin-binding protein C slow-type Gallus gallus | AGTKLRLEIP | ITGEPTPKVM | WS----KGD- | ---------- | ---------- | 594 |
| NP 001038124 myosin-binding protein C fast-type Gallus gallus | AGNKVRLDVP | ISGEPAPTVT | WK----RGD- | ---------- | ---------- | 574 |
| NP 001384345 myosin-binding protein C cardiac-type Gallus gallus | AGNKLRLDVP | ISGDPTPTVI | WQKVNKKGEL | VH-QSNEDSL | TPSENSSDLS | 710 |
| NP 001026199 myosin-binding protein H Gallus gallus | ---EEHAPAP | AAETPPAP-- | ---------- | ---------- | ---------- | 80 |
| NP 002456 myosin-binding protein C slow-type Homo sapiens | AGNKLRLEIP | ISGEPPPKAM | WS----RGD- | ---------- | ---------- | 588 |
| NP 004524 myosin-binding protein C fast-type Homo sapiens | AGNKLRLDVS | ITGEPPPVAT | WL----KGD- | ---------- | ---------- | 582 |
| NP 000247 myosin-binding protein C cardiac-type Homo sapiens | AGNKLRLDVP | ISGDPAPTVI | WQKAITQGNK | APARPAPDAP | EDTGDSDEWV | 712 |
| NP 004988 myosin-binding protein H Homo sapiens | ---------- | ---APAPQ-- | ---------- | ---------- | ---------- | 53 |
| NP 001010985 myosin-binding protein H-like Homo sapiens | ---------- | ---------- | ---------- | ---------- | ---------- | 34 |
| NP 001094228 myosin-binding protein C slow-type Rattus norvegicus | AGSKLRLEIP | VTGEPPPKAI | WS----RAD- | ---------- | ---------- | 589 |
| NP 001381991 myosin-binding protein C fast-type Rattus norvegicus | AGNKLRLDVA | ITGEPPPTAT | WL----KGD- | ---------- | ---------- | 579 |
| NP 001099960 myosin-binding protein C cardiac-type Rattus norvegicus | AGNKLRLDVP | ISGDPAPTVI | WQKTITQGKK | ASAGPPPGAP | EDAGADEEWV | 712 |
| NP 114001 myosin-binding protein H Rattus norvegicus | ---------- | ---------- | ---------- | ---------- | ---------- | 48 |
| NP 001014064 myosin-binding protein H-like Rattus norvegicus | ---------- | ---------- | ---------- | ---------- | ---------- | 30 |
| NP 001007323 myosin-binding protein C slow-type Danio rerio | AGNKLRLEIP | ISGEPAPRVV | WM----KGE- | ---------- | ---------- | 577 |
| NP 001039324 myosin binding protein C2a Danio rerio | AGNKLRFDVD | ITGEPPPTVA | WK----KGD- | ---------- | ---------- | 554 |
| NP 001013529 myosin binding protein C2b Danio rerio | AGNKLRLDVE | ITGEPAPTVC | WM----RDD- | ---------- | ---------- | 547 |
| NP 001037814 myosin-binding protein C cardiac-type Danio rerio | AGNKLRLDVP | ITGDPAPTVI | WT----KGE- | ---------- | ---------- | 698 |
| NP 956852 myosin binding protein Ha Danio rerio . | -------EAP | VAEGPAEE-- | ---------- | ---------- | ---------- | 70 |
| NP 001093607 myosin binding protein Hb Danio rerio | -------EAP | AEGPA----- | ---------- | ---------- | ---------- | 72 |
| Consensus | AGNKLRLDVP | ITGEPAPTV- | W-----KGD- | ---------- | ---------- |  |

Conservation 100% 0%

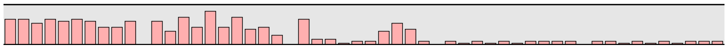

```
                                                                                       760        780        800
XP 046765984 myosin-binding protein C slow-type Gallus gallus            ---KWITD-S GRIRAETYSD SSCLVIDTAE REDSGPFRIT LKNEAGEDSA 640
NP 001038124 myosin-binding protein C fast-type Gallus gallus            ---QLFTATE GRVHIDSQAD LSSFVIESAE RSDEGRYCIT VTNPVGEDSA 621
NP 001384345 myosin-binding protein C cardiac-type Gallus gallus         TDSKLLFESE GRVRVEKHED HCVFIIEGAE KEDEGVYRVI VKNPVGEDKA 760
NP 001026199 myosin-binding protein H Gallus gallus                      ---------- -----EHPPD AEQPAAPAAE HAPTPTHEAA PAHEEGPPPA 115
NP 002456 myosin-binding protein C slow-type Homo sapiens                ---KAIMEGS GRIRTESYPD SSTLVIDIAE RDDSGVYHIN LKNEAGEAHA 635
NP 004524 myosin-binding protein C fast-type Homo sapiens                ---EVFTTTE GRTRIEKRVD CSSFVIESAQ REDEGRYTIK VTNPVGEDVA 629
NP 000247 myosin-binding protein C cardiac-type Homo sapiens             FDKKLLCETE GRVRVETTKD RSIFTVEGAE KEDEGVYTVT VKNPVGEDQV 762
NP 004988 myosin-binding protein H Homo sapiens                          ---------- ---------- ---------- ---------- ---------- 53
NP 001010985 myosin-binding protein H-like Homo sapiens                  ---------- ---------- ---------- ---------- ---------- 34
NP 001094228 myosin-binding protein C slow-type Rattus norvegicus        ---KAIMEGS GRIRAESYPD SSTLVIDVAE RDDSGVYNIN LKNEAGEAHA 636
NP 001381991 myosin-binding protein C fast-type Rattus norvegicus        ---EVFTVSE GRTRIEQRPD CSSFVIESAE RSDEGRYTIK VTNPAGEDVA 626
NP 001099960 myosin-binding protein C cardiac-type Rattus norvegicus     FDKKLLCETE GRVRVETTKD RSVFTVEGAE KEDEGVYTVT VKNPVGEDQV 762
NP 114001 myosin-binding protein H Rattus norvegicus                     ---------- ---------- ---------- ---------- -KQPQAQDPA 57
NP 001014064 myosin-binding protein H-like Rattus norvegicus             ---------- ---------- ---------- ---------- ---------- 30
NP 001007323 myosin-binding protein C slow-type Danio rerio              ---RVILDTG SRVHAETFAD HTCLTIDITE REDTGNYKIV LQNEAGEDTA 624
NP 001039324 myosin binding protein C2a Danio rerio                      ---MEISQAE GRVRVETRKA LSCFVIEGAE RSDEGLYHIT VTNPAGEDKA 601
NP 001013529 myosin binding protein C2b Danio rerio                      ---KPVTDSE GRVRVENKKD LSCFIIEGAE REDEGNYTIT VTNPAGEDKA 594
NP 001037814 myosin-binding protein C cardiac-type Danio rerio           ---KVLTDSD GRVHVESTKG HCIFTIEGAE RQDEGVYSVI VRNPAGEDTA 745
NP 956852 myosin binding protein Ha Danio rerio .                        ---------- ---------- ---------- ---------- -TAPAADAPA 79
NP 001093607 myosin binding protein Hb Danio rerio                       ---------E AAAPAEAPPA EAPPAEPVA- ---------- ---PPAADAE 99
Consensus                                                                ---K-----E GRVRVET-XD -S-FVIEGAE REDEGVY-IT VKNPAGEDXA
```

100%
Conservation
0%

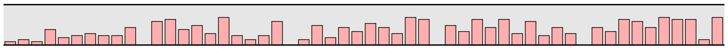

```
                                                                                             820        840
XP 046765984 myosin-binding protein C slow-type Gallus gallus            -LINIKVVDV PDPPQAPNVT EVGEDWCVMT WDPPANDGGS PILGYFIERK 689
NP 001038124 myosin-binding protein C fast-type Gallus gallus            -TLHVRVVDV PDPPQSVRVT SVGEDWAVLS WEAPPFDGGM PITGYLMERK 670
NP 001384345 myosin-binding protein C cardiac-type Gallus gallus         -DITVKVIDV PDPPEAPKIS NIGEDYCTVQ WQPPTYDGGQ PVLGYILERK 809
NP 001026199 myosin-binding protein H Gallus gallus                      -APAEAPAPE PEPEKPKE-- ---------- ---------- ---------- 132
NP 002456 myosin-binding protein C slow-type Homo sapiens                -SIKVKVVDF PDPPVAPTVT EVGDDWCIMN WEPPAYDGGS PILGYFIERK 684
NP 004524 myosin-binding protein C fast-type Homo sapiens                -SIFLQVVDV PDPPEAVRIT SVGEDWAILV WEPPMYDGGK PVTGYLVERK 678
NP 000247 myosin-binding protein C cardiac-type Homo sapiens             -NLTVKVIDV PDAPAAPKIS NVGEDSCTVQ WEPPAYDGGQ PILGYILERK 811
NP 004988 myosin-binding protein H Homo sapiens                          -APTASTATK PAPPSE---- ---------- ---------- ---------- 68
NP 001010985 myosin-binding protein H-like Homo sapiens                  ---------- ---------- ---------- ---------- ---------- 34
NP 001094228 myosin-binding protein C slow-type Rattus norvegicus        -SIKIKVVDI PDPPVAPNVT EVGDDWCIMN WEPPVYDGGS PILGYFIERK 685
NP 001381991 myosin-binding protein C fast-type Rattus norvegicus        -SIFLRVVDV PDPPEAVRVT SVGEDWAILV WEPPKYDGGQ PVTGYLMERK 675
NP 001099960 myosin-binding protein C cardiac-type Rattus norvegicus     -NLTVKVIDV PDAPAAPKIS NVGEDSCIVQ WEPPAYDGGQ PVLGYILERK 811
NP 114001 myosin-binding protein H Rattus norvegicus                     -AHEAPATPA TIKPEAPSE- ---------- ---------- ---------- 75
NP 001014064 myosin-binding protein H-like Rattus norvegicus             ---------- ---------- ---------- ---------- ---------- 30
NP 001007323 myosin-binding protein C slow-type Danio rerio              -SVKVKVVDI PDPPEAPLVT EVGGDWCTMT WEPPRYDGGS PILGYFIERK 673
NP 001039324 myosin binding protein C2a Danio rerio                      -DVFVKIVDV PDPPENVKCL GVGEDSASIE WEPPKFDGGV PVKGYLMERK 650
NP 001013529 myosin binding protein C2b Danio rerio                      -NLFIKIVDV PDPPENVKCT GVGEDTANIV WDPPKFDGGA PLKGYLMERK 643
NP 001037814 myosin-binding protein C cardiac-type Danio rerio           -DINVKVVDV PDPPEAPRIL SVGEDSCVVQ WDAPRFDGGQ PVIGYVLERK 794
NP 956852 myosin binding protein Ha Danio rerio .                        EAAPAEPAPV EEPPKAPT-- ---------- ---------- ---------- 97
NP 001093607 myosin binding protein Hb Danio rerio                       -APPAEEIKA PTPPPPP--- ---------- ---------- ---------- 115
Consensus                                                                -XIXVKVVDV PDPPEAPKVT -VGEDWC--- WEPP-YDGG- P-LGY--ERK
```

100%
Conservation
0%

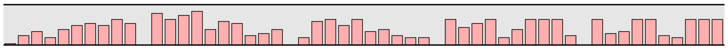

```
                                                                                       860        880        900
XP 046765984 myosin-binding protein C slow-type Gallus gallus            KKQSSRWMRL NFELCKETTF EPKKMIEGVA YEVRVFAVNA IGTSKPSMPS 739
NP 001038124 myosin-binding protein C fast-type Gallus gallus            KKGSMRWMKL NFEVFPDTTY ESTKMIEGVF YEMRVFAVNA IGVSQPSLNT 720
NP 001384345 myosin-binding protein C cardiac-type Gallus gallus         KKKSYRWMRL NFDLLKELTY EAKRMIEGVV YEMRIYAVNS IGMSRPSPAS 859
NP 001026199 myosin-binding protein H Gallus gallus                      ---------- ---------- ---------- ---------- ---------- 132
NP 002456 myosin-binding protein C slow-type Homo sapiens                KKQSSRWMRL NFDLCKETTF EPKKMIEGVA YEVRIFAVNA IGISKPSMPS 734
NP 004524 myosin-binding protein C fast-type Homo sapiens                KKGSQRWMKL NFEVFTETTY ESTKMIEGIL YEMRVFAVNA IGVSQPSMNT 728
NP 000247 myosin-binding protein C cardiac-type Homo sapiens             KKKSYRWMRL NFDLIQELSH EARRMIEGVV YEMRVYAVNA IGMSRPSPAS 861
NP 004988 myosin-binding protein H Homo sapiens                          ---------- ---------- ---------- ---------- ---------- 68
NP 001010985 myosin-binding protein H-like Homo sapiens                  ---------- ---------- ---------- ---------- ---------- 34
NP 001094228 myosin-binding protein C slow-type Rattus norvegicus        KKQSSRWMRL NFDLCKETTF EPKKMIEGVA YEVRIFAVNA IGISKPSMPS 735
NP 001381991 myosin-binding protein C fast-type Rattus norvegicus        KKGSHRWMKL NFEVFTDTTY ESTKMIEGVL YEMRVFAVNA IGVSQPSMNT 725
NP 001099960 myosin-binding protein C cardiac-type Rattus norvegicus     KKKSYRWMRL NFDLLRELSH EARRMIEGVA YEMRVYAVNA VGMSRPSPAS 861
NP 114001 myosin-binding protein H Rattus norvegicus                     ---------- ---------- ---------- ---------- ---------- 75
NP 001014064 myosin-binding protein H-like Rattus norvegicus             ---------- ---------- ---------- ---------- ---------- 30
NP 001007323 myosin-binding protein C slow-type Danio rerio              KKQSSRWMRL NFDLCKETTF EPKKMIEGVP YEVRIFAVNA IGVSKPSEPS 723
NP 001039324 myosin binding protein C2a Danio rerio                      KKGSSRWTKL NFDIYESTTY EAKKMIEGVF YEMRVFAVNG IGISQPSANS 700
NP 001013529 myosin binding protein C2b Danio rerio                      KKGSSRWTKL NFDVYESTTY EAKRMIEGIL YEMRVFAVNG IGISAPSLNS 693
NP 001037814 myosin-binding protein C cardiac-type Danio rerio           KKKSYRWMRL NFDPYKETTF EAKKMIEGVP YEMRVYAVNA IGMSRPSAAS 844
NP 956852 myosin binding protein Ha Danio rerio .                        ---------- ---------- ---------- ---------- ---------- 97
NP 001093607 myosin binding protein Hb Danio rerio                       ---------- ---------- ---------- ---------- ---------- 115
Consensus                                                                KK-S-RWMRL NFDL--ETT- E-KKMIEGV- YEMRVFAVNA IG-S-PS--S
```

100%
Conservation
0%

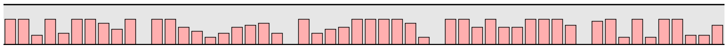

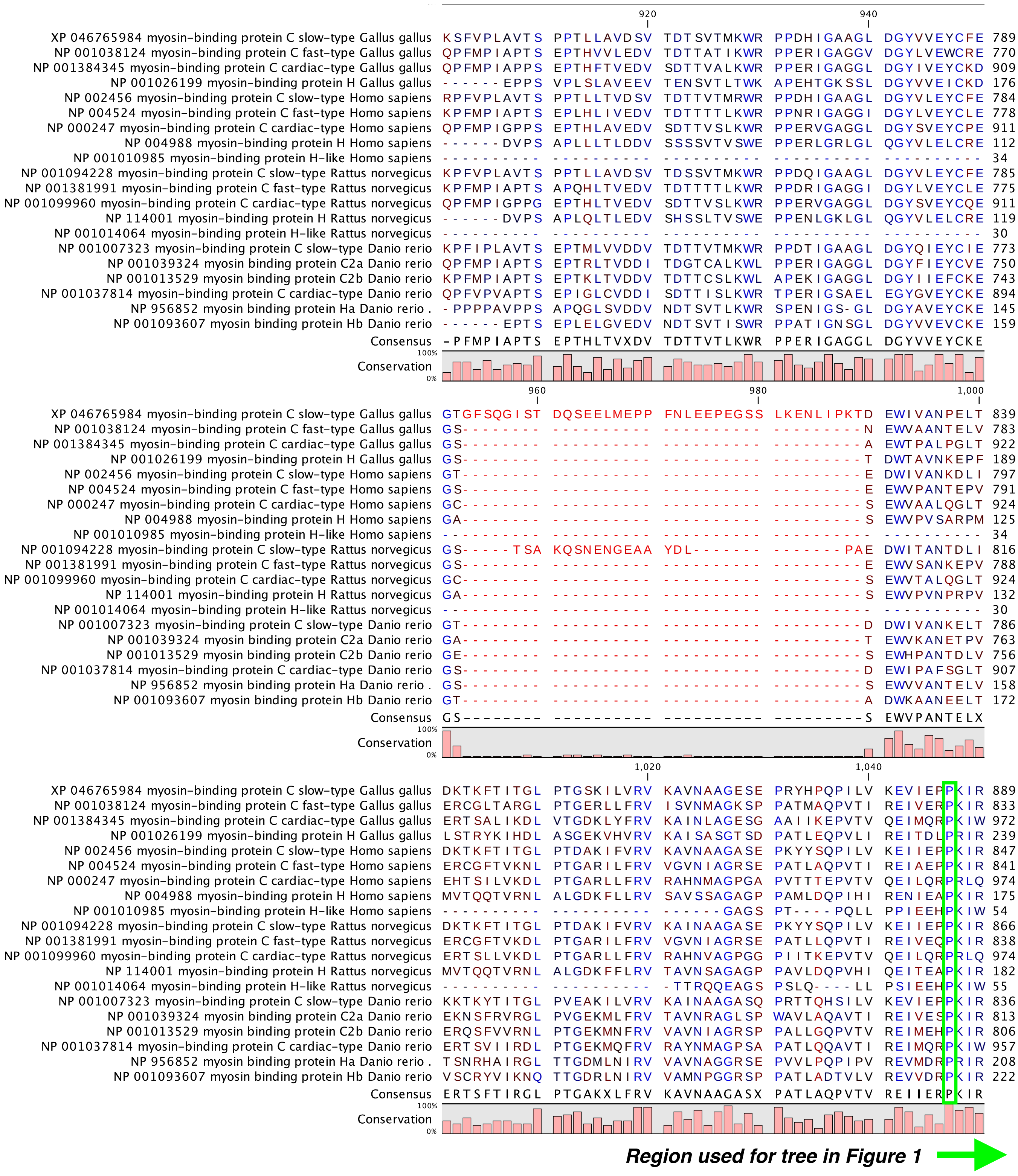
920 940
XP 046765984 myosin-binding protein C slow-type Gallus gallus KSFVPLAVTS PPTLLAVDSV TDTSVTMKWR PPDHIGAAGL DGYVVEYCFE 789
NP 001038124 myosin-binding protein C fast-type Gallus gallus QPFMPIAPTS EPTHVVLEDV TDTTATIKWR PPERIGAGGV DGYLVEWCRE 770
NP 001384345 myosin-binding protein C cardiac-type Gallus gallus QPFMPIAPPS EPTHFTVEDV SDTTVALKWR PPERIGAGGL DGYIVEYCKD 909
NP 001026199 myosin-binding protein H Gallus gallus ------EPPS VPLSLAVEEV TENSVTLTWK APEHTGKSSL DGYVVEICKD 176
NP 002456 myosin-binding protein C slow-type Homo sapiens RPFVPLAVTS PPTLLTVDSV TDTTVTMRWR PPDHIGAAGL DGYVLEYCFE 784
NP 004524 myosin-binding protein C fast-type Homo sapiens KPFMPIAPTS EPLHLIVEDV TDTTTTLKWR PPNRIGAGGI DGYLVEYCLE 778
NP 000247 myosin-binding protein C cardiac-type Homo sapiens QPFMPIGPPS EPTHLAVEDV SDTTVSLKWR PPERVGAGGL DGYSVEYCPE 911
NP 004988 myosin-binding protein H Homo sapiens ------DVPS APLLLTLDDV SSSSVTVSWE PPERLGRLGL QGYVLELCRE 112
NP 001010985 myosin-binding protein H-like Homo sapiens ---------- ---------- ---------- ---------- ---------- 34
NP 001094228 myosin-binding protein C slow-type Rattus norvegicus KPFVPLAVTS PPTLLAVDSV TDSSVTMKWR PPDQIGAAGL DGYVLEYCFE 785
NP 001381991 myosin-binding protein C fast-type Rattus norvegicus KPFMPIAPTS APQHLTVEDV TDTTTTLKWR PPDRIGAGGI DGYLVEYCLE 775
NP 001099960 myosin-binding protein C cardiac-type Rattus norvegicus QPFMPIGPPG EPTHLTVEDV SDTTVSLKWR PPERVGAGGL DGYSVEYCQE 911
NP 114001 myosin-binding protein H Rattus norvegicus ------DVPS APLQLTLEDV SHSSLTVSWE PPENLGKLGL QGYVLELCRE 119
NP 001014064 myosin-binding protein H-like Rattus norvegicus ---------- ---------- ---------- ---------- ---------- 30
NP 001007323 myosin-binding protein C slow-type Danio rerio KPFIPLAVTS EPTMLVVDDV TDTTVTMKWR PPDTIGAAGL DGYQIEYCIE 773
NP 001039324 myosin binding protein C2a Danio rerio QPFMPIAPTS EPTRLTVDDI TDGTCALKWL PPERIGAGGL DGYFIEYCVE 750
NP 001013529 myosin binding protein C2b Danio rerio KPFMPIAPTS EPTKLTVDDV TDTTCSLKWL APEKIGAGGL DGYIIEFCKE 743
NP 001037814 myosin-binding protein C cardiac-type Danio rerio QPFVPVAPTS EPIGLCVDDI SDTTISLKWR TPERIGSAEL EGYGVEYCKE 894
NP 956852 myosin binding protein Ha Danio rerio . -PPPPAVPPS APQGLSVDDV NDTSVTLKWR SPENIGS-GL DGYAVEYCKE 145
NP 001093607 myosin binding protein Hb Danio rerio ------EPTS EPLELGVEDV NDTSVTISWR PPATIGNSGL DGYVVEVCKE 159
Consensus -PFMPIAPTS EPTHLTVXDV TDTTVTLKWR PPERIGAGGL DGYVVEYCKE
100%
Conservation
0%
960 980 1,000
XP 046765984 myosin-binding protein C slow-type Gallus gallus GTGFSQGIST DQSEELMEPP FNLEEPEGSS LKENLIPKTD EWIVANPELT 839
NP 001038124 myosin-binding protein C fast-type Gallus gallus GS-------- ---------- ---------- ---------N EWVAANTELV 783
NP 001384345 myosin-binding protein C cardiac-type Gallus gallus GS-------- ---------- ---------- ---------A EWTPALPGLT 922
NP 001026199 myosin-binding protein H Gallus gallus GS-------- ---------- ---------- ---------T DWTAVNKEPF 189
NP 002456 myosin-binding protein C slow-type Homo sapiens GT-------- ---------- ---------- ---------E DWIVANKDLI 797
NP 004524 myosin-binding protein C fast-type Homo sapiens GS-------- ---------- ---------- ---------E EWVPANTEPV 791
NP 000247 myosin-binding protein C cardiac-type Homo sapiens GC-------- ---------- ---------- ---------S EWVAALQGLT 924
NP 004988 myosin-binding protein H Homo sapiens GA-------- ---------- ---------- ---------S EWVPVSARPM 125
NP 001010985 myosin-binding protein H-like Homo sapiens ---------- ---------- ---------- ---------- ---------- 34
NP 001094228 myosin-binding protein C slow-type Rattus norvegicus GS-----TSA KQSNENGEAA YDL------- -------PAE DWITANTDLI 816
NP 001381991 myosin-binding protein C fast-type Rattus norvegicus GS-------- ---------- ---------- ---------E EWVSANKEPV 788
NP 001099960 myosin-binding protein C cardiac-type Rattus norvegicus GC-------- ---------- ---------- ---------S EWVTALQGLT 924
NP 114001 myosin-binding protein H Rattus norvegicus GA-------- ---------- ---------- ---------S EWVPVNPRPV 132
NP 001014064 myosin-binding protein H-like Rattus norvegicus ---------- ---------- ---------- ---------- ---------- 30
NP 001007323 myosin-binding protein C slow-type Danio rerio GT-------- ---------- ---------- ---------D DWIVANKELT 786
NP 001039324 myosin binding protein C2a Danio rerio GA-------- ---------- ---------- ---------T EWVKANETPV 763
NP 001013529 myosin binding protein C2b Danio rerio GE-------- ---------- ---------- ---------S EWHPANTDLV 756
NP 001037814 myosin-binding protein C cardiac-type Danio rerio GS-------- ---------- ---------- ---------D EWIPAFSGLT 907
NP 956852 myosin binding protein Ha Danio rerio . GS-------- ---------- ---------- ---------S EWVVANTELV 158
NP 001093607 myosin binding protein Hb Danio rerio GT-------- ---------- ---------- ---------A DWKAANEELT 172
Consensus GS-------- ---------- ---------- ---------S EWVPANTELX
100%
Conservation
0%
1,020 1,040
XP 046765984 myosin-binding protein C slow-type Gallus gallus DKTKFTITGL PTGSKILVRV KAVNAAGESE PRYHPQPILV KEVIEPPKIR 889
NP 001038124 myosin-binding protein C fast-type Gallus gallus ERCGLTARGL PTGERLLFRV ISVNMAGKSP PATMAQPVTI REIVERPKIR 833
NP 001384345 myosin-binding protein C cardiac-type Gallus gallus ERTSALIKDL VTGDKLYFRV KAINLAGESG AAIIKEPVTV QEIMQRPKIW 972
NP 001026199 myosin-binding protein H Gallus gallus LSTRYKIHDL ASGEKVHVRV KAISASGTSD PATLEQPVLI REITDLPRIR 239
NP 002456 myosin-binding protein C slow-type Homo sapiens DKTKFTITGL PTDAKIFVRV KAVNAAGASE PKYYSQPILV KEIIEPPKIR 847
NP 004524 myosin-binding protein C fast-type Homo sapiens ERCGFTVKNL PTGARILFRV VGVNIAGRSE PATLAQPVTI REIAEPPKIR 841
NP 000247 myosin-binding protein C cardiac-type Homo sapiens EHTSILVKDL PTGARLLFRV RAHNMAGPGA PVTTTEPVTV QEILQRPRLQ 974
NP 004988 myosin-binding protein H Homo sapiens MVTQQTVRNL ALGDKFLLRV SAVSSAGAGP PAMLDQPIHI RENIEAPKIR 175
NP 001010985 myosin-binding protein H-like Homo sapiens ---------- ---------- -------GAGS PT----PQLL PPIEEHPKIW 54
NP 001094228 myosin-binding protein C slow-type Rattus norvegicus DKTKFTITGL PTDAKIFVRV KAINAAGASE PKYYSQPILV KEIIEPPKIR 866
NP 001381991 myosin-binding protein C fast-type Rattus norvegicus ERCGFTVKDL PTGARILFRV VGVNIAGRSE PATLLQPVTI REIVEQPKIR 838
NP 001099960 myosin-binding protein C cardiac-type Rattus norvegicus ERTSLLVKDL PTGARLLFRV RAHNVAGPGG PIITKEPVTV QEILQRPRLQ 974
NP 114001 myosin-binding protein H Rattus norvegicus MVTQQTVRNL ALGDKFFLRV TAVNSAGAGP PAVLDQPVHI QEITEAPKIR 182
NP 001014064 myosin-binding protein H-like Rattus norvegicus ---------- ---------- -TTRQQEAGS PSLQ----LL PSIEEHPKIW 55
NP 001007323 myosin-binding protein C slow-type Danio rerio KKTKYTITGL PVEAKILVRV KAINAAGASQ PRTTQHSILV KEVIEPPKIR 836
NP 001039324 myosin binding protein C2a Danio rerio EKNSFRVRGL PVGEKMLFRV TAVNRAGLSP WAVLAQAVTI REIVESPKIR 813
NP 001013529 myosin binding protein C2b Danio rerio ERQSFVVRNL PTGEKMNFRV VAVNIAGRSP PALLGQPVTV REIMEHPKIR 806
NP 001037814 myosin-binding protein C cardiac-type Danio rerio ERTSVIIRDL PTGEKMQFRV RAYNMAGPSA PATLQQAVTI REIMQRPKIW 957
NP 956852 myosin binding protein Ha Danio rerio . TSNRHAIRGL TTGDMLNIRV VAVNAGGRSE PVVLPQPIPV REVMDRPRIR 208
NP 001093607 myosin binding protein Hb Danio rerio VSCRYVIKNQ TTGDRLNIRV VAMNPGGRSP PATLADTVLV REVVDRPKIR 222
Consensus ERTSFTIRGL PTGAKXLFRV KAVNAAGASX PATLAQPVTV REIIERPKIR
100%
Conservation
0%
Region used for tree in Figure 1

```
                                                                                   1,060                1,080                1,100
XP 046765984 myosin-binding protein C slow-type Gallus gallus               LPRHLKQTYT RRVGETVNLV IPFQGRPRAK VSWQKNGSPI DKNQINIRNT 939
NP 001038124 myosin-binding protein C fast-type Gallus gallus               LPRHLRQTYI RRVGEQVNLV IPFQGKPRPQ VTWSREGGAL -PAEVQTRTS 882
NP 001384345 myosin-binding protein C cardiac-type Gallus gallus            LPRHLRQTLV KKVGETINIM IPFQGKPRPK ISWMKDGQTL DSKDVGIRNS 1022
NP 001026199 myosin-binding protein H Gallus gallus                         LPRQLRQVYV RHVGEAVNLL IPFQGKPQPQ VTWTKDNQPL DTSRVNIRNT 289
NP 002456 myosin-binding protein C slow-type Homo sapiens                   IPRHLKQTYI RRVGEAVNLV IPFQGKPRPE LTWKKDGAEI DKNQINIRNS 897
NP 004524 myosin-binding protein C fast-type Homo sapiens                   LPRHLRQTYI RKVGEQLNLV VPFQGKPRPQ VVWTKGGAPL DTSRVHVRTS 891
NP 000247 myosin-binding protein C cardiac-type Homo sapiens                LPRHLRQTIQ KKVGEPVNLL IPFQGKPRPQ VTWTKEGQPL AGEEVSIRNS 1024
NP 004988 myosin-binding protein H Homo sapiens                             VPRHLRQTYI RQVGETVNLQ IPFQGKPKPQ ATWTHNGHAL DSQRVSMRTG 225
NP 001010985 myosin-binding protein H-like Homo sapiens                     LPRALRQTYI RKVGDTVNLL IPFQGKPKPQ AIWTHDGCAL DTRRVSVRNG 104
NP 001094228 myosin-binding protein C slow-type Rattus norvegicus           IPRHLKQTYI RRVGEAVNLV IPFQGKPRPE LTWKKDGAEI DKNQINIRNS 916
NP 001381991 myosin-binding protein C fast-type Rattus norvegicus           LPRHLRQTYI RKVGEALNLV IPFQGKPRPQ VVWTKGGAPL DTSRVNVRTS 888
NP 001099960 myosin-binding protein C cardiac-type Rattus norvegicus        LPRHLRQTIQ KKVGEPVNLL IPFQGKPRPQ VTWTKEGQPL AGEEVSIRNS 1024
NP 114001 myosin-binding protein H Rattus norvegicus                        VPRHLRQTYI RQVGESVNLQ IPFQGKPKPQ ASWTHNGHAL DSQRVNVRSG 232
NP 001014064 myosin-binding protein H-like Rattus norvegicus                LPRALRQTYI RKVGETVNLL IPFQGKPKPQ TIWTHNGSAL DNSRVSVRNG 105
NP 001007323 myosin-binding protein C slow-type Danio rerio                 IPRHLKQTYI RKVGEVVNLV VPFVGKPRPK VTWLKEGQPI DPT-ISIRNS 885
NP 001039324 myosin binding protein C2a Danio rerio                         LPRHLRTKFI RQVGEKVNLV IPFQGKPRPM VTWLKDGAPV DTGLVNIRTS 863
NP 001013529 myosin binding protein C2b Danio rerio                         LPRDLRTKYI RRVGEKINLV IPFQGKPRPI ATWLKDGQPV DEKKVGVRNS 856
NP 001037814 myosin-binding protein C cardiac-type Danio rerio              LPRNLRQTLI KKVGETINLV IPFQGKPRPQ VSWTKNDEPL DPKSVSIRNS 1007
NP 956852 myosin binding protein Ha Danio rerio .                           MPRALRSKYI RHAGEQINLV IPFSGKPKPV ISWTKNGQPL DTKKVNVRNS 258
NP 001093607 myosin binding protein Hb Danio rerio                          LPRFLRQRYV KKVGEKINLV IPFGGKPQPV VSWTKDGQPI DTKKVNIRNS 272
Consensus                                                                   LPRHLRQTYI RKVGETVNLV IPFQGKPRPQ VTWTKDGQPL DTKRVNIRNS
```

Conservation (100% – 0%)

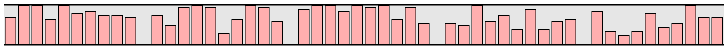

```
                                                                                                        1,120                1,140
XP 046765984 myosin-binding protein C slow-type Gallus gallus               ENDTIIFIRK AERSHSGEYD MKVEVENLVD KATIDIQIVD RPGPPEVVTI 989
NP 001038124 myosin-binding protein C fast-type Gallus gallus               DVDSVFFIRS AARPLSGNYE MRVRIDNMED CATLRLRVVE RPGPPQAVRV 932
NP 001384345 myosin-binding protein C cardiac-type Gallus gallus            STDTILFIRK AELHHSGAYE VTLQIENMTD TVAITIQIID KPGPPQNIKL 1072
NP 001026199 myosin-binding protein H Gallus gallus                         DKDTIFFIRT AQRSDSGKYQ LSVRINGAED KAILDIRVIE RPGPPQNLKL 339
NP 002456 myosin-binding protein C slow-type Homo sapiens                   ETDTIIFIRK AERSHSGKYD LQVKVDKFVE TASIDIQIID RPGPPQIVKI 947
NP 004524 myosin-binding protein C fast-type Homo sapiens                   DFDTVFFVRQ AARSDSGEYE LSVQIENMKD TATIRIRVVE KAGPPINVMV 941
NP 000247 myosin-binding protein C cardiac-type Homo sapiens                PTDTILFIRA ARRVHSGTYQ VTVRIENMED KATLVLQVVD KPSPPQDLRV 1074
NP 004988 myosin-binding protein H Homo sapiens                             DQDSILFIRS AQRSDSGRYE LTVRVEDLEA KAVIDILVIE KPGPPSSIRL 275
NP 001010985 myosin-binding protein H-like Homo sapiens                     EQDSILFIRE AQRADSGRYQ LRVQLGGLEA TATIDILVIE RPGPPQSIKL 154
NP 001094228 myosin-binding protein C slow-type Rattus norvegicus           ETDTIIFIRK AERSHSGKYD LEVKVDKYVE NASIDIQIVD RPGPPQAVTI 966
NP 001381991 myosin-binding protein C fast-type Rattus norvegicus           DFDTVFFVRQ AARSDSGEYE LSVQIENMKD TATIRIRVVE KAGPAENVMV 938
NP 001099960 myosin-binding protein C cardiac-type Rattus norvegicus        PTDTILFIRA AHRTHSGTYQ VTVRIENMED KATLVLQIVD KPSPPLDIRV 1074
NP 114001 myosin-binding protein H Rattus norvegicus                        DQDSILFIRS AQRSDSGRYE LTVRLEGLEA KAAIDILVIE KPGPPSSIKL 282
NP 001014064 myosin-binding protein H-like Rattus norvegicus                EHDSILFIRE AQRTDSGCYQ LCVQLGGLQA TATINILVIE KPGPPQSIKL 155
NP 001007323 myosin-binding protein C slow-type Danio rerio                 DCDSIMFIRK AERKHSGKYE LSVQVENHID TAMLDIQIVD LPGPPQAVKI 935
NP 001039324 myosin binding protein C2a Danio rerio                         DKDTILFIRQ SAREHSGLYT LCVQIENMKD KANLEIQIVD KPGPPMAVTV 913
NP 001013529 myosin binding protein C2b Danio rerio                         NVDSILFIRA AERDHSGKYT LTLKIENMED SATIEIRIVD KPGPPIGVHV 906
NP 001037814 myosin-binding protein C cardiac-type Danio rerio              DTDTILFIRR SERKDSGKYD VQVQIENVED KASVNIQIVD VPGPPQNLKV 1057
NP 956852 myosin binding protein Ha Danio rerio .                           DKDSILFIRK AERDDSGSYE MSVKVDSFED KTTLIIQIVD LPGPPVSVKL 308
NP 001093607 myosin binding protein Hb Danio rerio                          DKDSIFFIRA AERVDSGVYE MCVKVDSFED KAQLTLQIIE LPGPPASVKI 322
Consensus                                                                   DTDTILFIRK AERSDSGKYE LTVQIENMED KATIDIQIVD KPGPPQSVKV
```

Conservation (100% – 0%)

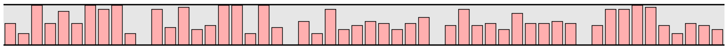

```
                                                                                   1,160                1,180                1,200
XP 046765984 myosin-binding protein C slow-type Gallus gallus               EDVWGENVNL SWKPPKDDGN AAITGYTIQK ADKKSMEWFT VIEHYHRTSA 1039
NP 001038124 myosin-binding protein C fast-type Gallus gallus               MEVWGSNALL QWEPPKDDGN AEISGYTVQK ADTRTMEWFT VLEHSRPTRC 982
NP 001384345 myosin-binding protein C cardiac-type Gallus gallus            ADVWGFNVAL EWTPPQDDGN AQILGYTVQK ADKKTMEWYT VYDHYRRTNC 1122
NP 001026199 myosin-binding protein H Gallus gallus                         VDVWGFNVAL EWSPPADNGN SEIKGYTVQK SDKKSGKWFT VLERCTRTSC 389
NP 002456 myosin-binding protein C slow-type Homo sapiens                   EDVWGENVAL TWTPPKDDGN AAITGYTIQK ADKKSMEWFT VIEHYHRTSA 997
NP 004524 myosin-binding protein C fast-type Homo sapiens                   KEVWGTNALV EWQAPKDDGN SEIMGYFVQK ADKKTMEWFN VYERNRHTSC 991
NP 000247 myosin-binding protein C cardiac-type Homo sapiens                TDAWGLNVAL EWKPPQDVGN TELWGYTVQK ADKKTMEWFT VLEHYRRTHC 1124
NP 004988 myosin-binding protein H Homo sapiens                             LDVWGCNAAL QWTPPQDTGN TELLGYMVQK ADKKTGQWFT VLERYHPTTC 325
NP 001010985 myosin-binding protein H-like Homo sapiens                     VDVWGFSATL EWTPPQDTGN TALLGYTVQK ADTKSGLWFT VLEHYHRTSC 204
NP 001094228 myosin-binding protein C slow-type Rattus norvegicus           EDVWGENVAL TWTPPKDDGN AAITGYTIQK ADKKSMEWFT VIEHYHRTNA 1016
NP 001381991 myosin-binding protein C fast-type Rattus norvegicus           KEVWGTNALV EWQPPKDDGN SEITGYFVQK ADKKTMEWFN VYEHNRHTSC 988
NP 001099960 myosin-binding protein C cardiac-type Rattus norvegicus        VETWGFSVAL EWKPPQDDGN TEIWGYTVQK ADKKTMEWFT VLEHYRQTHC 1124
NP 114001 myosin-binding protein H Rattus norvegicus                        LDVWGCNAAL EWMPPQDTGN TELLGYTVQK ADKKTGQWFT VLERYHPTTC 332
NP 001014064 myosin-binding protein H-like Rattus norvegicus                VDVWGANATL EWTPPQDTGN TALLGYTVQK ADKKSGLWFT VLERYHRTSC 205
NP 001007323 myosin-binding protein C slow-type Danio rerio                 EEIWGNNVAL DWSPPKDSGN APITGYTIQK ADKKTMEWYT CVEHYHRTCI 985
NP 001039324 myosin binding protein C2a Danio rerio                         TDVWGFNAAL EWKPPKDNGN TDIIGYTVQK ADKKTKEWFT VLEHNRRPSC 963
NP 001013529 myosin binding protein C2b Danio rerio                         TDVWGFNAAL EWKPPKDDGN CEITGYTIQK AEKKTKEWFT VYEHNRRTNC 956
NP 001037814 myosin-binding protein C cardiac-type Danio rerio              MDVWGFNVAL EWKPPKDNGN CDITGYNIQK ADKKTMEWYT VFEQYRRTNC 1107
NP 956852 myosin binding protein Ha Danio rerio .                           VDSWGFNAAL EWTPPKDNGN TEITGYTVQK ADKKTGEWFT VLEHFHRLNV 358
NP 001093607 myosin binding protein Hb Danio rerio                          VDTWGFNVAL EWTEPKDNGN TTITGYTIQK ADKKTGDWFT VLEHYHRLNA 372
Consensus                                                                   VDVWGFNXAL EWTPPKDDGN TEITGYTVQK ADKKTMEWFT VLEHYHRTSC
```

Conservation (100% – 0%)

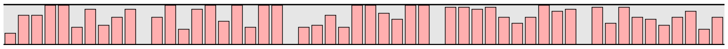

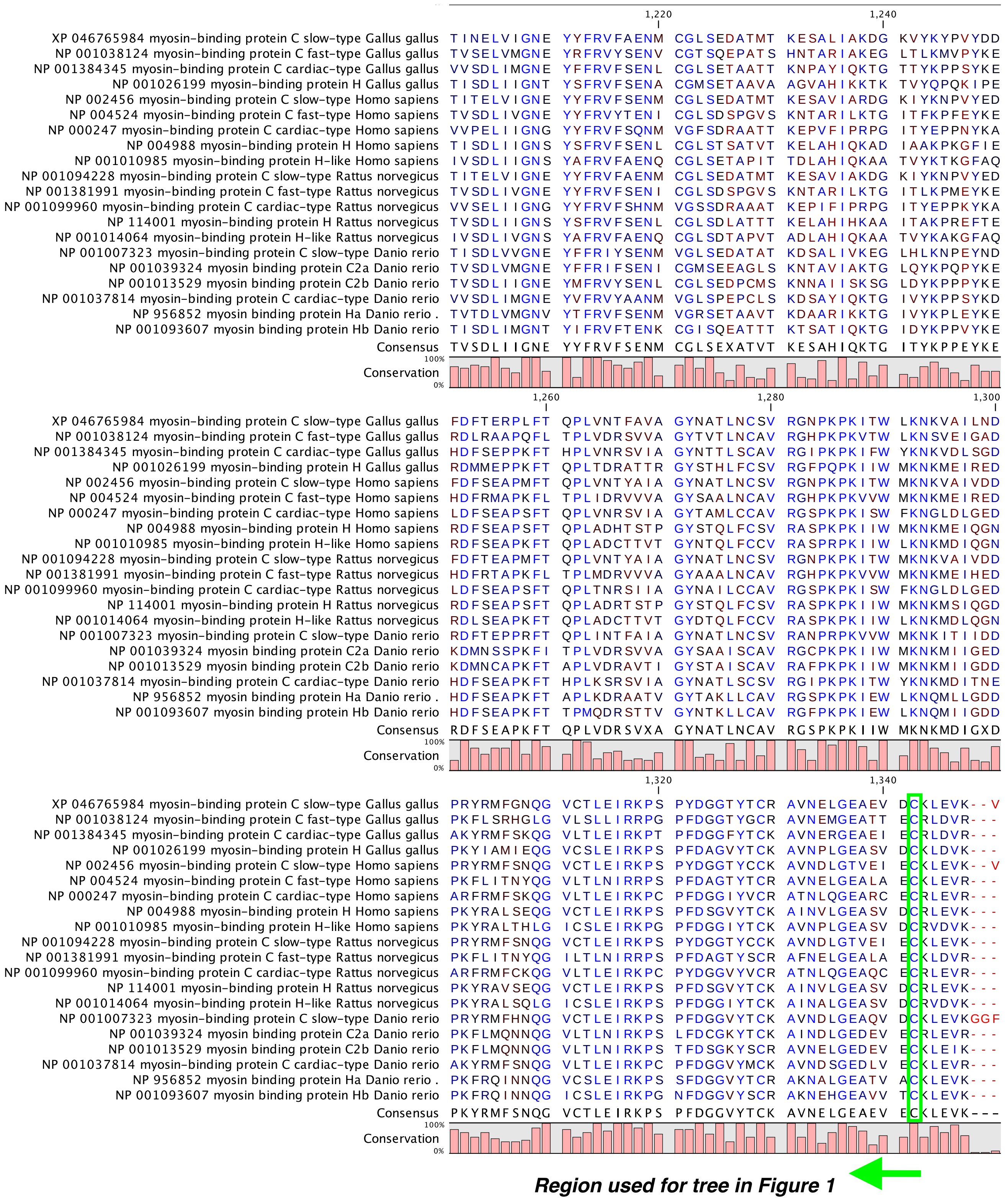
1,220
1,240
XP 046765984 myosin-binding protein C slow-type Gallus gallus TINELVIGNE YYFRVFAENM CGLSEDATMT KESALIAKDG KVYKYPVYDD 1089
NP 001038124 myosin-binding protein C fast-type Gallus gallus TVSELVMGNE YRFRVYSENV CGTSQEPATS HNTARIAKEG LTLKMVPYKE 1032
NP 001384345 myosin-binding protein C cardiac-type Gallus gallus VVSDLIMGNE YFFRVFSENL CGLSETAATT KNPAYIQKTG TTYKPPSYKE 1172
NP 001026199 myosin-binding protein H Gallus gallus TISDLIIGNT YSFRVFSENA CGMSETAAVA AGVAHIKKTK TVYQPQKIPE 439
NP 002456 myosin-binding protein C slow-type Homo sapiens TITELVIGNE YYFRVFSENM CGLSEDATMT KESAVIARDG KIYKNPVYED 1047
NP 004524 myosin-binding protein C fast-type Homo sapiens TVSDLIVGNE YYFRVYTENI CGLSDSPGVS KNTARILKTG ITFKPFEYKE 1041
NP 000247 myosin-binding protein C cardiac-type Homo sapiens VVPELIIGNG YYFRVFSQNM VGFSDRAATT KEPVFIPRPG ITYEPPNYKA 1174
NP 004988 myosin-binding protein H Homo sapiens TISDLIIGNS YSFRVFSENL CGLSTSATVT KELAHIQKAD IAAKPKGFIE 375
NP 001010985 myosin-binding protein H-like Homo sapiens IVSDLIIGNS YAFRVFAENQ CGLSETAPIT TDLAHIQKAA TVYKTKGFAQ 254
NP 001094228 myosin-binding protein C slow-type Rattus norvegicus TITELVIGNE YYFRVFAENM CGLSEDATMT KESAVIAKDG KIYKNPVYED 1066
NP 001381991 myosin-binding protein C fast-type Rattus norvegicus TVSDLIVGNE YYFRVFSENI CGLSDSPGVS KNTARILKTG ITLKPMEYKE 1038
NP 001099960 myosin-binding protein C cardiac-type Rattus norvegicus VVSELIIGNG YYFRVFSHNM VGSSDRAAAT KEPIFIPRPG ITYEPPKYKA 1174
NP 114001 myosin-binding protein H Rattus norvegicus TVSDLIIGNS YSFRVFSENL CGLSDLATTT KELAHIHKAA ITAKPREFTE 382
NP 001014064 myosin-binding protein H-like Rattus norvegicus IVSDLIVGNS YAFRVFAENQ CGLSDTAPVT ADLAHIQKAA TVYKAKGFAQ 255
NP 001007323 myosin-binding protein C slow-type Danio rerio TISDLVVGNE YFFRIYSENM VGLSEDATAT KDSALIVKEG LHLKNPEYND 1035
NP 001039324 myosin binding protein C2a Danio rerio TVSDLVMGNE YFFRIFSENI CGMSEEAGLS KNTAVIAKTG LQYKPQPYKE 1013
NP 001013529 myosin binding protein C2b Danio rerio TVSDLIMGNE YMFRVYSENL CGLSEDPCMS KNNAIISKSG LDYKPPPYKE 1006
NP 001037814 myosin-binding protein C cardiac-type Danio rerio VVSDLIMGNE YVFRVYAANM VGLSPEPCLS KDSAYIQKTG IVYKPPSYKD 1157
NP 956852 myosin binding protein Ha Danio rerio . TVTDLVMGNV YTFRVFSENM VGRSETAAVT KDAARIKKTG IVYKPLEYKE 408
NP 001093607 myosin binding protein Hb Danio rerio TISDLIMGNT YIFRVFTENK CGISQEATTT KTSATIQKTG IDYKPPVYKE 422
Consensus TVSDLIIGNE YYFRVFSENM CGLSEXATVT KESAHIQKTG ITYKPPEYKE
100%
Conservation
0%
1,260
1,280
1,300
XP 046765984 myosin-binding protein C slow-type Gallus gallus FDFTERPLFT QPLVNTFAVA GYNATLNCSV RGNPKPKITW LKNKVAILND 1139
NP 001038124 myosin-binding protein C fast-type Gallus gallus RDLRAAPQFL TPLVDRSVVA GYTVTLNCAV RGHPKPKVTW LKNSVEIGAD 1082
NP 001384345 myosin-binding protein C cardiac-type Gallus gallus HDFSEPPKFT HPLVNRSVIA GYNTTLSCAV RGIPKPKIFW YKNKVDLSGD 1222
NP 001026199 myosin-binding protein H Gallus gallus RDMMEPPKFT QPLTDRATTR GYSTHLFCSV RGFPQPKIIW MKNKMEIRED 489
NP 002456 myosin-binding protein C slow-type Homo sapiens FDFSEAPMFT QPLVNTYAIA GYNATLNCSV RGNPKPKITW MKNKVAIVDD 1097
NP 004524 myosin-binding protein C fast-type Homo sapiens HDFRMAPKFL TPLIDRVVVA GYSAALNCAV RGHPKPKVVW MKNKMEIRED 1091
NP 000247 myosin-binding protein C cardiac-type Homo sapiens LDFSEAPSFT QPLVNRSVIA GYTAMLCCAV RGSPKPKISW FKNGLDLGED 1224
NP 004988 myosin-binding protein H Homo sapiens RDFSEAPSFT QPLADHTSTP GYSTQLFCSV RASPKPKIIW MKNKMEIQGN 425
NP 001010985 myosin-binding protein H-like Homo sapiens RDFSEAPKFT QPLADCTTVT GYNTQLFCCV RASPRPKIIW LKNKMDIQGN 304
NP 001094228 myosin-binding protein C slow-type Rattus norvegicus FDFTEAPMFT QPLVNTYAIA GYNATLNCSV RGNPKPKITW MKNKVAIVDD 1116
NP 001381991 myosin-binding protein C fast-type Rattus norvegicus HDFRTAPKFL TPLMDRVVVA GYAAALNCAV RGHPKPKVVW MKNKMEIHED 1088
NP 001099960 myosin-binding protein C cardiac-type Rattus norvegicus LDFSEAPSFT QPLTNRSIIA GYNAILCCAV RGSPKPKISW FKNGLDLGED 1224
NP 114001 myosin-binding protein H Rattus norvegicus RDFSEAPSFT QPLADRTSTP GYSTQLFCSV RASPKPKIIW MKNKMSIQGD 432
NP 001014064 myosin-binding protein H-like Rattus norvegicus RDLSEAPKFT QPLADCTTVT GYDTQLFCCV RASPKPKIIW LKNKMDLQGN 305
NP 001007323 myosin-binding protein C slow-type Danio rerio RDFTEPPRFT QPLINTFAIA GYNATLNCSV RANPRPKVVW MKNKITIIDD 1085
NP 001039324 myosin binding protein C2a Danio rerio KDMNSSPKFI TPLVDRSVVA GYSAAISCAV RGCPKPKIIW MKNKMIIGED 1063
NP 001013529 myosin binding protein C2b Danio rerio KDMNCAPKFT APLVDRAVTI GYSTAISCAV RAFPKPKIIW MKNKMIIGDD 1056
NP 001037814 myosin-binding protein C cardiac-type Danio rerio HDFSEAPKFT HPLKSRSVIA GYNATLSCSV RGIPKPKITW YKNKMDITNE 1207
NP 956852 myosin binding protein Ha Danio rerio . HDFSEAPKFT APLKDRAATV GYTAKLLCAV RGSPKPKIEW LKNQMLLGDD 458
NP 001093607 myosin binding protein Hb Danio rerio HDFSEAPKFT TPMQDRSTTV GYNTKLLCAV RGFPKPKIEW LKNQMIIGDD 472
Consensus RDFSEAPKFT QPLVDRSVXA GYNATLNCAV RGSPKPKIIW MKNKMDIGXD
100%
Conservation
0%
1,320
1,340
XP 046765984 myosin-binding protein C slow-type Gallus gallus PRYRMFGNQG VCTLEIRKPS PYDGGTYTCR AVNELGEAEV DCKLEVK--V 1187
NP 001038124 myosin-binding protein C fast-type Gallus gallus PKFLSRHGLG VLSLLIRRPG PFDGGTYGCR AVNEMGEATT ECRLDVR--- 1129
NP 001384345 myosin-binding protein C cardiac-type Gallus gallus AKYRMFSKQG VLTLEIRKPT PFDGGFYTCK AVNERGEAEI ECRLDVR--- 1269
NP 001026199 myosin-binding protein H Gallus gallus PKYIAMIEQG VCSLEIRKPS PFDAGVYTCK AVNPLGEASV DCKLDVK--- 536
NP 002456 myosin-binding protein C slow-type Homo sapiens PRYRMFSNQG VCTLEIRKPS PYDGGTYCCK AVNDLGTVEI ECKLEVK--V 1145
NP 004524 myosin-binding protein C fast-type Homo sapiens PKFLITNYQG VLTLNIRRPS PFDAGTYTCR AVNELGEALA ECKLEVR--- 1138
NP 000247 myosin-binding protein C cardiac-type Homo sapiens ARFRMFSKQG VLTLEIRKPC PFDGGIYVCR ATNLQGEARC ECRLEVR--- 1271
NP 004988 myosin-binding protein H Homo sapiens PKYRALSEQG VCTLEIRKPS PFDSGVYTCK AINVLGEASV DCRLEVK--- 472
NP 001010985 myosin-binding protein H-like Homo sapiens PKYRALTHLG ICSLEIRKPG PFDGGIYTCK AVNPLGEASV DCRVDVK--- 351
NP 001094228 myosin-binding protein C slow-type Rattus norvegicus PRYRMFSNQG VCTLEIRKPS PYDGGTYCCK AVNDLGTVEI ECKLEVK--- 1163
NP 001381991 myosin-binding protein C fast-type Rattus norvegicus PKFLITNYQG ILTLNIRRPS PFDAGTYSCR AFNELGEALA ECKLDVR--- 1135
NP 001099960 myosin-binding protein C cardiac-type Rattus norvegicus ARFRMFCKQG VLTLEIRKPC PYDGGVYVCR ATNLQGEAQC ECRLEVR--- 1271
NP 114001 myosin-binding protein H Rattus norvegicus PKYRAVSEQG VCTLEIRKPS PFDSGVYTCK AINVLGEASV DCRLEVK--- 479
NP 001014064 myosin-binding protein H-like Rattus norvegicus PKYRALSQLG ICSLEIRKPS PFDGGIYTCK AINALGEASV DCRVDVK--- 352
NP 001007323 myosin-binding protein C slow-type Danio rerio PRYRMFHNQG VCTLEIRKPS PFDGGVYTCK AVNDLGEAQV DCKLEVKGGF 1135
NP 001039324 myosin binding protein C2a Danio rerio PKFLMQNNQG VLTLNIRKPS LFDCGKYTCK AINDLGEDEV ECRLEVR--- 1110
NP 001013529 myosin binding protein C2b Danio rerio PKFLMQNNQG VLTLNIRKPS TFDSGKYSCR AVNELGEDEV ECKLEIK--- 1103
NP 001037814 myosin-binding protein C cardiac-type Danio rerio AKYRMFSNQG VLTLEIRKPC PFDGGVYMCK AVNDSGEDLV ECKLEVR--- 1254
NP 956852 myosin binding protein Ha Danio rerio . PKFRQINNQG VCSLEIRKPS SFDGGVYTCR AKNALGEATV ACKLEVK--- 505
NP 001093607 myosin binding protein Hb Danio rerio PKFRQINNQG ICSLEIRKPG NFDGGVYSCR AKNEHGEAVV TCKLEVK--- 519
Consensus PKYRMFSNQG VCTLEIRKPS PFDGGVYTCK AVNELGEAEV ECKLEVK---
100%
Conservation
0%
Region used for tree in Figure 1

| | 1,360 | | 1,380 | | |
|---|---|---|---|---|---|
| XP 046765984 myosin-binding protein C slow-type Gallus gallus | IYQGVINPGE | PI-----FMA | GNKQ-LQNKD | F-- | 1212 |
| NP 001038124 myosin-binding protein C fast-type Gallus gallus | ---------V | PQ-------- | ---------- | --- | 1132 |
| NP 001384345 myosin-binding protein C cardiac-type Gallus gallus | ---------V | PQ-------- | ---------- | --- | 1272 |
| NP 001026199 myosin-binding protein H Gallus gallus | ---------M | PK-------- | ---------- | --- | 539 |
| NP 002456 myosin-binding protein C slow-type Homo sapiens | IYQGVNTPGQ | PV-----FLE | GQQQSLHNKD | F-- | 1171 |
| NP 004524 myosin-binding protein C fast-type Homo sapiens | ---------V | PQ-------- | ---------- | --- | 1141 |
| NP 000247 myosin-binding protein C cardiac-type Homo sapiens | ---------V | PQ-------- | ---------- | --- | 1274 |
| NP 004988 myosin-binding protein H Homo sapiens | ---------A | SA-------- | -----AH--- | --- | 477 |
| NP 001010985 myosin-binding protein H-like Homo sapiens | ---------V | PN-------- | ---------- | --- | 354 |
| NP 001094228 myosin-binding protein C slow-type Rattus norvegicus | --------VV | AQ-------- | ---------- | --- | 1167 |
| NP 001381991 myosin-binding protein C fast-type Rattus norvegicus | ---------V | PQ-------- | ---------- | --- | 1138 |
| NP 001099960 myosin-binding protein C cardiac-type Rattus norvegicus | ---------V | PQ-------- | ---------- | --- | 1274 |
| NP 114001 myosin-binding protein H Rattus norvegicus | ---------A | SA-------- | -----TH--- | --- | 484 |
| NP 001014064 myosin-binding protein H-like Rattus norvegicus | ---------A | PH-------- | ---------- | --- | 355 |
| NP 001007323 myosin-binding protein C slow-type Danio rerio | TFFELMKRGV | PLHLIDKYMN | ESKTSESEKK | GSD | 1168 |
| NP 001039324 myosin binding protein C2a Danio rerio | ---------V | VT-------- | ------EKEE | AKK | 1120 |
| NP 001013529 myosin binding protein C2b Danio rerio | --------AI | PT-------- | ---------E | KKD | 1111 |
| NP 001037814 myosin-binding protein C cardiac-type Danio rerio | ---------L | PQ-------- | -----VTKED | IKK | 1265 |
| NP 956852 myosin binding protein Ha Danio rerio . | ---------Q | VT-------- | -----IQEEK | DKK | 516 |
| NP 001093607 myosin binding protein Hb Danio rerio | ---------Q | PV-------- | ------IADA | DKK | 529 |
| Consensus | ---------V | PQ-------- | ---------- | --- |  |

Conservation

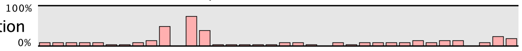
